# Dual Acting Small-Molecule Inhibitors Targeting Mycobacterial DNA Replication

**DOI:** 10.1101/561506

**Authors:** Meenakshi Singh, Stefan Ilic, Benjamin Tam, Yasmin Ben-Ishay, Dror Sherf, Doron Pappo, Barak Akabayov

## Abstract

*Mycobacterium tuberculosis* (*Mtb*) is a pathogenic bacterium and a causative agent of tuberculosis (TB), a disease that kills more than 1.5 million people worldwide annually. One of the main reasons for this high mortality rate is the evolution of new *Mtb* strains that are resistant to available antibiotics. Therefore, new therapeutics for TB are in constant demand. Here we report the development of such inhibitors that target two DNA replication enzymes of *Mtb*, namely DnaG primase and DNA gyrase, which share a conserved TOPRIM fold near the inhibitors’ binding site. The molecules were developed on the basis of previously reported inhibitors for T7 DNA primase that bind near the TOPRIM fold. In order to improve the physicochemical properties of the molecules as well as their inhibitory effect on primase and gyrase, 49 novel compounds were synthesized as potential drug candidates in three stages of optimization. The last stage of chemical optimization yielded two novel inhibitors for the fast-growing nonpathogenic model *Mycobacterium smegmatis* (*Msmg*).

## Introduction

Replication of the chromosome is performed by coordinated activity of different enzymes comprising the replisome. Extensive characterization of the components of the *E. coli* replisome has revealed the molecular mechanisms within the replisome ^[1]^. Based on the structural and functional differences between bacterial and human replication apparatus, it was suggested that the bacterial replisome could serve as a novel drug target ^[2]^. During DNA replication, DnaB helicase unwinds double-stranded DNA and exposes two individual strands ^[3]^. One DNA strand is copied continuously (the leading strand) and the other discontinuously (the lagging strand). On the lagging strand, DnaG primase synthesizes short RNA primers that mark the starting points of “Okazaki fragments” synthesis by DNA polymerase III holoenzyme. The process of DNA unwinding by DnaB helicase causes an increase in twisting tension at the coiled portion of DNA as the replication fork advances. This tension interferes with a further unwinding of the double-stranded DNA. Therefore, Gyr, a type II topoisomerase, acts by creating a transient double-stranded DNA breaks to release the tension and ensures that the chromosome is always negatively supercoiled ^[4]^. Both DnaG primase and Gyr are essential for viability of bacterial cells, and have different structural and functional settings in human replisomes (Figure 1A), thereby providing attractive candidates for antibiotic targeting ^[5]^.

**Figure 1.**
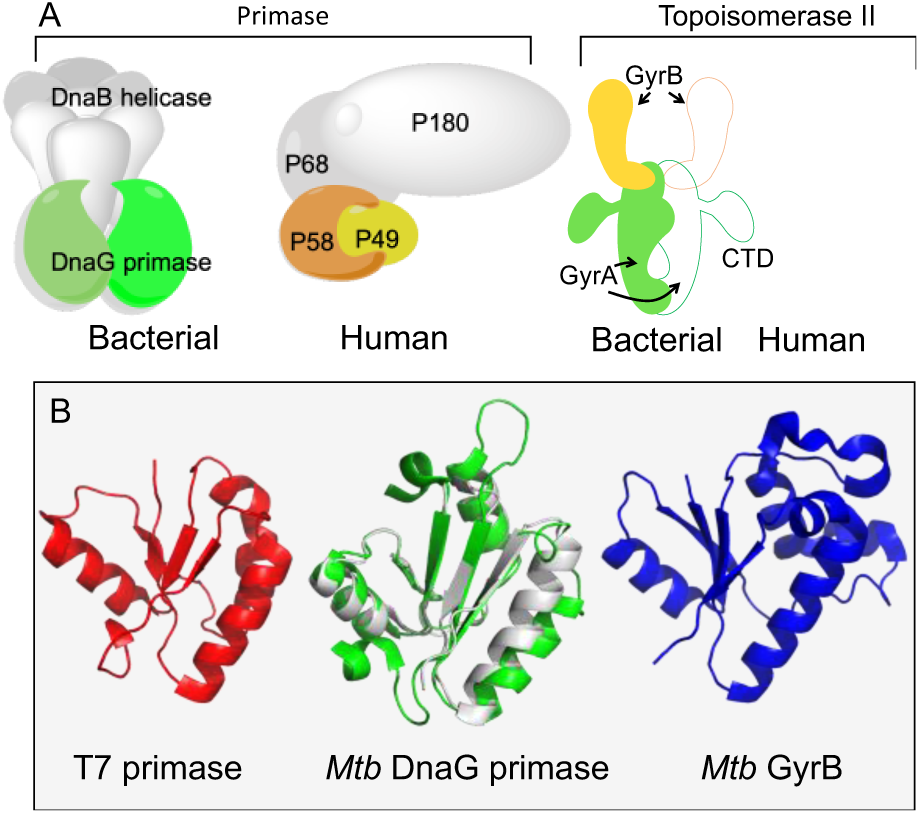
Bacterial targets in this study: DNA primase and gyrase. (A) Schematic representation of DNA priming system and DNA topoisomerase II in bacteria compared to their analogues in humans. (B) Structural homology of the TOPRIM fold in Gyr and DnaG. T7 DNA primase (red, PDBID 1NUI ^[6]^), part of the fused helicase-primase gp4 of bacteriophage T7, shares a similar structure with DnaG from *Mtb* (green, PDBID 5W33 ^[7]^) and *S. aureus* (grey PDBID 4E2K ^[8]^). *Mtb* Gyr (PDBID 3M4I ^[9]^) is presented in blue.

Bacterial DnaG-type primases, as well as type IA and type II topoisomerases (including Gyr), belong to the same enzyme superfamily based on the presence of a conserved TOPRIM fold ^[10]^ (Figure 1B). This fold consists of approximately 100 amino acids and includes two motifs, one containing a very conserved glutamate residue in its center, whereas the other involves two conserved aspartates (DxD motif, Supporting Information Figure S1) ^[10]^. The two conserved motifs have an essential role in both primases (e.g., nucleotide polymerization) ^[11]^ and topoisomerases (e.g. strand cleavage and rejoining, DNA binding, metal coordination) ^[12]^. It is therefore expected that inhibitors that target the TOPRIM fold can cross-interact with different TOPRIM-containing enzymes as in the case of coumarin. Coumarin is an inhibitor that binds to the TOPRIM fold of GyrB ^[13]^, and also inhibits DnaG from *Mtb* ^[14]^.

*Mtb*, whose replication proteins are discussed herein, is a Gram-positive pathogenic bacterium that is the causative agent of tuberculosis (TB) which infects a third of the world’s population. New strains of *Mtb*, that are resistant to all known classes of antibiotics, pose a global healthcare threat ^[15]^. DNA replication is an underexploited target for antibiotics but has the potential to yield novel inhibitors for *Mtb* ^[2]^. Proteins of high potential to become effective drug targets are those that present marked structural difference compared to the human analogs, such as DnaG primase and DNA gyrase.

Unlike DnaG, which serves as a novel drug target, Gyr has already been exploited as a clinical antitubercular target. For example, fluoroquinolones that stabilize the DNA-gyrase complex and promote genomic DNA cleavage ^[16]^, along with Gyr inhibitors, such as novobiocin, represent antibacterial agents with proven therapeutic efficiency ^[17]^. Also, it is important to note that the resistance to fluoroquinolones (e.g., moxifloxacin and gatifloxacin) remains uncommon in clinical isolates of *Mtb* ^[18]^.

We have previously reported a method that combines fragment-based screening by NMR with *in-silico* optimization steps to identify inhibitors for DnaG-type DNA primase ^[19]^. Fragment-based screening is advantageous over commonly used high-throughput screening for slow enzymes such as DNA primase, where the readout of biochemical activity is extremely low ^[5, 20]^. Subset of drug-sized molecules, that contained same scaffold as the initial fragment molecules, were selected from the Zinc database ^[21]^ and docked into the crystal structure of T7 primase. The molecules with the lowest predicted binding energy were used for further biochemical and structural studies. This method, designated as fragment-based virtual screening ^[19b]^, yielded five small molecule inhibitors that target T7 DNA primase through common binding mechanism verified by high resolution NMR analysis ^[19b]^.

Herein, we describe the development of novel small molecule inhibitors, which target *Mtb* DnaG, based on the molecules that were found to inhibit T7 DNA primase. Some of these molecules also inhibit Gyr, presumably through binding near the TOPRIM fold. Two of the molecules that inhibit both *Mtb* DnaG and Gyr were found to inhibit growth of *Msmg*, thus offering hope for obtaining novel drugs against TB.

## Results

### Small molecules that inhibit T7 DNA primase also inhibit *Mtb* DnaG

The primase domain of gene 4 protein of bacteriophage T7 (T7 DNA primase) was used as a model target for the development of anti-DnaG inhibitors ^[5]^. By using NMR-fragment based screening combined with computational optimization, we found five small-molecules that inhibit T7 DNA primase ^[19b]^. The binding mechanism of three inhibitors (one containing 2H-chromene and two containing indole) was revealed by high field NMR spectroscopy ^[19b]^, whereas three amino acids (Val101, Met105, Tyr106) were found to mediate the binding of all three molecules. Therefore, we tested the ability of these molecules, designed initially to inhibit T7 DNA primase, to impair RNA primers synthesis by *Mtb* DnaG due to the high structural similarity of these proteins (details on the initial small-molecules are presented in Supporting Information Table 1). Structural and sequence similarity within the TOPRIM superfamily of enzymes as well as the close proximity of the initial molecules’ binding site to the TOPRIM fold of T7 primase, led us to hypothesize that other TOPRIM containing enzymes, such as *Mtb* DnaG primase and Gyr could also be targets for these inhibitors (Supporting Information Figure S2-S3, the active site residues and the conserved active site motifs are indicated). We also performed a docking analysis (Supporting Information Figure S4) and confirmed, based on the negative ΔG values, that the molecules may indeed bind to this region.

The reaction involved incubating the *Mtb* DnaG with an oligonucleotide template (5’-CCGACCCGTCCGTAATACAGAGGTAATTGTCACGGT - 3’), each one of the four rNTPs, and [α-^32^P]-ATP. The radioactively labeled RNA products were separated by denaturing gel analysis and visualized on an autoradiogram (Figure 2A). The decrease in the radioactive signal strength, which is directly proportional to the number of synthesized RNA primers, indicates the inhibition of primase activity. Remarkably, two of the five molecules that inhibit T7 DNA primase, 3-(2-(ethoxycarbonyl)-5-nitro-1H-indol-3-yl)propanoic acid (Compound **A13**) and 7-nitro-1H-indole-2-carboxylic acid (Compound **A17**), were also found to inhibit the RNA primer synthesis of *Mtb* DnaG (Figure 2A, lanes 13 and 17).

**Figure 2.**
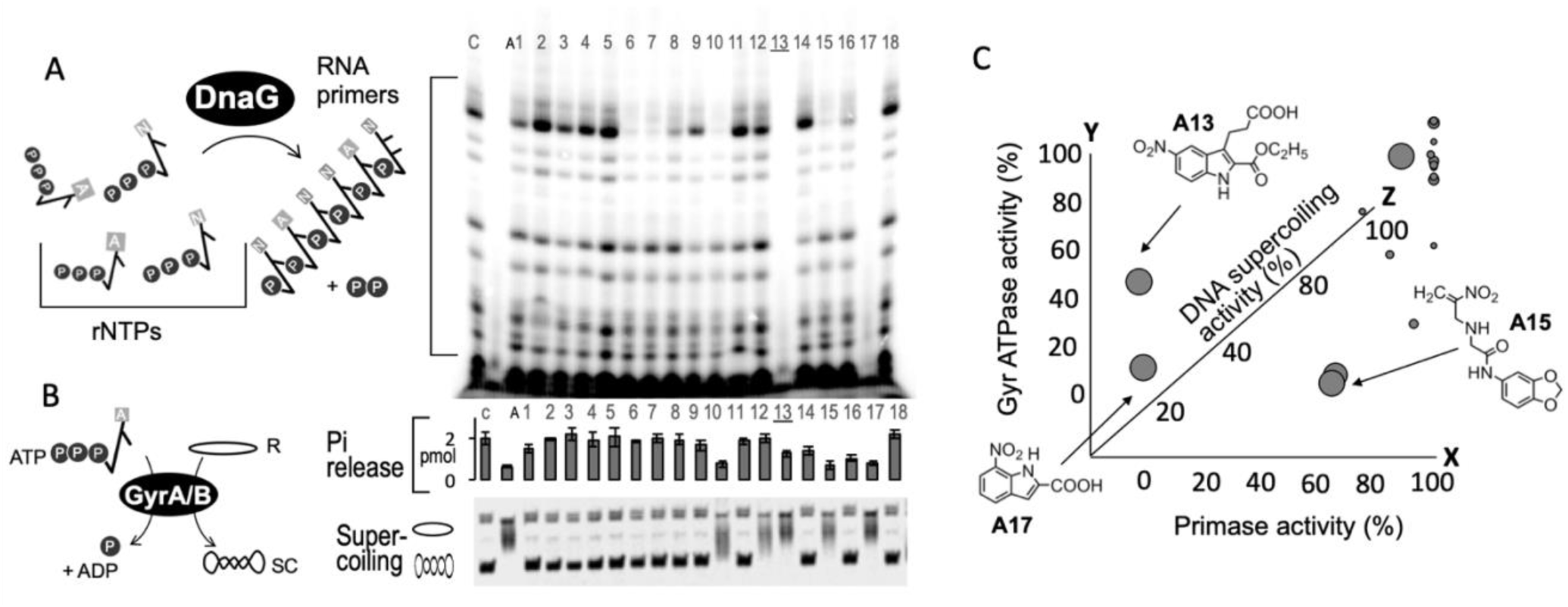
Effect of selected small molecules on the activities of DnaG primase and gyrase from *Mtb*. Screening for best precursor molecule among 18 commercially available compounds by examining their inhibitory effect on specific *Mtb* DnaG primase (template-directed ribonucleotide synthesis) and gyrase activity (ATPase activity and DNA-supercoiling). (A) General scheme of the template-directed ribonucleotide synthesis catalyzed by the primase. *Mtb* DnaG (500 nM) was incubated in the presence of 50 μM ssDNA template, 250 μM GTP, CTP, UTP, α-^32^P-ATP and 4 mM of the selected inhibitor. Radioactive RNA-products at the end of the reaction were separated by gel-electrophoresis and visualized by autoradiography. Gel bands represent RNA primers formed by *Mtb* DnaG. Lower intensity of the bands indicates *Mtb* DnaG inhibition. (B) General scheme showing the enzymatic activity of the Gyrase – catalysis of negative supercoiling of a plasmid by using the energy obtained by hydrolyzing two ATP molecules. Top: Inhibition of ATP hydrolysis by small molecule inhibitors. The reaction contained 36nM DNA GyrA, 18nM GyrB, 13.3 ng/µl pBR322 DNA and 10 μM γ-^32^P-ATP in the presence of 500 μM each small molecule. The reaction was stopped by adding Norit solution (*see* Online Materials and Methods) and the radioactivity of the supernatant was measured. Bottom: Inhibition of DNA supercoil ing by small molecule inhibitors. DNA supercoiling assay was carried out under the same reaction conditions as the ATPase assay except that 1mM non-radioactive ATP was used instead of 10 µM [γ-32P]-ATP 100 nCi. The Reactions stopped, treated by chloroform/isoamyl alcohol (v:v, 24:1) and loaded onto 1% TBE X1 agarose gel. (C) Distribution map illustrating the effect of each inhibitor on primase/Gyr activities. Each axis represents the specific activity (in percent) of the enzymes in the presence of a given inhibitor, relative to the control (enzymatic activity without any inhibitor); X-axis: primase activity, y-axis: Gyr ATPase activity, z-axis: Gyr supercoiling activity (circle size is proportional to magnitude of inhibition). Values used to create this panel can be found in Additional Materials.

**Scheme 1.**
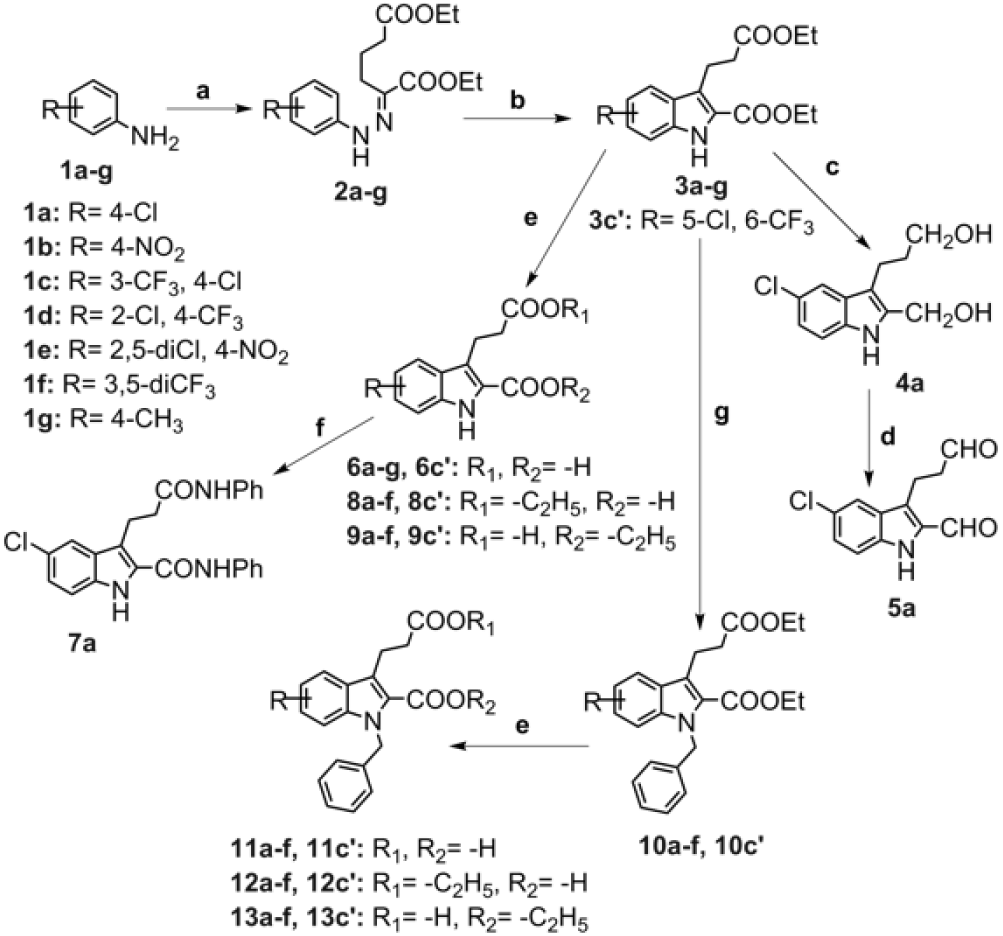
Synthetic scheme of small-molecule inhibitors. Reagents and conditions: a. i. HCl, NaNO_2_, H_2_O, 0°C, 1h; ii. ethyl 2-oxocyclopentane carboxylate, KOH (4 M), 0°C; 1 h; iii. H_2_SO_4_, ethanol, 100°C, 3 h; b. *p*-toluene sulfonic acid (PTSA), benzene, 110°C, 24 h; c. LiAlH_4_, THF, H_2_SO_4_, rt, 2 h; d. pyridinium chlorochromate (PCC), DCM, rt, 2-3 h; e. NaOH (0.01 M)/THF, rt, 18-24 h; f. DCC, HOBt, dry DMF, aniline, rt, 24h; g. benzyl bromide, Cs_2_CO_3_, KI, DMF, 60°C, 2 h.

### Few inhibitors for *Mtb* DnaG cross-interact with Gyr of *Mtb* (*Mtb* Gyr)

The compounds found to inhibit DnaG primase were also tested against *Mtb* Gyr, since both enzymes share a conserved TOPRIM fold. The original inhibitors, from which the current molecules were developed, were bound near the conserved TOPRIM fold of the T7 primase. To evaluate the inhibition of two activities of *Mtb* Gyr, ATP hydrolysis rate and DNA supercoiling assay were carried out (Figure 2B). Upon incubation of *Mtb* Gyr with DNA and [γ-^32^P]-ATP a radioactive P_i_ is formed and remains free in solution after treatment with active charcoal. In the presence of a small-molecule inhibitor, the decrease in the amount of ^32^P_i_, indicates a decrease in ATP hydrolysis by *Mtb* Gyr (Figure 2B, top panel). The results are presented as percent of activity, relative to the control sample that didn’t contain any inhibitor.

The effect of small molecule inhibitors on the formation of cleavable-complex species by *Mtb* Gyr was also measured. In the presence of ATP and absence of inhibitor the enzyme shows marked negative supercoiling on pBR322 DNA. The inhibition of supercoiling activity is characterized by the appearance of a ladder of gel bands (Figure 2B, bottom panel).

Comparison of all three measured activities of Gyr and primase places the best inhibitor in the bottom left corner of the distribution map (Figure 2C). Some of the molecules (**A13, A15** and **A17**) that were found to reduce [^32^P]_i_ release (ATP hydrolysis) also inhibit the supercoiling of pBR322 DNA by *Mtb* Gyr (Figure 2C, top-left). Two of the molecules (**A13** and **A17**) that were found as best inhibitors for *Mtb* DnaG and Gyr contained the indole scaffold substituted with carboxylic acid/ester in position two/three and electron-withdrawing nitro group in position five/seven. We reasoned that the bulkier structure of molecule **A13** may enable broader network of interactions with the target macromolecule, thus promoting tighter binding and better inhibition.

### Preparation of library of derivatives for the inhibition of *Mtb* DnaG

A library of 49 novel derivatives was synthesized on the basis of molecule **A13** in order to improve inhibition efficacy. The synthesis of ethyl 3-(3-ethoxy-3-oxopropyl)-1H-indole-2-carboxylate derivatives (**3a-g, 3c’, 4a, 5a, 6a-g, 6c’, 7a, 8a-f, 8c’, 9a-f, 9c’**) and the *N*-benzylated derivatives of ethyl 3-(3-ethoxy-3-oxopropyl)-1H-indole-2-carboxylate (**10a-f, 10c’, 11a-f, 11c’, 12a-f, 12c’, 13a-f** and **13c’**) is depicted in Scheme 1 and Supporting Information Figure S5.

The synthesis commenced by Japp-Klingemann condensations of the commercially available aniline derivatives **1a-g** and afforded the corresponding diethyl derivatives of phenylhydrazones (**2a-g**) in 60-80% yields. Subsequent Fischer indole cyclization of the latter phenylhydrazones (p-toluenesulfonic acid, benzene, reflux) ^[22]^, produced the key indole derivatives (**3a-g, 3c’**) in 15–80% yields. Compound 3c and 3c’ are the two regioisomers that are produced after Fischer cyclization in the ratio of 1:1.25 (3c’:3c). These regioisomers were isolated through column chromatography using ethyl acetate/hexane (1:8) as mixture for elution. Hydrolysis of the latter diesters under mild basic conditions (0.01M NaOH) gave a mixture of the corresponding dicarboxylic acid derivatives **6a-g, 6c’** and mono-carboxylic acid derivatives **8a-f, 8c’, 9a-f, 9c’**. Reduction of indole diester **3a** using LiAlH_4_ (THF, 4 °C) produced 3-(5-chloro-2-(hydroxymethyl)-1H-indol-3-yl) propan-1-ol (**4a**) in 50% yield, while selective oxidation of the latter di-alcohol **4a** using PCC afforded dicarboxaldehyde **5a** in 40% yield. Diamide **7a** was prepared from acid **6a** (DCC and HoBt, dry DMF) in 60% yield. Benzylation of compounds **3a-f, 3c’** (BnBr, Cs_2_CO_3_, DMF) afforded obtaining *N*-benzylindoles **10a-f, 10c’** in 85-95% yields ^[23]^. Partial hydrolysis of the latter diesters (**10a-f, 10c’**) under mild basic conditions (0.01M NaOH) led to the formation of the corresponding dicarboxylic and mono-carboxylic acids **11a-f, 11c’, 12a-f, 12c’, 13a-f**, and **13c’** in 10-75% yields.

### SAR studies of newly synthesized indole derivatives

The newly synthesized molecules (Figure 3A) were tested for their ability to inhibit the activities of *Mtb* DnaG and Gyr (Supporting Information Figure S6). Dose response experiments revealed micromolar range for inhibition of all three activities (Supporting Information Figure S7). We have observed that the presence of electron donating groups, especially methyl group (**3g**), resulted in decreased potency of inhibition compared to compound **3c’**. The reduction of ester function (**3a**) to alcohol group (**4a**) and then oxidation to aldehyde group (**5a**) led to loss of activity. The amide coupling to carboxylic acid derivative (**6a**) gave rise to compound (**7a**), which presented some inhibitory activity on Gyr but not on primase. In order to explore the effect of *N*-benzylation of indole on inhibitory activity, compounds (**10a-f, 10c’, 11a-f, 11c’, 12a-f, 12c’, 13a-f** and **13c’**) were synthesized and their efficacies were tested. When the benzyl group was introduced at the NH-position of the indole ring, the inhibitory effects of the resulting compounds increased dramatically in comparison with compounds (**3a-g, 3c’, 6a-g, 6c’, 8a-f, 8c’, 9a-f, 9c’**). Most notably, the *N*-benzylation of compounds (**3a-g, 3c’**) gave rise to compounds (**13a-f** and **13c’**), which presented maximum inhibitory activity thus far.

**Figure 3.**
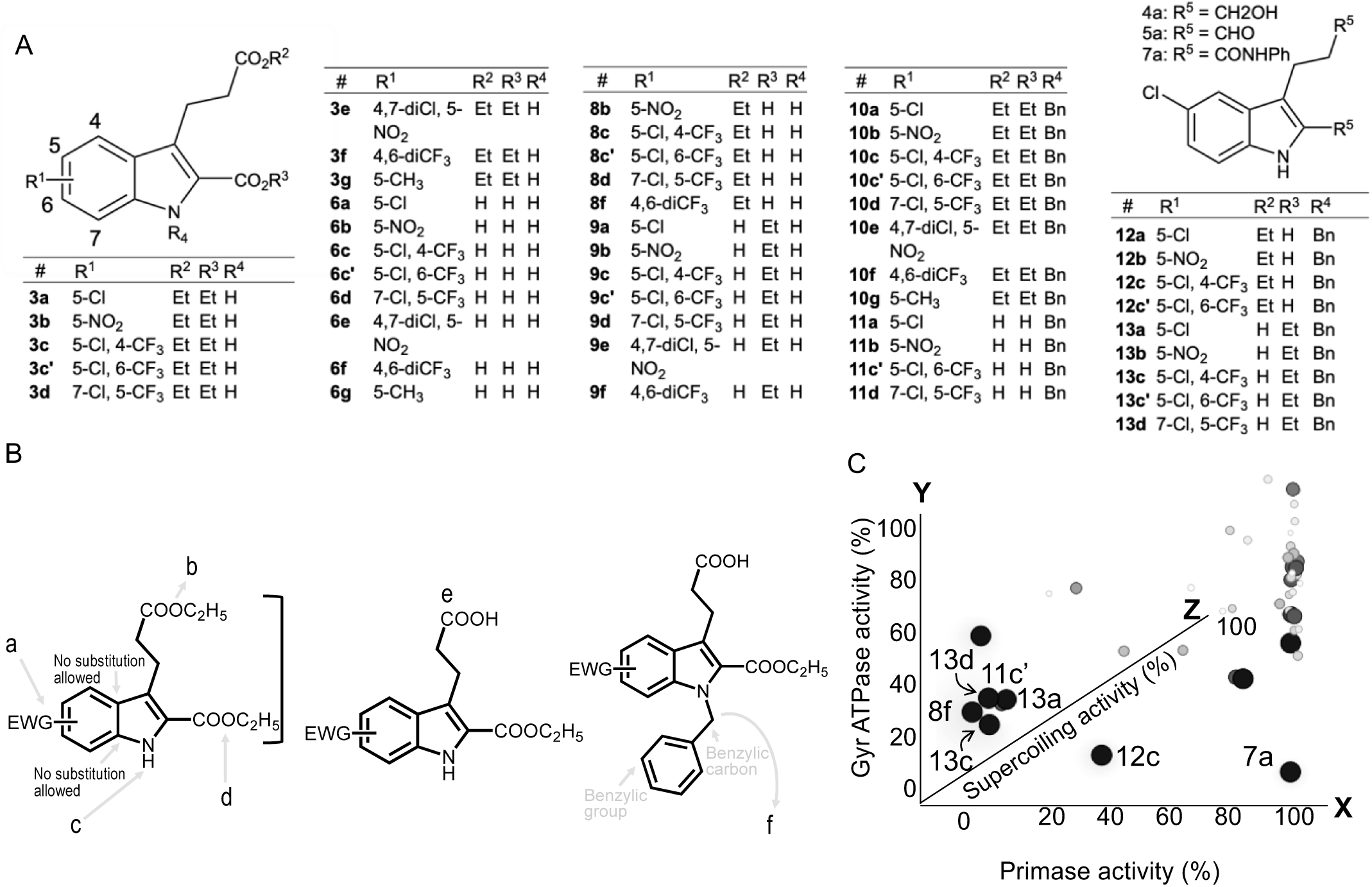
SAR-guided optimization of small molecule inhibitors for DnaG/GyrB of *Mtb*. (A) Optimization of small molecule inhibitors was performed in three rounds of synthesis. Round I included development of indole derivatives (3a – 7a) viz. indole diesters, indole diacids, indole dialcohols, indole dicarboxaldehyde and indole diamides. Round II included development of indole derivatives (8b – 9f) viz. indole monoesters and indole monoacids. Round III included development of indole derivatives (10a – 13d) viz. N-benzylated indole diesters, N-benzylated indole diacids, N-benzylated indole monoester and N-benzylated indole monoacid. (B) Scheme of synthesis of derivative molecules based on SAR conclusions. Group 1) a. Electron withdrawing group (e.g., NO _2_, CF_3_, Cl) render phenyl ring less nucleophilic and support activity. Incorporation of electron donating group (e.g., CH_3_, OH) hinders activity. b. Bulky groups introduce steric hindrance and decrease activity. Five membered nitrogen heterocyclic ring system is essential for activity. c. Free nitrogen is essential for activity. Lone pair located on the nitrogen atom is in resonance, and therefore, it stabilizes the structure. d. Carboxylate ester is essential for activity. Activity is reduced when modified to alcohols, amides, aldehydes but increased upon conversion to acid. Group 2) e. Carboxyl group is essential. Activity is reduced when modified to ester, alcohols, amides, aldehydes. Group 3) f. Utilization of free –NH linkage of Indole. Substituting the –H of –NH of Indole with benzyl group increases the inhibition. (C) Correlation of enzymatic activities in the presence of derivative molecules (results of all three activities as well as the reaction conditions are summarized in Supporting Information Figure S6). X-axis: inhibition of primase activity, y-axis: inhibition of ATPase activity of Gyr, z-axis: inhibition of Gyr supercoiling activity (circle size is proportional to magnitude of inhibition). Molecules that weren’t successfully synthesized or weren’t fully characterized are not included in the functional analysis. Values used to create this panel can be found in Additional Materials.

Importantly, when the ester group was replaced with a carboxylic acid moiety in position 3, the molecules (**13c** and **13d**) were more active both in enzymatic and bacterial assays (Figures 3-4), indicating that the presence of carboxyethyl and propanoic acid group at position 2 and 3, respectively, is essential for an extended interaction with the primase/Gyr.

As we have seen so far, some structural variations in the molecules allowed SAR continuity that resulted in increased potency. On the other hand, discontinuity in SAR that markedly reduced the potency of an inhibitor after a small chemical modification, termed “the functional cliff”, also occurred. For example, we have revealed that the substituent on phenyl ring of indole may have a significant effect on the potency of the inhibitor. The presence of trifluoromethyl group (**13c** and **13d**) on the phenyl ring of the round III series of compounds led to a significant increase in the inhibitory effect compared to the chloro and nitro substituted compounds (**10a** and **10b**), respectively (Null activity). A detailed illustration of this functional cliff is presented in Supporting Information Figure S8, where three sub-structures that construct compounds **10c** and **10d**, and their corresponding analogs **13c** and **13d**, are defined (Supporting Information Figure S8, region A-blue, region B-red, and region C-green). The increase in potency upon modifications of region C and the distal parts of region B, whereas modifications of region A reduced potency, indicate that region A is the most important functional group of the pharmacophore.

Taken together, the SAR studies demonstrated that a variety of hydrophobic substituents introduced on a phenyl ring of indole were well tolerated, and among them the chloro, trifluoromethyl and nitro moiety were preferred. The carboxyethyl group at position 2, free carboxylic acid in position 3, an electron withdrawing group on phenyl ring of indole-3-carboxylic acid scaffold, as well as *N*-benzylation, are all essential for the biological activity (the summary of SAR conclusions is presented in Figure 3B).

Overall, out of 49 newly synthesized molecules, positional isomers 3-(1-benzyl-5-chloro-2-(ethoxycarbonyl)-4-(trifluoromethyl)-1H-indol-3-yl)propanoic acid (**13c**) and 3-(1-benzyl-7-chloro-2-(ethoxycarbonyl)-5-(trifluoromethyl)-1H-indol-3-yl)propanoic acid (**13d**) present the best inhibitors in all measured activities for DnaG and Gyr of *Mtb* (Figure 3C). Dose response of selected molecules on the DnaG/Gyr activities is presented in Supporting Information Figure S7. Structural characterization of all molecules is presented in Figures S9-S83. In view of our results, it is rationally presumed that these two inhibitors present a viable template to discover more potent compounds, by variations of substituents on the phenyl ring or - NH of indole.

### Some of the inhibitors stop the proliferation of *Mycobacterium smegmatis (Msmg)*

For this study we have used *Msmg*, a nonpathogenic relative of *Mtb* often used to test antitubercular agents due to the similar morphological and biochemical properties (e.g. DnaG primase as well as GyrB subunit, exhibit more than 80% sequence identity between *Mtb* and *Msmg*).

The primase/Gyr inhibitors were screened for inhibition of growth of *Msmg* mc_2_155 over a time course of 20h at 37°C. The antibacterial activity of molecules **13c** and **13d** were determined by assaying their minimal inhibitory concentration (MIC) against *Msmg*. The results are summarized in Figure 4. Both molecules inhibited the growth of *Msmg* in accordance with their inhibition of mycobacterial primase/Gyr. Other molecules that showed inhibition of the two enzymes *in vitro* had little or no effect against *Msmg* (Figure 4, growth curve, colored blue). The MIC_90_ of molecules **13c** and **13d** were 86.6 and 227 µM, respectively (Figure 4A). These values were comparable to the effective dose measured for isoniazid (50 µM), a first-line anti-TB drug and a central drug in common treatment regimens (Figure 4A, indicated in grey).

**Figure 4.**
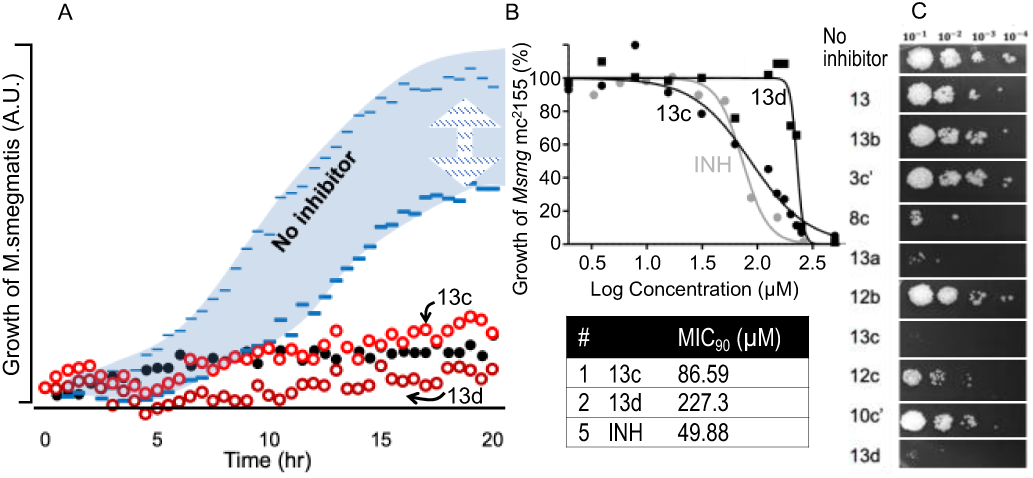
Effect of selected small molecule inhibitors *on Msmg* culture growth. (A) Growth curves of *Msmg* culture expressing mCherry reporter protein. The culture was grown in 7H9 liquid medium (0.4% glycerol, 0.05% Tween-80) containing different small molecule inhibitors in 250 µM concentration. The fluorescent signal originating form mCherry reporter protein was measured over the course of 20 hours and used to estimate viability of bacterial cells. Data represents the average of three replicates (n=3). The blue area represents the range observed for bacterial growth in the presence of different molecules presented in this study. The effect of 15 mg/L isoniazid (INH) is indicated in black. (B) The effect of molecules **13c** and **13d** on the viability of *Msmg* (compared to INH, indicated in grey). Table presents the MIC_90_ values of the three compounds. (C) Dilution colony assay. Aliquots of samples from the assay described in **A** were collected after 2h of incubation, serially diluted in 1:10 ratio. Each dilution was platted on a 7H10+ agar plate and incubated until bacterial colonies became visible.

## Conclusions

In summary, we described the rational design of small-molecules that inhibit activity of two DNA replication enzymes of *M. tuberculosis*. These enzymes were chosen as selective targets for anti-tuberculous agents, considering their pronounced structural differences when compared to human analogues.

Molecule **A13** that served as a promising inhibitor in the initial screening was chosen as a precursor for optimization. Three rounds of optimizations allowed us to draw SAR conclusions related to the inhibitor scaffold and design new molecules with improved inhibitory activity. Some molecules were found to inhibit both the DNA primase and the Gyr of *Mtb* and therefore were considered as dual acting inhibitors. Targeting several enzymes with common TOPRIM fold is a desirable goal for increasing potency and promoting bacterial cell toxicity. Although the IC_50_ values determined for each enzyme remained in the micromolar range (Supporting Information Figure S7), a few molecules were able to effectively inhibit the growth of bacterial culture (*Msmg*), presumably due to a dual mechanism of action (synergistic effect). The physicochemical and pharmacodynamic properties of the small-molecule inhibitors need to be optimized further in order to obtain new potent lead molecules. So far, compound 13c and 13d presented the best inhibitors, for both DNA replication enzymes in *Mtb*: DnaG primase and Gyr. Based on SAR analysis, we have suggested possible chemical manipulations that are expected to improve potency and physicochemical properties of these small molecules.

Our results offer valuable information on the design and synthesis of novel inhibitors for bacterial DNA replication as well as new treatment opportunities for *M. tuberculosis*.

## Experimental Section

### Materials

All chemicals and solvents used for the synthesis and characterization of compounds were purchased from Sigma-Aldrich and Acros, and were of reagent grade without purification or drying before use. Chemicals used for biochemical assays were molecular-biology grade, purchased from Sigma-Aldrich. Ribonucleotides (ATP, CTP, GTP, and UTP) were purchased from New England Biolabs (NEB). [α–_32_P] ATP (3000 Ci/mmol) and [γ–_32_P] ATP (3000 Ci/mmol) were purchased from Perkin Elmer. _1_H NMR data were recorded on a 400/500 MHz Bruker NMR instrument, and _13_C NMR data were recorded on a 101 MHz Bruker NMR instrument. Chemical shifts were reported in δ (ppm) downfield from tetramethylsilane (TMS) and were recorded in appropriate NMR solvents (CDCl_3_, δ 7.26; CD_3_OD, δ 3.31; (CD_3_)_2_CO, δ 2.05; or other solvents as mentioned). All coupling constants (J) are given in Hertz. All of the NMR spectra were processed by MestReNova software (www.mestrelab.com). The following abbreviations are used: s, singlet; d, doublet; t, triplet; m, multiplet; br s, broad singlet peak. The synthesized compounds were assessed for their purity determination using high performance liquid chromatography (HPLC). Purity of all final compounds was 95% or higher. Mass spectra were recorded only for the molecules that exhibited best activity, using an HRMS instrument. Thin-layer chromatography (TLC) was performed on Merck aluminum backed plates, pre-coated with silica (0.2mm, 60F254) and the detection of molecules was performed by UV fluorescence (254 nm). Column chromatography separations were performed using silica gel (0.04-0.06 mesh) as a stationary phase.

N,N-Dimethylformamide (DMF) and dichloromethane (DCM) were purchased as biotech grade from Avantor J.T. Baker. Analytical HPLC was performed on a Dionex 1100 using a reverse phase C18 column at a flow rate of 1.5 mL/min. Preparative HPLC was performed on a Dionex Ultimate 3000 instrument using a C18 reverse phase preparative column at a flow rate of 20 mL/min. Mass spectrometry analysis was performed by LC-MS Thermo Surveyor 355.

### Expression-purification of DnaG primase from *Mtb*

The DNA sequence encoding DnaG primase from *Mtb* H37Rv was inserted into pET28 vector. The transformed *Escherichia coli* BL21 (DE3) cells were grown in LB medium containing 50 mg/L kanamycin until OD_600_ = 0.7 and induced with 0.5 mM IPTG. After induction, the culture was incubated with shaking for 4h at 37°C. Cells were harvested by centrifugation (4700 x g, 15 minutes) and the pellet was resuspended in the lysis buffer (50 mM Tris-HCl pH 7.5, 500 mM NaCl, 10% v/v glycerol, and 1 mM dithiothreitol). Lysis was performed using pressure homogenizer. The lysate was clarified by centrifugation (21000 x g, 30 minutes), filtered through 0.45 µM filter and loaded onto Ni-NTA column (HisTrap FF, GE Healthcare). The column was equilibrated with buffer A (50 mM Tris-HCl pH 7.5, 500 mM NaCl, 10% v/v glycerol, 1 mM dithiothreitol and 10 mM imidazole) and the protein was eluted with a linear gradient of imidazole (10-500 mM) in buffer A. The fractions containing Mtb DnaG primase were joined and EDTA was added to a final concentration of 1 mM. Ammonium-sulfate was added (0.4 g/mL of sample) with stirring and left for several hours at 4°C to precipitate the protein. Centrifugation was performed (21000 x g, 30 minutes) and the protein precipitate was dissolved in buffer C (50 mM Tris-HCl pH 7.5, 500 mM NaCl, 10% v/v glycerol, 1 mM dithiothreitol, 1 mM EDTA) and loaded onto Superdex S200 column (GE Healthcare). The protein was eluted in buffer C. Fractions containing purified Mtb DnaG primase were joined and dialyzed against buffer C containing 50% glycerol. The protein samples were stored at −20 °C.

### Expression-purification of gyrase from *Mtb*

*Mtb* Gyr subunit A and B were purified separately. The DNA sequences encoding each Gyr subunit from *Mtb* H37Rv were inserted into separate pET28 vectors. The expression vectors were transformed into *Escherichia coli* BL21 (DE3) that were grown in LB medium containing 50 mg/L kanamycin until OD_600_ = 0.6 and induced with 0.5 mM IPTG and 0.2% arabinose. After induction, the culture was incubated with shaking for 4h at 30 °C. Cells were harvested by centrifugation (4700 x g, 15 minutes) and the pellet was resuspended in buffer A (50 mM Tris-HCl pH 8, 500 mM NaCl, 10 mM imidazole, 10% v/v glycerol, and 0.5 mM dithiothreitol). Lysis was performed using pressure homogenizer and sonication. The lysate was clarified by centrifugation (21000 x g, 60 minutes), filtered through 0.45 µM filter and loaded onto Ni-NTA column (HisTrap FF, GE Healthcare). The column was equilibrated with buffer A and the protein was eluted with a linear gradient of imidazole (10-400 mM) in buffer A. The fractions containing Mtb Gyr were joined and concentrated by dialysis against buffer B (20 mM Tris-HCl pH 8, 100 mM NaCl, 50% v/v glycerol, and 2 mM dithiothreitol) and loaded onto Superdex S200 column (GE Healthcare) The column was equilibrated and the protein was eluted with buffer C (20 mM Tris-HCl pH 8, 500 mM NaCl, 10% v/v glycerol, and 1 mM dithiothreitol). The fractions containing purified *Mtb* Gyr were joined and dialyzed against buffer B and stored at −20 °C.

### Oligoribonucleotide synthesis assay by *Mtb* DnaG primase. Standard

10 μL reaction contained 50 μM of DNA template (5’-CCGACCCGTCCGTAATACAGAGGTAATTGTCACGGT-3, that contain *Mtb* DnaG recognition site, found as reported in ^[24]^), 250 μM GTP, CTP, UTP and [α-_32_P]-ATP, and 500 nM *Mtb* DnaG primase in a buffer containing 40 mM Tris-HCl (pH 7.5), 1 mM MnCl_2_, 10 mM DTT, and 50 mM KGlu. The reaction was incubated at 37°C for 60 minutes with 4 mM of the small molecule for experiments reported in Figures 2-3, or 0.5 mM – 4 mM for experiments reported in Supporting Figure S7. The reaction was terminated by adding an equal volume of sequencing buffer containing 98% formamide, 0.1% bromophenol blue, and 20 mM EDTA. The samples were loaded onto a 25% polyacrylamide sequencing gel containing 7 M urea, and visualized by autoradiography. Quantification of the signal was performed by ImageQuant TL 8.1.

### DNA supercoiling assay for *Mtb* Gyr

DNA supercoiling activity was tested with purified *Mtb* GyrA and GyrB subunits mixed in 2:1 ratio. The reaction mixture contained 50 mM HEPES [pH 7.9], 100 mM potassium glutamate, 6 mM magnesium acetate, 2 mM spermidine, 4 mM dithiothreitol, 50 µg/ml bovine serum albumin, 1 mM ATP and 13.34 ng/µl relaxed pBR322 DNA, and 500 μM small molecules (10% V/V DMSO final). One unit of Gyr proteins was added (one unit of enzyme activity was defined as the amount of Gyr that converted 400 ng of relaxed pBR322 to the supercoiled form in 1 h at 37°C in 30 µl reaction), and the reaction mixtures were incubated at 37°C for 1 h. Reactions were terminated by adding one reaction volume of Stopping Buffer (40% sucrose, 100 mM Tris-HCl pH 8, 10 mM EDTA, 0.5 mg/ml Bromophenol Blue) and two reaction volumes of chloroform/isoamyl alcohol (v:v, 24:1). The samples were vortexed, centrifuged briefly and loaded onto 1% agarose gel in 1X Tris-borate-EDTA buffer, pH 8.3 (TBE). Electrophoresis was performed for 4h at 40 V, and the gel was subsequently stained with ethidium bromide (0.5 mg/ml).

### ATPase Norit assay for *Mtb* Gyr

The standard assay for ATPase measures the formation of Norit-adsorbable _32_P-labeled phosphate arising from the hydrolysis of [γ-_32_P]-ATP. ATPase assay was carried out according to the same reaction conditions as for DNA supercoiling except that 10 µM [γ-_32_P]-ATP 100 nCi was used instead of non-radioactive 1 mM ATP. The reaction mixture (10 μl) was incubated at 37 °C for 60 minutes. The reaction was stopped by the addition of 90 μl of a solution consisting of 0.6 ml of 1 M HCl, 0.1 M Na_4_P_2_O_7_, 0.02 M KH_2_PO_4_, 0.2 ml of a Norit A suspension (20% packed volume), and 0.1 ml of bovine serum albumin (5 mg/ml). The solutions were mixed and allowed to sit for 5 min at 0 °C. After centrifugation, 20 μl of the supernatant was spotted in a scintillation tube and upon the addition of Ultima Gold scintillation liquid (Perkin Elmer) the signal was measured by scintillation counter.

### Bacterial growth assays. Bacterial strains and growth media

*Msmg* cells mc_2_155 (wild type), containing Hygromycin resistance gene and transcriptional red fluorescent protein gene (mCherry), were grown in Middlebrook 7H9 broth (Fluka) liquid medium supplemented with glycerol (0.4%, v/v) and Tween-80 (0.05%, v/v) (7H9++ liquid medium). Solid medium was composed of Middlebrook 7H10 agar (Difco) and 0.4% (v/v) glycerol (7H10+ agar solid medium). Hygromycin was added to both liquid and solid media to a final concentration of 50 mg/L. The experiments were conducted under strictly sterile conditions. All instruments, materials and media involved were sterilized before use. Experimental procedures were performed in a biological hood with a laminar flow.

### Bacterial growth of *Msmg* culture

*Msmg* culture was first inoculated onto 7H10+ agar plate (medium components are described above) containing 50 mg/L hygromycin and incubated at 30°C for approximately four days. The primary liquid culture was prepared by transferring numerous colonies from a previously inoculated 7H10+ agar plate into 10 mL of 7H9++ liquid medium containing 50 mg/L hygromycin. Cells were incubated overnight (∼16 hours) at 30 °C with shaking (200 rpm). The secondary culture was prepared by transferring 1 mL of *Msmg* primary culture into 50 mL 7H9++ liquid medium (1:50 ratio, v/v) supplemented with 50 mg/L hygromycin. Again, cells were grown overnight at 30 °C with shaking (200 rpm) to mid-log phase (OD_600_ ∼0.5). Then the culture was diluted to OD_600_ = 0.025, dispensed in wells of Thermo Scientific™ Nunc™ F96 MicroWell™ black polystyrene plate and treated with different small molecule inhibitors at a concentration of 250 µM. The plate was incubated at 30°C with continuous orbital shaking (567 cpm) over the course of 20 hours. The cell growth was monitored simultaneously by Synergy H1 microplate reader (Biotek) using mCherry as a reporter protein (mCherry fluorescent signal directly correlates with the number of live bacterial cells). Data represent the average of three biological replicates (n=3).

### Dilution colony assay (*Msmg* colonies)

The samples from the previously described procedure (Bacterial growth of Msmg culture) were also used for dilution colony assay, as explained in the following text. After two hours of monitoring the bacterial growth in Synergy H1 microplate reader, 3μL of each sample was collected and serially diluted in 1:10 ratio (x10^−1^, x10^−2^,x10^−3^, x10^−4^). Then 3 μL of each dilution was transferred onto 7H10+ agar plate and incubated at 30°C for four days or until colonies were observed in the sample that didn’t contain any inhibitor.

### Synthesis of indole derivatives

The scheme for the synthesis of all molecules below is presented in Scheme 1 and Supporting Information Figure S5.

A. General procedures for the synthesis of compounds (2a-g).
  i. **Benzenediazonium salt derivatives** (mixture I). To a well-stirred solution of aniline derivatives 1a-g (10 mmol) in 5M aq. HCl (16 ml) at 0°C was added dropwise a cold solution (0–5°C) of sodium nitrite (1.38 g, 20 mmol, 2 equiv.) in 10 ml water. To maintain the reaction temperature at 0°C the addition rate kept slow. The resulting mixture was stirred at 0–5°C for an additional 30 min in an ice bath.
  ii. **2-(ethoxycarbonyl)cyclopentanone anion** (mixture II). A solution of 2-(ethoxycarbonyl)cyclopentanone (2.512 ml, 1.344g, 15 mmol) in ethanol (5 ml) was cooled to 0–5°C. Then, a potassium hydroxide solution (5.040 g, 90 mmol, 6 equiv.) in water (5 ml) previously cooled to 0–5°C was added dropwise for 30 min in order to keep the reaction temperature below 8°C. The white-milky appearance mixture was stirred for further 30 min at 0–5°C.
  iii. **Monoethyl derivatives of phenylhydrazones** (mixture I+ mixture II). Ice (50 g) was added to mixture II with stirring at 0–5°C in an ice bath, followed by the addition of mixture I, and stirring continued for 1 h at 50°C. The combined mixtures were brought to room temperature and the pH was subsequently adjusted to 4–5 with 1 M aq. HCl. The mixture was extracted with diethyl ether (50 ml×3) and the combined organic layer was dried over magnesium sulfate and filtered. The volatiles were removed under reduced pressure affording a gummy material (95–100%) that was used without further purification in the next step.
  iv. **Diethyl derivatives of phenylhydrazones (2a-g)**. Concentrated H_2_SO_4_ (2.7 ml, 50.5 mmol, 5.1 equiv.) was added dropwise to a stirred solution of monoethyl derivatives (10 mmol) in absolute ethanol (100 ml). The reaction mixture was then heated to reflux for 3 h at 100°C. Then the ethanol was removed under reduced pressure and the residue was treated with 100 ml of ice water. The aqueous solution was extracted with dichloromethane (3 × 50 ml). The combined organic layer was dried over magnesium sulfate, filtered and concentrated. The residue was purified by silica gel column chromatography (ethyl acetate:hexane, 1:4) led to the desired product in 80–95% yield as a yellow/brown solid.
B. General procedures for the synthesis of ethyl 3-(3-ethoxy-3-oxopropyl)-1H-indole-2-carboxylate derivatives (3a-g, 3c’). A mixture of p-toluenesulfonic acid (3.800 g, 20 mmol, 4 equiv.) and 100 ml of benzene was refluxed for 2 h at 110°C under azeotropic condition. Subsequently, 5 mmol of diethyl derivatives of phenylhydrazone (**2a**–**g**) was added and the mixture refluxed for 24 h before the volatiles were distilled. The mixture was washed with a saturated solution of NaHCO_3_ (2 × 30 ml) and water (30 ml). The crude reaction mixture was extracted with dichloromethane (3 × 30 ml) and then washed with water (3 × 20 ml). The organic layer was dried over magnesium sulfate, filtered, and concentrated under reduced pressure. The residue was purified by silica gel column chromatography using ethyl acetate/hexane (1:9 to 1:6) as eluent producing pale white/yellow solid (3a-g, 3c’) in 15–80% yield.
C. Synthesis of 3-(5-chloro-2-(hydroxymethyl)-1H-indol-3-yl)propan-1-ol (4a). Dry THF (20 ml) was added dropwise under N_2_ atmosphere to an ice-cooled RBF containing LiAlH_4_ (342 mg, 9.0 mmol, 6 equiv.) at 0°C. After completion of THF addition, conc. H_2_SO_4_ (240 µl, 4.5 mmol, 3 equiv.) was added to the reaction mixture and the resulting mixture was left to stir at 0°C for 30 min. To the stirred reaction mixture, compound 3a (1.5 mmol) in dry THF was slowly added within 30 min at 0°C and the resulting mixture was left to stir at room temperature for 2 h. After completion of the reaction, ice was added to the resulting reaction mixture and was filtered. The filter cake was washed with ethyl acetate (3 × 20 ml) and then with water (3 × 10 ml). The organic phase was collected, dried over magnesium sulfate and filtered. The solvent was removed under vacuum to produce grey solid. The residue was purified by silica gel column chromatography using methanol/dichloromethane (1:19) as eluent led to the final products in 50% yield as grey-white solid.
D. Synthesis of 5-chloro-3-(3-oxopropyl)-1H-indole-2-carbaldehyde (5a). To the solution of compound **4a** (0.05 mmol) in dry DCM (5 ml) was added PCC (32 mg, 0.15 mmol, 3 equiv). The reaction mixture was stirred for 3h at room temperature. After completion of the reaction, powdered MgSO_4_ was added to the resulting reaction mixture and then anhydrous ether (5 × 5 ml) was added, 5X of reaction solvent. The reaction mixture was then, filtered and concentrated. The residue was purified by silica gel column chromatography using ethyl acetate/hexane (1:4) as eluent led to the desired product in 40% yield as a pale white solid.
E. Synthesis of 5-chloro-3-(3-oxo-3-(phenylamino)propyl)-N-phenyl-1H-indole-2-carboxamide (7a). HOBt (25mg, 0.16 mmol, 2 equiv.) was added to a stirred solution of compound **6a** (0.08 mmole) in dry DMF (10 ml) under N_2_ atmosphere. Subsequently, the solution of aniline (15mg, 0.16 mmol, 2 equiv.) in dry DMF was added to the reaction mixture followed by the addition of DCC (34mg, 0.16 mmol, 2 equiv.) and DIPEA (5% equiv.). The resulting mixture was left to stir at room temperature for 48 h. After completion of the reaction, the reaction mixture was washed with 1M HCl (3 × 15 ml) followed by 0.1M Na_2_CO_3_ aqueous solution (3 × 20 ml) to quench the reaction. The crude mixture was extracted with ethyl acetate. The organic layer was then washed with brine (3 × 15 ml), dried over magnesium sulfate, filtered, and evaporated to dryness. The residue was purified by silica gel column chromatography (ethyl acetate:hexane, 1:6) as eluent led to compound 7a in 60% yield as a white solid.
F. General procedures for the synthesis of compounds (6a–g, 6c’), (8a-f, 8c’), (9a-f, 9c’) and (11a–f, 11c’), (12a-f, 12c’), (13a-f, 13c’). Compound **3a–g, 3c’** (0.5 mmol) and **10a-f, 10c’** (0.5 mmol) were dissolved in 50 ml tetrahydrofuran (THF) and the mixture stirred at room temperature for 1 h. Then a solution of NaOH (0.01M) was added and the reaction stirred at room temperature for 18-24 h. After completion of the reaction, the pH was adjusted to 2–3 using 1M aq. HCl, and the product was extracted with diethyl ether (3 × 20 ml). The combined organic layer was dried over magnesium sulfate, filtered, and evaporated to dryness. The residue was purified by silica gel column chromatography (ethyl acetate/hexane, 1:10 to 1:5) as eluent affording products **6a–g, 6c’, 8a-f, 8c’, 9a-f, 9c’** as well as **11a–f, 11c’, 12a-f, 12c’** and **13a-f, 13c’** in as white solids.
G. General Procedures for the Synthesis of N- benzylated derivatives of ethyl 3-(3-ethoxy-3-oxopropyl)-1H-indole-2-carboxylate (10a-f, 10c’). A catalytic amount of KI (8 mg, 0.05 mmol) and benzyl bromide (214 mg, 1.25 mmol, 2.5 equiv) were added to stirred solution of compounds **3a-f, 3c’** (0.5 mmol) and Cs_2_CO_3_ (488 mg, 1.5 mmol, 3 equiv) in DMF (20 ml). The reaction mixture was heated to 60°C for 2 h, and then concentrated under vacuum. The residue was dissolved in EtOAc (25 ml), washed with water (3 X10 ml), dried with anhydrous MgSO_4_ and concentrated under reduced pressure. The crude product was purified by silica gel column chromatography (ethyl acetate/hexane, 1:9) as eluent affording compounds **10a-f** and **10c’**) in high yields (85-95%) as yellow oils. Characterization of synthesized molecules

**Diethyl (*E*)-2-(2-(4-chlorophenyl)hydrazono)hexanedioate (2a)** 1H NMR (400 MHz, CDCl3) δ 7.14 – 7.02 (m, 4H), 4.27 (q, *J* = 7.1 Hz, 2H), 4.12 (q, *J* = 7.1 Hz, 2H), 2.54 (t, *J* = 7.3 Hz, 2H), 2.39 (t, *J* = 7.5 Hz, 2H), 1.96 (p, *J* = 7.5 Hz, 2H), 1.35 (t, *J* = 7.1 Hz, 3H), 1.24 (t, *J* = 7.1 Hz, 3H); 13C NMR (101 MHz, CDCl3) δ 173.9, 163.9, 141.4, 131.3, 129.9, 126.9, 113.7, 60.7, 60.4, 33.7, 32.1, 23.0, 20.8, 14.4.

**Diethyl (*E*)-2-(2-(4-nitrophenyl)hydrazono)hexanedioate (2b)** 1H NMR (400 MHz, CDCl3) δ 7.17 – 7.08 (m, 4H), 4.31 (q, *J* = 7.2 Hz, 2H), 4.21 (q, *J* = 7.2 Hz, 2H), 2.62 (t, *J* = 7.2 Hz, 2H), 2.40 (t, *J* = 7.5 Hz, 2H), 1.99 (p, *J* = 7.4 Hz, 2H), 1.39 (t, *J* = 7.2 Hz, 3H), 1.26 (t, *J* = 7.1 Hz, 3H); 13C NMR (101 MHz, CDCl3) δ 171.6, 165.8, 142.6, 132.2, 129.1, 125.5, 114.2, 61.6, 61.0, 34.8, 33.1, 23.7, 20.8, 15.6.

**Diethyl (*E*)-2-(2-(4-chloro-3- (trifluoromethyl)phenyl)hydrazono)hexanedioate (2c)** 1H NMR (400 MHz, CDCl3) δ 7.44 (d, *J* = 2.6 Hz, 1H), 7.35 (d, *J* = 8.7 Hz, 1H), 7.21 (dd, *J* = 8.7, 2.7 Hz, 1H), 4.27 (q, *J* = 7.1 Hz, 2H), 4.10 (q, *J* = 7.1 Hz, 2H), 2.55 (t, *J* = 7.4 Hz, 2H), 2.37 (t, *J* = 7.4 Hz, 2H), 1.94 (p, *J* = 7.4 Hz, 2H), 1.35 (t, *J* = 7.1 Hz, 3H), 1.22 (t, *J* = 7.1 Hz, 3H); 13C NMR (101 MHz, CDCl3) δ 173.6, 163.7, 142.5, 132.3, 130.3, 129.0, 123.5, 121.8, 117.3, 112.5, 61.2, 60.4, 33.6, 32.2, 22.7, 14.3, 14.2.

**Diethyl (*E*)-2-(2-(2-chloro-4- (trifluoromethyl)phenyl)hydrazono)hexanedioate (2d)** 1H NMR (400 MHz, CDCl3) δ 7.68 (d, *J* = 8.7 Hz, 1H), 7.57 (d, *J* = 1.7 Hz, 1H), 7.46 (dd, *J* = 8.7, 2.0 Hz, 1H), 4.34 (q, *J* = 7.1 Hz, 2H), 4.13 (q, *J* = 7.1 Hz, 2H), 2.61 (t, *J* = 7.3 Hz, 2H), 2.40 (t, *J* = 7.4 Hz, 2H), 1.99 (p, *J* = 7.4 Hz, 2H), 1.38 (t, *J* = 7.1 Hz, 3H), 1.24 (t, *J* = 7.2 Hz, 3H); 13C NMR (101 MHz, CDCl3) δ 173.6, 163.3, 142.9, 132.7, 126.8, 125.1, 123.6, 118.4, 114.0, 61.5, 60.5, 33.6, 32.4, 22.6, 14.4, 14.3.

**Diethyl (*E*)-2-(2-(2**,**5-dichloro-4-nitrophenyl)hydrazono)hexanedioate (2e)** 1H NMR (400 MHz, CDCl3) δ 8.12 (s, 1H), 7.69 (s, 1H), 4.36 (q, *J* = 7.1 Hz, 2H), 4.14 (q, *J* = 7.1 Hz, 2H), 2.64 (t, *J* = 7.4 Hz, 2H), 2.41 (t, *J* = 7.3 Hz, 2H), 2.04 – 1.95 (m, 2H), 1.39 (t, *J* = 7.1 Hz, 3H), 1.25 (t, *J* = 7.1 Hz, 3H); 13C NMR (101 MHz, CDCl3) δ 173.4, 163.2, 144.3, 139.4, 136.1, 128.9, 128.1, 116.2, 115.7, 62.1, 60.6, 33.5, 32.6, 22.4, 14.4, 14.2.

**Diethyl (*E*)-2-(2-(3**,**5- bis(trifluoromethyl)phenyl)hydrazono)hexanedioate (2f)** 1H NMR (400 MHz, CDCl3) δ 7.42 (d, *J* = 2.6 Hz, 1H), 7.30 (d, *J* = 1.7 Hz, 1H), 7.15 (d, *J* = 2.1Hz, 1H), 4.28 (q, *J* = 7.4 Hz, 2H), 4.15 (q, *J* = 7.2 Hz, 2H), 2.65 (t, *J* = 7.1 Hz, 2H), 2.39 (t, *J* = 7.4 Hz, 2H), 1.98 (p, *J* = 7.2 Hz, 2H), 1.37 (t, *J* = 7.2 Hz, 3H), 1.26 (t, *J* = 7.1 Hz, 3H); 13C NMR (101 MHz, CDCl3) δ 174.2, 161.9, 144.5, 133.4, 132.1, 129.7, 124.3, 123.9, 120.8, 115.2, 114.6, 60.8, 60.5, 33.4, 31.7, 21.4, 14.5, 14.2.

**Diethyl (*E*)-2-(2-(p-tolyl)hydrazono)hexanedioate (2g)** 1H NMR (400 MHz, CDCl3) δ 7.15 – 7.01 (m, 4H), 4.24 (q, *J* =7.2Hz, 2H), 4.15 (q, *J* = 7.2 Hz, 2H), 2.58 (s, 3H), 2.53 (t, *J* = 7.2 Hz, 2H), 2.40 (t, *J* = 7.3 Hz, 2H), 1.98 (p, *J* = 7.6 Hz, 2H), 1.37 (t, *J* = 7.1 Hz, 3H), 1.22 (t, *J* = 7.1 Hz, 3H); 13C NMR (101 MHz, CDCl3) δ 172.9, 164.0, 140.9, 130.3, 129.3, 128.5, 112.7, 61.6, 60.9, 35.4, 33.6, 31.8, 24.3, 22.8, 15.4.

**Ethyl 5-chloro-3-(3-ethoxy-3-oxopropyl)-1H-indole-2-carboxylate (3a)** TLC (EtAc/Hex 15:85) Rf = 0.3; 70% yield;orange-brown solid; 1H NMR (400 MHz, CDCl3) δ 9.26 (s, 1H), 8.73 (d, *J* = 2.1 Hz, 1H), 8.20 (dd, *J* = 9.1, 2.2 Hz, 1H), 7.42 (d, *J* = 9.1 Hz, 1H), 4.46 (q, *J* = 7.1 Hz, 2H), 4.11 (q, *J* = 7.1 Hz, 2H), 3.44 (t, *J* = 7.7 Hz, 2H), 2.72 (t, *J* = 7.6 Hz, 2H), 1.45 (t, *J* = 7.1 Hz, 3H), 1.23 (t, *J* = 7.2 Hz, 3H); 13C NMR (101 MHz, CDCl3) δ 172.8, 161.5, 142.3, 138.3, 127.2, 126.7, 125.0, 120.9, 118.6, 112.2, 61.8, 60.8, 35.4, 20.4, 14.4, 14.3.

**Ethyl 3-(3-ethoxy-3-oxopropyl)-5-nitro-1H-indole-2-carboxylate (3b)** TLC (EtAc/Hex 15:85) Rf = 0.2; 53% yield; pale-yellow solid; 1H NMR (400 MHz, CDCl3) δ 9.20 (s, 1H), 8.73 (d, *J* = 2.1 Hz, 1H), 8.21 (dd, *J* = 9.1, 2.2 Hz, 1H), 7.42 (d, *J* = 9.1 Hz, 1H), 4.46 (q, *J* = 7.1 Hz, 2H), 4.11 (q, *J* = 7.1 Hz, 2H), 3.44 (t, *J* = 7.7 Hz, 2H), 2.71 (t, *J* = 7.7 Hz, 2H), 1.45 (t, *J* = 7.1 Hz, 3H), 1.23 (t, *J* = 7.2 Hz, 3H); 13C NMR (101 MHz, CDCl3) δ 173.5, 172.8, 142.3, 138.3, 127.3, 126.7, 125.0, 120.9, 118.6, 112.1, 61.8, 60.8, 35.4, 20.4, 14.5, 14.3.

**Ethyl 5-chloro-3-(3-ethoxy-3-oxopropyl)-4-(trifluoromethyl)-1H- indole-2-carboxylate (3c)** TLC (EtAc/Hexane15:85) Rf = 0.2; 25% yield; off-white solid; 1H NMR (400 MHz, CDCl3) δ 9.39 (s, 1H), 7.47 (d, *J* = 8.8 Hz, 1H), 7.35 (d, *J* = 8.8 Hz, 1H), 4.45 (q, *J* = 7.1 Hz, 2H), 4.17 (q, *J* = 7.1 Hz, 2H), 3.53 – 3.38 (m, 2H), 2.70 – 2.54 (m, 3H), 1.43 (t, *J* = 7.1 Hz, 3H), 1.27 (t, *J* = 7.1 Hz, 3H); 13C NMR (101 MHz, CDCl3) δ 173.2, 161.5, 135.2, 128.6, 127.1, 126.4, 126.4, 125.2, 124.6, 122.5, 116.7, 61.8, 60.5, 35.4, 21.3, 21.2, 14.4.

**Ethyl 5-chloro-3-(3-ethoxy-3-oxopropyl)-6-(trifluoromethyl)-1H- indole-2-carboxylate (3c’)** TLC (EtAc/Hex 15:85)Rf = 0.3; 20% yield; off-white solid; 1H NMR (400 MHz, CDCl3) δ 9.19 (s, 1H), 7.85 (s, 1H), 7.76 (s, 1H), 4.45 (q, J = 7.1 Hz, 2H), 4.10 (q, J = 7.2 Hz, 2H), 3.36 (t, J = 7.7 Hz, 2H), 2.67 (t, J = 7.7 Hz, 2H), 1.44 (t, J = 7.1 Hz, 3H), 1.22 (t, J = 7.2 Hz, 3H); 13C NMR (101 MHz, CDCl3) δ 172.9, 161.5, 132.7, 130.5, 127.2, 124.8, 123.4, 123.0, 122.0, 112.1, 112.0, 61.7, 60.7, 35.3, 20.3, 14.4, 14.3; HRMS (ESI) calcd for C17H17ClF3NO4 for [M+ H] 392.0871, found 386.0857; HPLC purity 100% (tR = 8.35).

**Ethyl 7-chloro-3-(3-ethoxy-3-oxopropyl)-5-(trifluoromethyl)-1H- indole-2-carboxylate (3d)** TLC (EtAc/Hex 15:85) Rf= 0.3; 30% yield; white solid; 1H NMR (400 MHz, CDCl3) δ 9.07 (s, 1H), 8.08 – 7.81 (m, 1H), 7.56 (d, *J* = 1.0 Hz, 1H), 4.47 (q, *J* = 7.1 Hz, 2H), 4.09 (q, *J* = 7.1 Hz, 2H), 3.53 – 3.27 (m, 2H), 2.68 (t, *J* = 7.7 Hz, 2H), 1.46 (t, *J* = 7.1 Hz, 3H), 1.21 (t, *J* = 7.2 Hz, 4H); 13C NMR (101 MHz, CDCl3) δ 172.8, 161.3, 134.3, 128.3, 126.0, 124.6, 124.1, 121.4, 118.0, 117.7, 117.6, 61.7, 60.7, 35.3, 20.5, 14.5, 14.3.

**Ethyl 4**,**7-dichloro-3-(3-ethoxy-3-oxopropyl)-5-nitro-1H-indole-2- carboxylate (3e)** TLC (EtAc/Hex 15:85) Rf = 0.2;20% yield; pale-yellow solid; 1H NMR (400 MHz, CDCl3) δ 9.26 (s, 1H), 7.87 (s, 1H), 4.48 (q, *J* = 7.1 Hz, 2H), 4.15 (q, *J* = 7.1 Hz, 2H), 3.90 – 3.61 (m, 2H), 2.92 – 2.51 (m, 2H), 1.46 (t, *J* = 7.1 Hz, 3H), 1.25 (t, *J* = 7.1 Hz, 3H); 13C NMR (101 MHz, CDCl3) δ 172.5, 160.8, 142.5, 134.8, 127.8, 126.5, 125.1, 122.1, 121.4, 116.4, 62.2, 60.7, 36.0, 20.8, 14.4, 14.4.

**Ethyl 3-(3-ethoxy-3-oxopropyl)-4**,**6-bis(trifluoromethyl)-1H-indole-2- carboxylate (3f)** TLC (EtAc/Hex 15:85) Rf =0.5; 42% yield; off-white solid; 1H NMR (400 MHz, CDCl3) δ 10.35 (s, 1H), 7.74 (s, 1H), 7.39 (s, 1H), 4.37 – 4.31 (m, 2H), 4.31 – 4.25 (m, 2H), 2.71 – 2.59 (m, 2H), 2.55 – 2.45 (m, 2H), 1.40 (t, *J* = 7.1 Hz, 3H), 1.34 (t, *J* = 7.1 Hz, 3H);13C NMR (101 MHz, CDCl3) δ 176.1, 164.9, 145.3, 138.0, 132.7, 132.4, 127.6, 124.9, 122.9, 122.2, 114.3, 113.8, 61.7, 61.6, 32.2, 24.8, 14.4, 14.2.

**Ethyl 3-(3-ethoxy-3-oxopropyl)-5-methyl-1H-indole-2-carboxylate (3g)** TLC (EtAc/Hex 15:85) Rf = 0.3; 80% yield;pale white solid; 1H NMR (500 MHz, CDCl3) δ 8.97 (s, 1H), 7.58 (s, 1H), 7.37 (d, J = 8.0 Hz, 1H), 7.25 (d, J = 8.3 Hz, 1H), 4.52 (q, J = 6.7 Hz, 2H), 4.23 (q, J = 6.7 Hz, 2H), 3.50 (t, J = 7.7 Hz, 2H), 2.78 (t, J = 7.7 Hz, 2H), 2.56 (s, 3H), 1.53 (t, J = 6.8 Hz, 3H), 1.33 (t, J = 6.9 Hz, 3H); 13C NMR (126 MHz, CDCl3) δ 173.1, 162.1, 134.1, 129.4, 127.7, 127.5, 123.4, 122.1, 119.8, 111.3, 60.7, 60.2, 35.3, 21.3, 20.3, 14.2, 14.0.

**3-(5-chloro-2-(hydroxymethyl)-1H-indol-3-yl)propan-1-ol (4a)** TLC (EtAc/Hex 15:85) Rf = 0.3; 80% yield; pale whitesolid; 1H NMR (500 MHz, MeOD) δ 7.44 (dd, J = 2.0, 0.4 Hz, 1H), 7.22 (dd, J = 8.6, 0.5 Hz, 1H), 6.98 (dd, J = 8.6, 2.0 Hz, 1H), 3.51 (t, J = 6.3 Hz, 2H), 3.27 (dt, J = 3.3, 1.6 Hz, 2H), 2.83 – 2.74 (m, 2H), 1.86 – 1.75 (m, 2H); 13C NMR (126 MHz, MeOD) δ 137.3, 136.0, 130.7, 125.4, 122.6, 119.1, 113.3, 113.0, 62.3, 56.5, 34.8, 21.0.

**5-chloro-3-(3-oxopropyl)-1H-indole-2-carbaldehyde (5a)** TLC (EtAc/Hex 20:80) Rf = 0.3; 60% yield; pale yellow solid;1H NMR (400 MHz, Chloroform-d) δ 10.12 (s, 1H), 9.83 (s, 1H), 7.67 (s, 1H), 7.34 (d, J = 1.3 Hz, 2H), 3.38 (t, J = 7.2 Hz, 2H), 2.95 (t, J = 7.2 Hz, 2H); 13C NMR (101 MHz, DMSO) δ 202.7, 182.4, 135.9, 133.1, 127.5, 126.6, 125.4, 124.5, 120.4, 114.6, 44.6, 15.7.

**3-(2-carboxyethyl)-5-chloro-1H-indole-2-carboxylic acid (6a)** TLC (MeOH/DCM 1:9) Rf = 0.5; 20% yield; off-white solid; 1H NMR (400 MHz, MeOD) δ 7.68 (d, *J* = 1.5 Hz, 1H), 7.37 (d, *J* = 8.7 Hz, 1H), 7.20 (dd, *J* = 8.8, 1.9 Hz, 1H), 3.35 (t, *J* = 7.7 Hz, 2H), 2.63 (t, *J* = 7.7 Hz, 2H); 13C NMR (101 MHz, MeOD) δ 177.1, 164.7, 136.0, 129.7, 126.6, 126.4, 126.3, 122.7, 120.6, 114.5, 36.2, 21.2.

**3-(2-carboxyethyl)-5-nitro-1H-indole-2-carboxylic acid (6b)** TLC (MeOH/DCM 1:9) Rf = 0.4; 23% yield; pale-yellow solid; 1H NMR (400 MHz, MeOD) δ 8.74 (d, *J* = 2.2 Hz, 1H), 8.13 (dd, *J* = 9.1, 2.2 Hz, 1H), 7.50 (d, *J* = 9.1 Hz, 1H), 3.44 (t, *J* = 7.6 Hz, 2H), 2.69 (t, *J* = 7.6 Hz, 2H); 13C NMR (101 MHz, MeOD) δ 176.9, 169.6, 151.7, 143.0, 140.2, 128.1, 125.4, 120.6, 119.1, 113.4, 36.1, 21.1.

**3-(2-carboxyethyl)-5-chloro-4-(trifluoromethyl)-1H-indole-2- carboxylic acid (6c)** TLC (MeOH/DCM 1:9) Rf = 0.3; 15% yield; white solid; 1H NMR (400 MHz, MeOD) δ 7.62 (dd, *J* = 8.8, 0.7 Hz, 1H), 7.36 (dd, *J* = 8.8, 0.6 Hz, 1H), 3.48 – 3.38 (m, 2H), 2.62 – 2.53 (m, 2H); 13C NMR (101 MHz, MeOD) δ 177.1, 164.1, 137.2, 129.3, 128.5, 126.9, 126.4, 125.0, 122.3, 118.7, 35.9, 21.9.

**3-(2-carboxyethyl)-5-chloro-6-(trifluoromethyl)-1H-indole-2- carboxylic acid (6c’)** TLC (MeOH/DCM 1:9) Rf = 0.3; 15% yield; white solid; 1H NMR (400 MHz, Acetone-d6) δ 8.10 (s, 1H), 8.00 (s, 1H), 3.44 (t, J = 7.6 Hz, 2H), 2.74 (t, J = 7.6 Hz, 2H); 13C NMR (101 MHz, Acetone-d6) δ 174.1, 162.9, 133.9, 131.4, 126.1, 124.3, 122.3, 122.0, 113.4, 113.3, 35.3, 20.6.

**3-(2-carboxyethyl)-7-chloro-5-(trifluoromethyl)-1H-indole-2- carboxylic acid (6d)** TLC (MeOH/DCM 1:9) Rf = 0.3; 18% yield; off-white solid; 1H NMR (400 MHz, Acetone-d6) δ 8.19 (s, 1H), 7.59 (s, 1H), 3.47 (t, *J* = 7.5 Hz, 2H), 2.74 (t, *J* = 7.6 Hz, 2H); 13C NMR (101 MHz, Acetone-d6) δ 173.3, 162.4, 134.2, 132.7, 128.6, 128.3, 123.6, 122.4, 120.1, 119.9, 117.6, 34.3, 19.7.

**3-(2-carboxyethyl)-4**,**7-dichloro-5-nitro-1H-indole-2-carboxylic acid (6e)** TLC (MeOH/DCM 1:9) Rf = 0.3; 10% yield; pale-yellow solid; 1H NMR (400 MHz, MeOD) δ 7.86 (s, 1H), 3.81 – 3.57 (m, 2H), 2.72 – 2.54 (m, 2H); 13C NMR (101 MHz, MeOD) δ 176.5, 163.7, 143.4, 139.1, 136.4, 131.4, 126.2, 125.9, 121.5, 117.9, 36.8, 21.7.

**3-(2-carboxyethyl)-4**,**6-bis(trifluoromethyl)-1H-indole-2-carboxylic acid (6f)** TLC (MeOH/DCM 1:9) Rf = 0.4; 16%yield; off-white solid; 1H NMR (400 MHz, MeOD) δ 8.04 (s, 1H), 7.67 (s, 1H), 3.52 – 3.43 (m, 2H), 2.64 – 2.52 (m, 2H); 13C NMR (101 MHz, MeOD) δ 176.8, 163.7, 137.6, 130.9, 126.8, 126.2, 125.9, 125.1, 124.5, 124.2, 121.7, 115.7, 35.8, 21.2.

**3-(2-carboxyethyl)-5-methyl-1H-indole-2-carboxylic acid (6g)** TLC (MeOH/DCM 1:9) Rf = 0.5; 20% yield; off-whitesolid; 1H NMR (400 MHz, MeOD) δ 7.45 (s, 1H), 7.29 (d, J = 8.4 Hz, 1H), 7.09 (d, J = 8.5 Hz, 1H), 3.40 – 3.33 (m, 2H), 2.62 (t, J = 7.9 Hz, 2H), 2.42 (s, 3H); 13C NMR (101 MHz, MeOD) δ 177.4, 165.4, 136.3, 130.0, 128.8, 128.1, 125.0, 122.9, 120.4, 112.9, 36.3, 21.6, 21.3.

**5-chloro-3-(3-oxo-3-(phenylamino)propyl)-N-phenyl-1H-indole-2- carboxamide (7a**) TLC (EtAc/Hex 20:80) Rf = 0.3; 60% yield; pale white solid;1H NMR (400 MHz, Acetone-d6) δ 11.38 (s, 1H), 10.76 (s, 1H), 9.56 (s, 1H), 8.03 – 7.98 (m, 2H), 7.74 (d, J = 1.8 Hz, 1H), 7.62 – 7.58 (m, 2H), 7.52 (d, J = 8.8 Hz, 1H), 7.43 – 7.36 (m, 2H), 7.29 – 7.20 (m, 3H), 7.13 (t, J = 7.4 Hz, 1H), 7.03 (t, J = 7.4 Hz, 1H), 3.51 – 3.39 (m, 2H), 3.27 – 3.18 (m, 2H). 13C NMR (126 MHz, Acetone-d6) δ 172.8, 160.1, 139.8, 138.6, 134.7, 131.2, 128.8, 124.7, 124.2, 123.9, 123.6, 119.8, 119.8, 119.6, 119.5, 119.4, 114.3, 113.8, 36.0, 19.5.

**5-chloro-3-(3-ethoxy-3-oxopropyl)-4-(trifluoromethyl)-1H-indole-2- carboxylic acid (8c)** TLC (EtAc/Hex 40:60) Rf =0.5; 53% yield; pale-brown solid; 1H NMR (400 MHz, MeOD) δ 7.62 (d, J = 8.8 Hz, 1H), 7.35 (d, J = 8.8 Hz, 1H), 4.14 (q, J = 7.0 Hz, 2H), 3.50 – 3.38 (m, 2H), 2.63 – 2.53 (m, 2H), 1.26 (t, J = 6.8 Hz, 3H); 13C NMR (101 MHz, MeOD) δ 175.3, 169.0, 139.3, 137.3, 131.4, 128.7, 126.7, 125.1, 122.2, 120.5, 118.9, 61.7, 36.4, 22.1, 14.7. HRMS (ESI) calcd for C15H13ClF3NO4 for [M+ Na] 386.0377, found 386.0367.

**7-chloro-3-(3-ethoxy-3-oxopropyl)-5-(trifluoromethyl)-1H-indole-2- carboxylic acid (8d)** TLC (EtAc/Hex 40:60) Rf =0.6; 33% yield; off-white solid; 1H NMR (400 MHz, Acetone-d6) δ 8.21 (d, J = 0.7 Hz, 1H), 7.66 – 7.57 (m, 1H), 4.42 (q, J = 7.1 Hz, 2H), 3.47 – 3.39 (m, 2H), 2.67 (t, J = 7.7 Hz, 2H), 1.39 (t, J = 7.1 Hz, 3H); 13C NMR (101 MHz, Acetone-d6) δ 174.1, 161.8, 134.0, 131.3, 128.6, 126.0, 124.4, 123.3, 122.5, 122.2, 113.4, 61.8, 35.2, 20.6, 14.5; HRMS (ESI) calcd for C15H13ClF3NO4 for [M+ Na] 386.0377, found 386.0363.

**Ethyl 5-chloro-3-(propanoic acid)-1H-indole-2-carboxylate (9a)** TLC (EtAc/Hex 40:60) Rf = 0.6; 62% yield; off-whitesolid; 1H NMR (400 MHz, Acetone-d6) δ 7.82 (d, *J* = 2.0 Hz, 1H), 7.50 (d, *J* = 8.8 Hz, 1H), 7.26 (dd, *J* = 8.8, 2.0 Hz, 1H), 4.39 (q, *J* = 7.1 Hz, 2H), 3.47 – 3.28 (m, 2H), 2.74 – 2.60 (m, 2H), 1.39 (t, *J* = 7.1 Hz, 3H); 13C NMR (101 MHz, Acetone-d6) δ 173.2, 161.3, 134.7, 128.4, 125.1, 125.0, 124.9, 121.7, 119.7, 113.7, 60.4, 34.4, 19.8, 13.6.

**3-(2-(ethoxycarbonyl)-5-nitro-1H-indol-3-yl)propanoic acid (9b)** TLC (EtAc/Hex 40:60) Rf = 0.5; 40% yield; pale-yellow solid; 1H NMR (400 MHz, Acetone-d6) δ 8.83 (d, *J* = 2.2 Hz, 1H), 8.17 (dd, *J* = 9.1, 2.2 Hz, 1H), 7.65 (d, *J* = 9.1 Hz, 1H), 4.43 (q, *J* = 7.1 Hz, 2H), 3.48 (t, *J* = 7.6 Hz, 2H), 2.73 (t, *J* = 7.6 Hz, 2H), 1.41 (t, *J* = 7.1 Hz, 3H); 13C NMR (101 MHz, Acetone- d6) δ 174.0, 161.9, 142.8, 139.7, 131.8, 127.7, 125.6, 120.7, 119.2, 113.6, 61.8, 35.4, 20.7, 14.6.

**Ethyl 5-chloro-3-(3-ethoxy-3-oxopropyl)-6-(trifluoromethyl)-1H- indole-2-carboxylate (9c’)** TLC (EtAc/Hex 40:60)Rf = 0.6; 41% yield; off-white solid; 1H NMR (400 MHz, Acetone-d6) δ 8.08 (s, 1H), 7.97 (s, 1H), 4.43 (q, *J* = 7.1 Hz, 2H), 3.40 (t, *J* = 7.6 Hz, 2H), 2.70 (t, *J* = 7.6 Hz, 2H), 1.41 (t, *J* = 7.1 Hz, 3H); 13C NMR (101 MHz, Acetone-d6) δ 174.0, 161.8, 134.0, 131.3, 128.6, 126.0, 124.4, 123.3, 122.5, 122.2, 113.4, 61.8, 35.2, 20.6, 14.5.

**Ethyl 1-benzyl-5-chloro-3-(3-ethoxy-3-oxopropyl)-1H-indole-2- carboxylate (10a)** TLC (EtAc/Hex 15:85) Rf = 0.4;95% yield;pale brown liquid; 1H NMR (400 MHz, CDCl3) δ 7.74 (s, 1H), 7.30 – 7.27 (m, 2H), 7.26 – 7.20 (m, 3H), 7.00 (d, *J* = 6.7 Hz, 2H), 5.78 (s, 2H), 4.37 (q, *J* = 7.1 Hz, 2H), 4.15 (q, *J* = 7.1 Hz, 2H), 3.49 – 3.28 (m, 2H), 2.78 – 2.62 (m, 2H), 1.37 (t, *J* = 7.1 Hz, 3H), 1.26 (t, *J* = 7.1 Hz, 3H); 13C NMR (101 MHz, CDCl3) δ 182.0, 173.1, 138.2, 137.0, 128.7, 127.6, 127.3, 126.3, 126.2, 126.1, 125.9, 123.3, 120.1, 112.0, 61.1, 60.6, 48.5, 35.7, 21.3, 14.3, 14.2.

**Ethyl 1-benzyl-3-(3-ethoxy-3-oxopropyl)-5-nitro-1H-indole-2- carboxylate (10b)** TLC (EtAc/Hex 15:85) Rf = 0.3;90% yield;pale-yellow liquid; 1H NMR (400 MHz, CDCl3) δ 8.76 (d, *J* = 2.2 Hz, 1H), 8.19 (dd, *J* = 9.2, 2.2 Hz, 1H), 7.38 (d, *J* = 9.3 Hz, 1H), 7.32 – 7.22 (m, 3H), 7.00 (d, *J* = 6.5 Hz, 2H), 5.83 (s, 2H), 4.39 (q, *J* = 7.1 Hz, 2H), 4.13 (q, *J* = 7.1 Hz, 2H), 3.52 – 3.40 (m, 2H), 2.77 – 2.67 (m, 2H), 1.37 (t, *J* = 7.1 Hz, 3H), 1.24 (t, *J* = 7.2 Hz, 3H); 13C NMR (101 MHz, CDCl3) δ 172.7, 161.6, 146.8, 142.4, 140.9, 137.3, 128.9, 127.9, 127.7, 126.2, 126.0, 120.7, 118.5, 111.1, 61.5, 60.7, 48.9, 35.6, 21.2, 14.3, 14.2.

**Ethyl-1-benzyl-5-chloro-3-(3-ethoxy-3-oxopropyl)-4-(trifluoromethyl)- 1H-indole-2-carboxylate (10c)** TLC(EtAc/Hex 15:85) Rf = 0.5; 87% yield; pale-brown liquid; 1H NMR (400 MHz, CDCl3) δ 7.40 (d, *J* = 9.0 Hz, 1H), 7.32 (d, *J* = 8.9 Hz, 1H), 7.31 – 7.19 (m, 3H), 6.96 (d, *J* = 7.3 Hz, 2H), 5.72 (s, 2H), 4.38 (q, *J* = 7.1 Hz, 2H), 4.17 (q, *J* = 7.0 Hz, 2H), 3.55 – 3.35 (m, 2H), 2.76 – 2.55 (m, 2H), 1.33 (t, *J* = 7.1 Hz, 3H), 1.28 (t, *J* = 7.0 Hz, 3H); 13C NMR (101 MHz, CDCl3) δ 173.2, 162.1, 137.7, 137.3, 133.4, 129.7, 128.9, 128.2, 127.7, 126.6, 126.0, 125.3, 123.8, 121.9, 115.5, 61.8, 60.4, 48.7, 35.8, 21.7, 14.4, 14.1.

**Ethyl-1-benzyl-5-chloro-3-(3-ethoxy-3-oxopropyl)-6-(trifluoromethyl)- 1H-indole-2-carboxylate (10c’)** TLC(EtAc/Hex 15:85) Rf = 0.4; 90% yield; pale-brown liquid; 1H NMR (400 MHz, CDCl3) δ 7.88 (s, 1H), 7.69 (s, 1H), 7.31– 7.20 (m, 3H), 6.97 (d, *J* = 6.8 Hz, 2H), 5.79 (s, 2H), 4.37 (q, *J* = 7.0 Hz, 2H), 4.13 (q, *J* = 7.2 Hz, 2H), 3.45 – 3.33 (m, 2H), 2.79 – 2.53 (m, 2H), 1.36 (t, *J* = 7.0 Hz, 3H), 1.24 (t, *J* = 7.2 Hz, 3H); 13C NMR (101 MHz, CDCl3) δ 172.9, 161.7, 137.5, 135.6, 129.3, 128.9, 128.4, 127.7, 126.1, 124.8, 123.4, 123.1, 122.9, 122.1, 110.9, 61.5, 60.7, 48.7, 35.6, 21.1, 14.3, 14.2.

**Ethyl-1-benzyl-7-chloro-3-(3-ethoxy-3-oxopropyl)-5-(trifluoromethyl)- 1H-indole-2-carboxylate (10d)** TLC(EtAc/Hex 15:85) Rf = 0.5; 85% yield; pale-brown liquid; 1H NMR (400 MHz, CDCl3) δ 7.95 (d, *J* = 0.7 Hz, 1H), 7.53 (d, *J* = 1.1 Hz, 1H), 7.31 – 7.17 (m, 3H), 6.89 (d, *J* = 6.7 Hz, 2H), 6.28 (s, 2H), 4.36 (q, *J* = 7.1 Hz, 2H), 4.13 (q, *J* = 7.1 Hz, 2H), 3.47 – 3.33 (m, 2H), 2.74 – 2.61 (m, 2H), 1.35 (t, *J* = 7.1 Hz, 3H), 1.23 (t, *J* = 7.1 Hz, 3H); 13C NMR (101 MHz, CDCl3) δ 172.8, 161.6, 139.4, 134.9, 129.1, 128.9, 128.7, 127.2, 125.7, 124.9, 124.1, 123.4, 118.3, 117.4, 117.3, 61.5, 60.7, 49.3, 35.6, 21.0, 14.3, 14.1.

**1-benzyl-3-(2-carboxyethyl)-5-chloro-6-(trifluoromethyl)-1H-indole-2- carboxylic acid (11c’)** TLC (MeOH/DCM 1:9)Rf = 0.3; 15% yield; pale white solid; 1H NMR (400 MHz, Acetone-d6) δ 8.13 (s, 1H), 8.02 (s, 1H), 7.30 – 7.15 (m, 3H), 7.06 (d, J = 6.9 Hz, 2H), 6.01 (s, 2H), 3.45 (t, J = 7.6 Hz, 2H), 2.74 (t, J = 7.6 Hz, 2H); 13C NMR (101 MHz, Acetone-d6) δ 173.2, 162.2, 138.2, 135.3, 129.5, 129.4, 128.4, 127.1, 126.1, 125.0, 123.5, 122.7, 122.3, 121.6, 111.3, 111.3, 47.8, 34.4, 20.3.

**1-benzyl-5-chloro-3-(3-ethoxy-3-oxopropyl)-4-(trifluoromethyl)-1H- indole-2-carboxylic acid (12c)** TLC (EtAc/Hex40:60) Rf = 0.7; 30% yield; pale-brown solid; 1H NMR (400 MHz, MeOD) δ 7.64 (d, J = 9.0 Hz, 1H), 7.36 (d, J = 8.9 Hz, 1H), 7.24 (dq, J = 14.3, 7.0 Hz, 3H), 7.03 (d, J = 7.4 Hz, 2H), 5.83 (s, 2H), 4.15 (q, J = 7.1 Hz, 2H), 3.47 – 3.34 (m, 2H), 2.72 – 2.60 (m, 2H), 1.26 (t, J = 7.1 Hz, 3H); 13C NMR (101 MHz, MeOD) δ 183.1, 175.5, 147.7, 139.3, 139.0, 139.0, 137.7, 129.6, 128.2, 127.5, 126.6, 126.2, 125.3, 117.0, 116.4, 61.4, 49.6, 36.9, 22.5, 14.5; HRMS (ESI) calcd for C22H19ClF3NO4 for [M+ H] 454.1028, found 454.1026.

**3-(1-benzyl-5-chloro-2-(ethoxycarbonyl)-1H-indol-3-yl)propanoic acid (13a)** TLC (EtAc/Hex 40:60) Rf = 0.7; 75%yield; pale-brown solid; 1H NMR (400 MHz, MeOD) δ 7.75 (d, *J* = 1.5 Hz, 1H), 7.37 (d, *J* = 8.9 Hz, 1H), 7.25 (d, *J* = 2.0 Hz, 1H), 7.24 – 7.14 (m, 3H), 6.94 (d, *J* = 6.9 Hz, 2H), 5.78 (s, 2H), 4.33 (q, *J* = 7.1 Hz, 2H), 3.36 (t, *J* = 7.7 Hz, 2H), 2.62– 2.58 (m, 2H), 1.33 (t, *J* = 7.1 Hz, 3H); 13C NMR (101 MHz, MeOD) δ 176.9, 163.3, 139.8, 138.3, 129.5, 128.7, 128.1, 127.2, 126.9, 124.5, 124.1, 120.9, 113.4, 111.6, 62.0, 49.6, 36.4, 22.0, 14.4; HRMS (ESI) calcd for C21H20ClNO4 for [M+ Na] 408.0973, found 408.0962.

**3-(1-benzyl-2-(ethoxycarbonyl)-5-nitro-1H-indol-3-yl)propanoic acid (13b)** TLC (EtAc/Hex 40:60) Rf = 0.7; 54%yield; pale-yellow solid; 1H NMR (400 MHz, MeOD) δ 8.74 (s, 1H), 8.13 (d, J = 9.2 Hz, 1H), 7.52 (d, J = 9.2 Hz, 1H), 7.21 (dq, J = 14.3, 7.1 Hz, 3H), 6.98 (d, J = 6.9 Hz, 2H), 5.83 (s, 2H), 4.35 (q, J = 7.1 Hz, 2H), 3.41 (t, J = 7.6 Hz, 2H), 2.66 (t, J = 7.6 Hz, 2H), 1.34 (t, J = 7.1 Hz, 3H); 13C NMR (101 MHz, MeOD) δ 176.6, 162.8, 143.5, 142.1, 139.2, 129.7, 129.2, 128.4, 127.3, 127.1, 127.1, 121.2, 119.2, 112.5, 62.4, 49.6, 36.3, 21.9, 14.4; HRMS (ESI) calcd for C21H20N2O6 for [M+ H] 397.1394, found 397.1399.

**3-(1-benzyl-5-chloro-2-(ethoxycarbonyl)-4-(trifluoromethyl)-1H-indol- 3-yl)propanoic acid (13c)** TLC (EtAc/Hex40:60) Rf = 0.7; 10% yield; pale-white solid; 1H NMR (400 MHz, MeOD) δ 7.69 (d, J = 8.9 Hz, 1H), 7.41 (d, J = 9.1 Hz, 1H), 7.29 – 7.16 (m, 3H), 6.95 (d, J = 7.2 Hz, 2H), 5.79 (s, 2H), 4.35 (q, J = 7.0 Hz, 2H), 3.38 (t, 2H), 2.59 (t, 2H), 1.30 (t, J = 7.0 Hz, 3H). 13C NMR (101 MHz, MeOD) δ 176.8, 163.2, 141.9, 139.0, 139.0, 131.4, 129.7, 128.9, 128.5, 127.3, 127.1, 124.5, 124.0, 122.4, 117.5, 62.8, 49.6, 36.5, 22.5, 14.3; HRMS (ESI) calcd for C22H19ClF3NO4 for [M+ H] 454.1028, found 454.1036.

**3-(1-benzyl-5-chloro-2-(ethoxycarbonyl)-6-(trifluoromethyl)-1H-indol- 3-yl)propanoic acid (13c’)** TLC (EtAc/Hex40:60) Rf = 0.7; 40% yield; off-white solid; 1H NMR (400 MHz, Acetone-d6) δ 8.14 (s, 1H), 8.04 (s, 1H), 7.25 (dq, *J* = 14.3, 7.1 Hz, 3H), 7.05 (d, *J* = 7.3 Hz, 2H), 4.38 (q, *J* = 7.1 Hz, 2H), 3.42 (t, *J* = 7.6 Hz, 2H), 2.70 (t, *J* = 7.6 Hz, 2H), 1.35 (t, *J* = 7.1 Hz, 3H); 13C NMR (101 MHz, Acetone-d6) δ 174.0, 162.2, 139.1, 136.3, 130.4, 129.5, 128.1, 127.1, 126.0, 124.6, 123.8, 123.3, 122.8, 112.4, 112.3, 62.1, 49.0, 35.5, 21.3, 14.3.

**3-(1-benzyl-7-chloro-2-(ethoxycarbonyl)-5-(trifluoromethyl)-1H-indol- 3-yl)propanoic acid (13d)** TLC (EtAc/Hex40:60) Rf = 0.8; 25% yield; white solid; 1H NMR (400 MHz, MeOD) δ 8.09 (d, *J* = 0.8 Hz, 1H), 7.53 (d, *J* = 1.3 Hz, 1H), 7.27 – 7.13 (m, 3H), 6.82 (d, *J* = 6.8 Hz, 2H), 6.26 (s, 2H), 4.33 (q, *J* = 7.1 Hz, 2H), 3.39 (t, *J* = 7.6 Hz, 2H), 2.63 (t, *J* = 7.6 Hz, 2H), 1.31 (t, *J* = 7.1 Hz, 3H); 13C NMR (101 MHz, MeOD) δ 176.5, 162.7, 140.8, 130.6, 130.5, 129.5, 128.0, 126.5, 126.2, 124.6, 124.5, 124.3, 119.3, 118.7, 118.7, 62.6, 50.1, 36.2, 21.7, 14.3; HRMS (ESI) calcd for C22H19ClF3NO4 for [M+ H] 454.1028, found 454.1034.

## Acknowledgements

This research was supported by the United States-Israel Binational Science Foundation (BSF, Grant No. 2016142), the IMTI (TAMAT)/Israel Ministry of Industry–KAMIN Program, Grant No. 59081, and by the Israel Science Foundation (ISF, Grant No. 1023/18).

