## Supplementary material for "Dual Acting Small-Molecule Inhibitors Targeting Mycobacterial DNA Replication": Suplemental Figures and Tables

a.

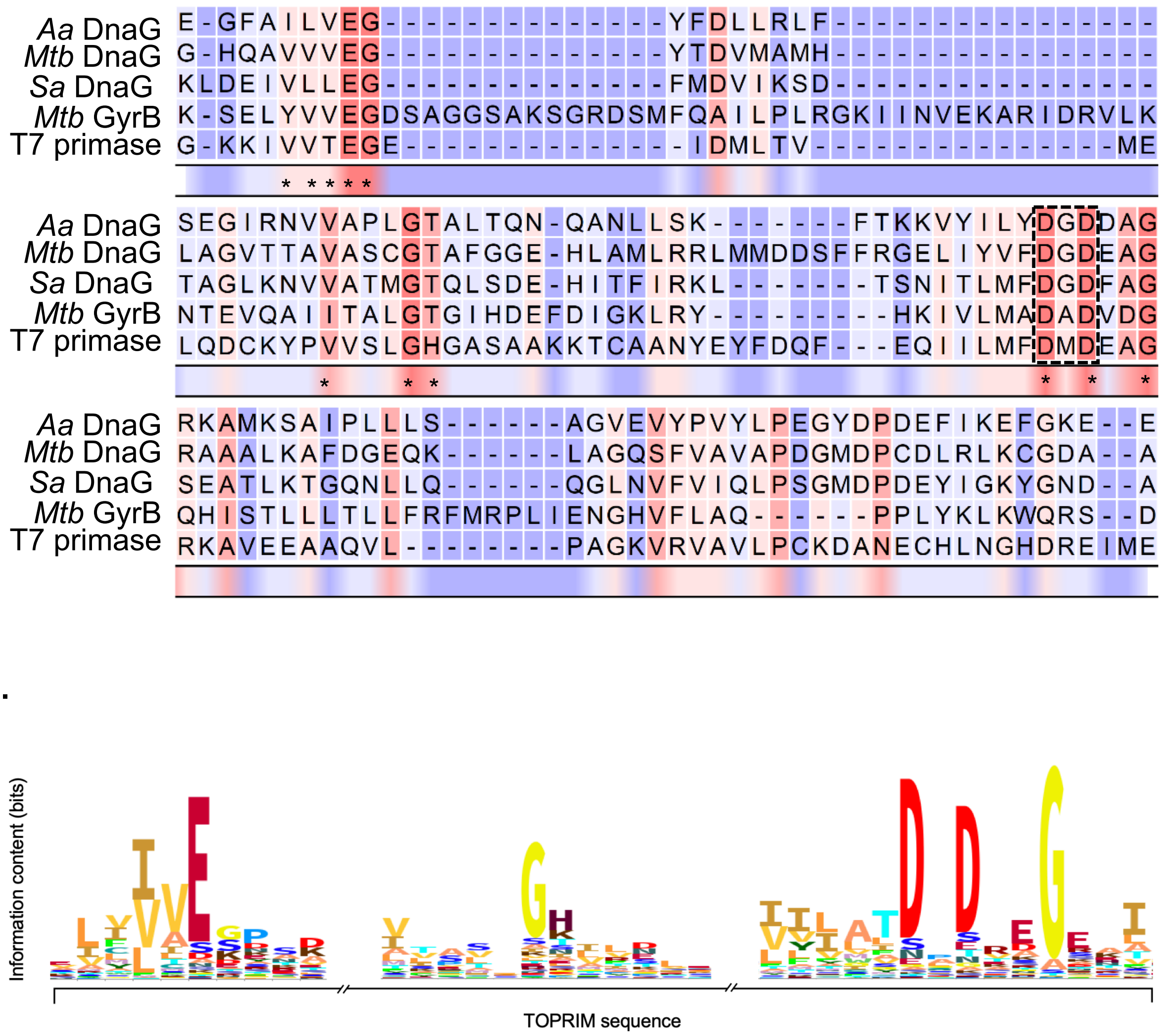

b.

**Figure S1. Amino acid sequence homology of DnaG-like primases.**
Multiple sequence alignment of bacterial DnaG primase [*Aa- Aquifex aeolicus*, *Mtb-Mycobacterium tuberculosis*, *Sa- Staphylococcus aureus*, T7- bacteriophage T7]. Amino acid residues are coloured from red for the most conserved amino acids to blue for the less conserved ones. b. Graphical representation of a sequence alignment using range of different sequences for TOPRIM fold. The height of each stack along selected parts of the TOPRIM sequence represent the relative frequency of an amino acid, and hence its conservation, in each position. All residues marked in asterisk in (a) appear at high frequency in the TOPRIM fold. Sequence logo used for graphical representation of the frequency of TOPRIM residues was created using the web interface: https://pfam.xfam.org/family/Toprim.

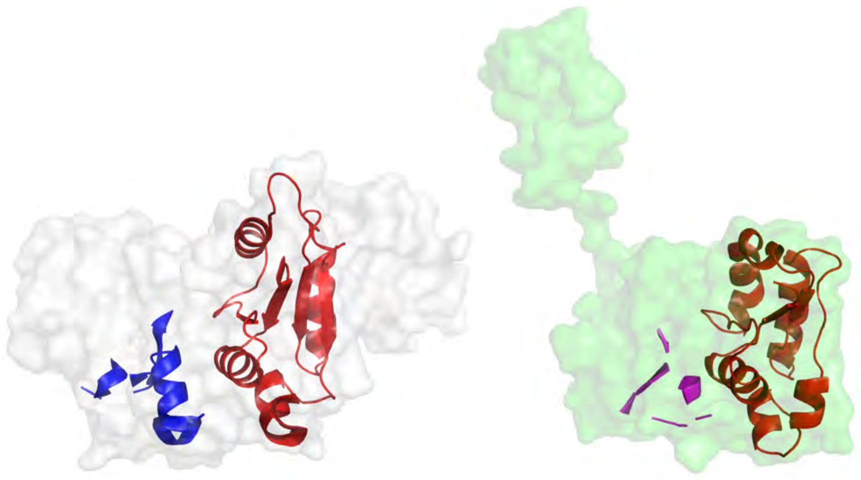

**Figure S2. Binding prediction of T7 DNA primase inhibitors within Mtb DnaG.**
Binding prediction of T7 DNA primase inhibitors within Mtb DnaG. Left: crystal structure of Mtb DnaG (PDB ID 5w33). Right: crystal structure of T7 DNA primase (PDB ID 1nui). TOPRIM fold is colored red; binding residues determined by NMR analysis are colored magenta. The predicted binding region, based on the structural alignment between the two enzymes, is colored blue. The figure was rendered in PyMOL (www.pymol.org).

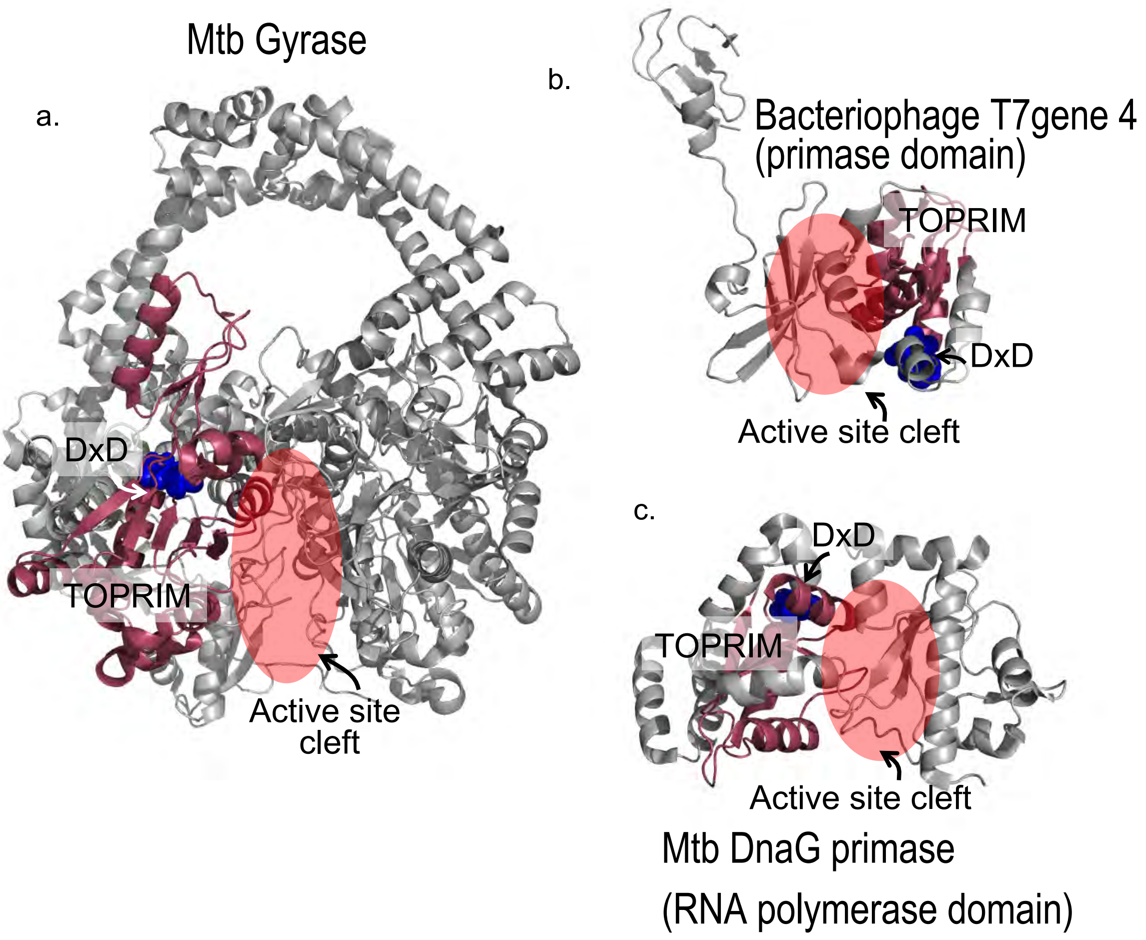

**Figure S3. Conserved structural motifs in DNA gyrase and DNA primase.**
DxD motif that coordinates the two magnesium ions is coloured blue and TOPRIM fold in pink. Active site cleft in all three enzymes is marked in red circles. (a) *Mtb* GyrB (PDBID: 5BS8), DxD motif D532-D534, active site residues of the catalytic core of GyrB: E459, T448-V675. TOPRIM residues (T448-V675). (b) Bacteriophage T7 gene 4 primase domain (PDBID: 1NUI) TOPRIM 151-228, DxD motif D207-D209. (c) *Mtb* DnaG (PDBID: 5W33), DxD motif D319-D321, active site catalytic residues (E268, G112-W425). TOPRIM residues (Q263-V346). The figure was created using PyMOL (www.pymol.org).

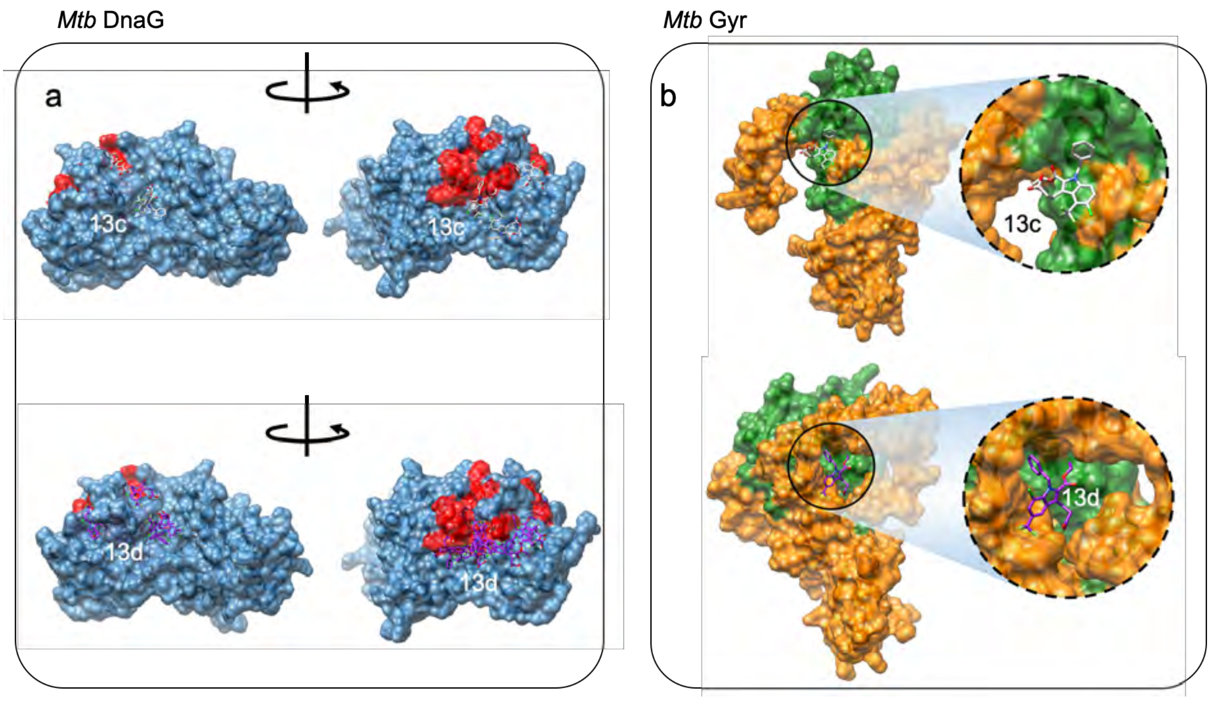

**Figure S4. Docking of small molecule inhibitors to available crystal structures of *Mtb* DnaG primase and *Mtb* Gyr.** a. Structural representation of molecular docking results performed by Autodock (http://autodock.scripps.edu/), for DNA primase (RNA polymerase domain, marked in blue) from *Mtb* (PDB entry: 5W33) and its putative inhibitors 13c (top, marked in light grey) and 13d (bottom, marked in purple). Pairwise structural alignment of the protein with its homolog from T7 bacteriophage (PDB entry: 1NUI), which was experimentally tested *via* NMR for the binding of other potential inhibitors (Ilic et al. 2016), suggests specific residues (shown in red) hypothetically involved in the binding of 13c to *Mtb* DNA primase. The results show numerous conformations of 13c, docked in the overt clefts of the protein where the predicted binding residues are situated, thus conform well to the experimentally based evidences. b. Structural representation of molecular docking results performed by Autodock, for the B subunit of DNA gyrase (surface representation, marked in orange) from *Mycobacterium tuberculosis* (PDB entry: 5BS8) and its putative inhibitors: 13c (top, marked in light grey) and 13d (bottom, marked in purple). Both molecules, 13c and 13d are, bound in the C-terminal cleft of the gyrase, in proximity to the TOPRIM domain (marked in green). Figures were created using Chimera software for molecular visualization (https://www.cgl.ucsf.edu/chimera/)

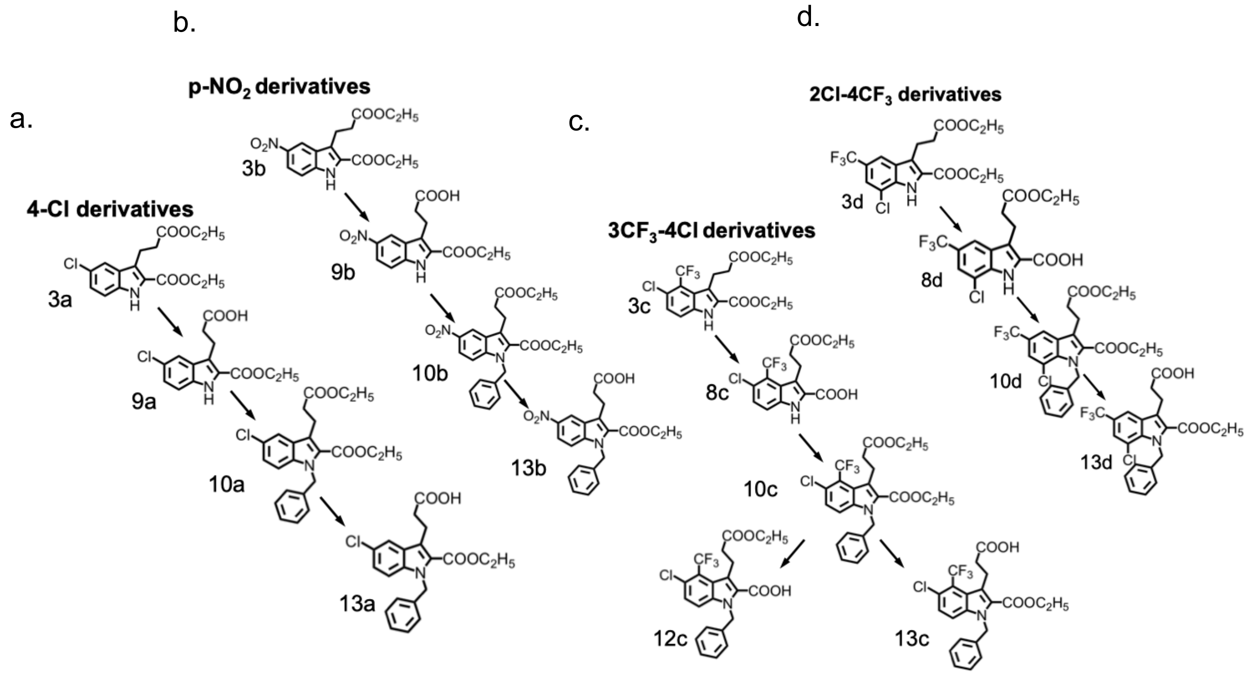

**Figure S5. Schematic representation of optimization process of four different aniline derivatives.**

Every derivative (a-d) underwent three rounds of optimization yielding five best analogs belonging to group III.

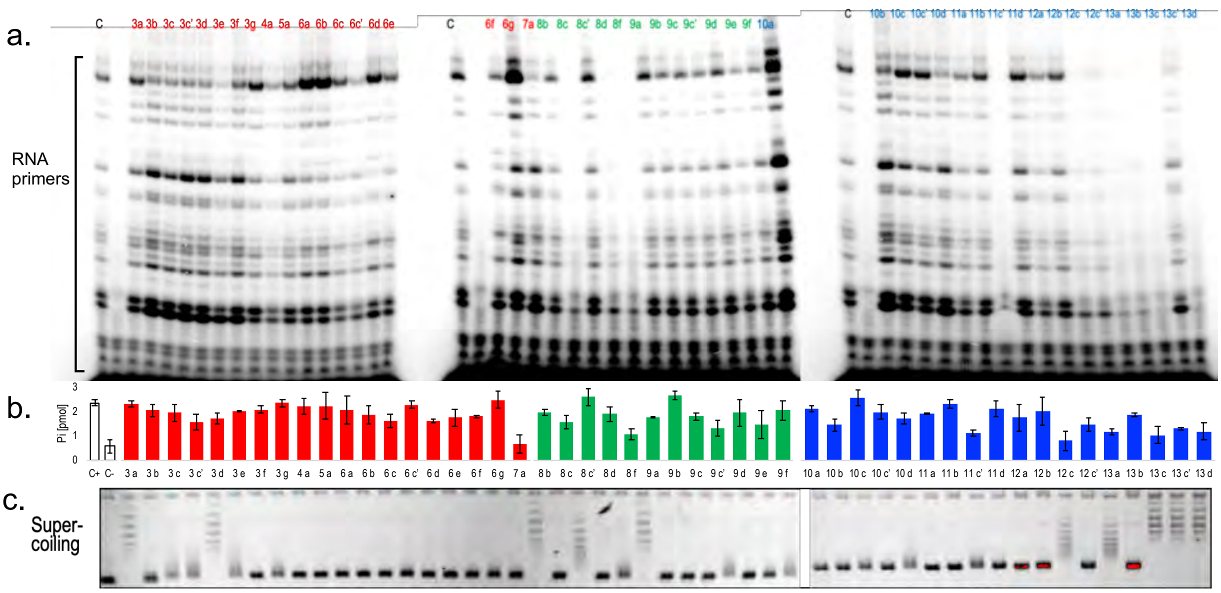

**Figure S6. Effect of newly synthesized small molecules on RNA primers synthesis catalyzed by DnaG primase and GyrA/B from *Mtb*.**

A. Effect of small molecules on the activity of DnaG primase. The reaction contained 50 μM oligonucleotide 5'- CCGACCCGTCCGTAATACAGAGGTAATTGTCACGGT-3', 250 μM CTP, GTP, UTP, α-$^{32}$P-ATP in a standard reaction mixture, and 4 mM of each small molecule. The reaction was initiated by adding *Mtb* DnaG primase to a final concentration of 500 nM. After incubation, the radioactive products were analyzed by electrophoresis through a 25% polyacrylamide gel containing 7 M urea, and visualized using autoradiography. B. Effect of newly synthesized small molecules on ATP hydrolysis activity by GyrB of *Mtb*. C. Effect of newly synthesized small molecules on supercoiling activity by GyrA of *Mtb*. The experiments in (B) and (C) were performed as described in Fig. 2b.

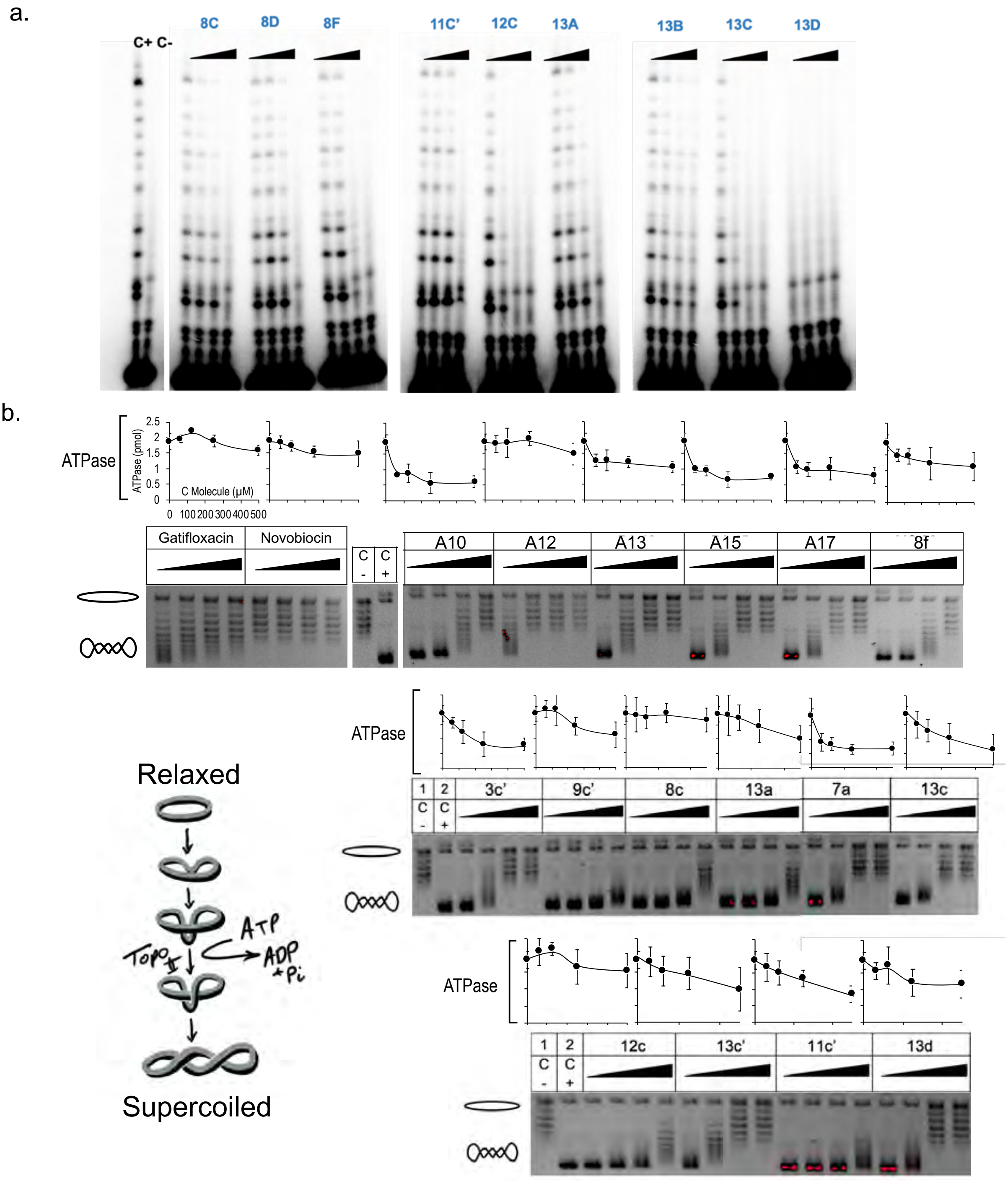

**Figure S7. Dose response of selected small molecule inhibitors on the activities of DnaG primase and GyrA/B from *Mtb*.**

A. Inhibition of *Mtb* DnaG activity. The reaction contained 200 nM *Mtb* DnaG, oligonucleotide (5'-CCGACCCGTCCGTAATACAGAGGTAATTGTCACGGT-3'), [α-$^{32}$P]–ATP, CTP, GTP, UTP and synthesized compounds in increasing concentrations of 0.5, 1.0, 2.0 and 4.0 mM. After incubation of 1hr, the radioactive products were separated by electrophoresis through a 25% polyacrylamide gel containing 7 M urea and visualized by autoradiography. Inhibition of primase activity is characterized by the decrease in RNA primer formation, observed as the decrease of radioactive signal. B. Dose response of selected small molecule inhibitors on Gyr supercoiling activity (gel) and ATPase activity (graph). The molecules were tested in concentration of 62.5 µM, 125µM, 250µM and 500µM. Gatifloxacin and Novobiocin were used as controls.

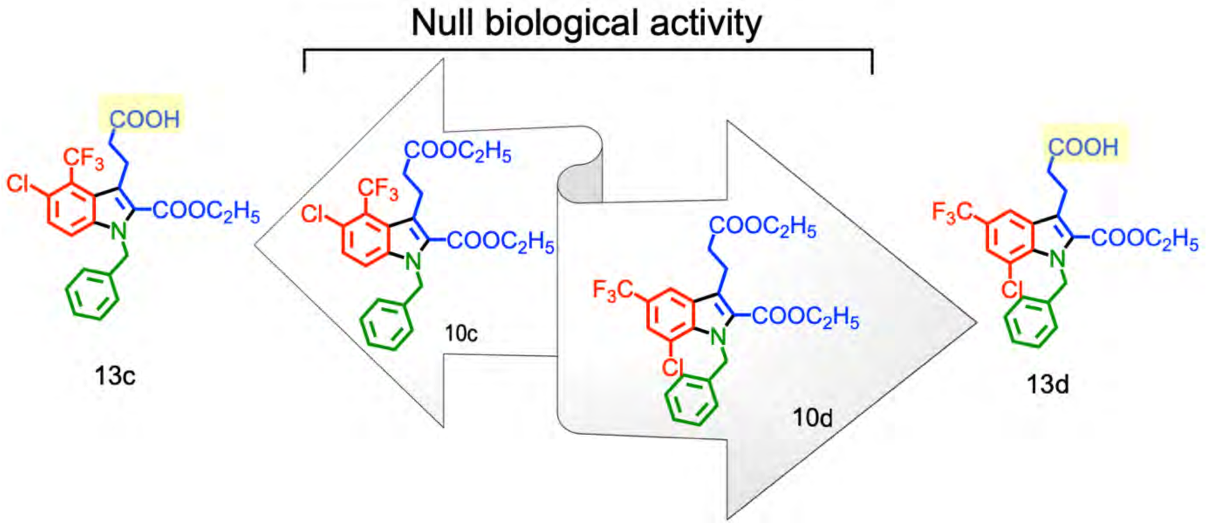

**Figure S8. Summary of binding-activity cliff studies on compounds 10c and 10d and their corresponding analogues 13c and 13d.**
Colour coding is used to highlight functional regions of the lead compounds. Changes in region A of compound 10c and 10d are shown in blue, changes in region B are shown in red, and changes in region C are shown in green. a decrease in potency of molecules 6a-g, 6c', 4a, 5a, and 7a was attributed to the absence of carboxyethyl and propanoic acid groups at position 2 and 3, respectively. In addition, introduction of a benzyl group in -NH position of indole ring increased potency (as noted for the resulting Group III series of molecules).

**Figure S9-S83. Characterization of synthetic small-molecules.**

a.

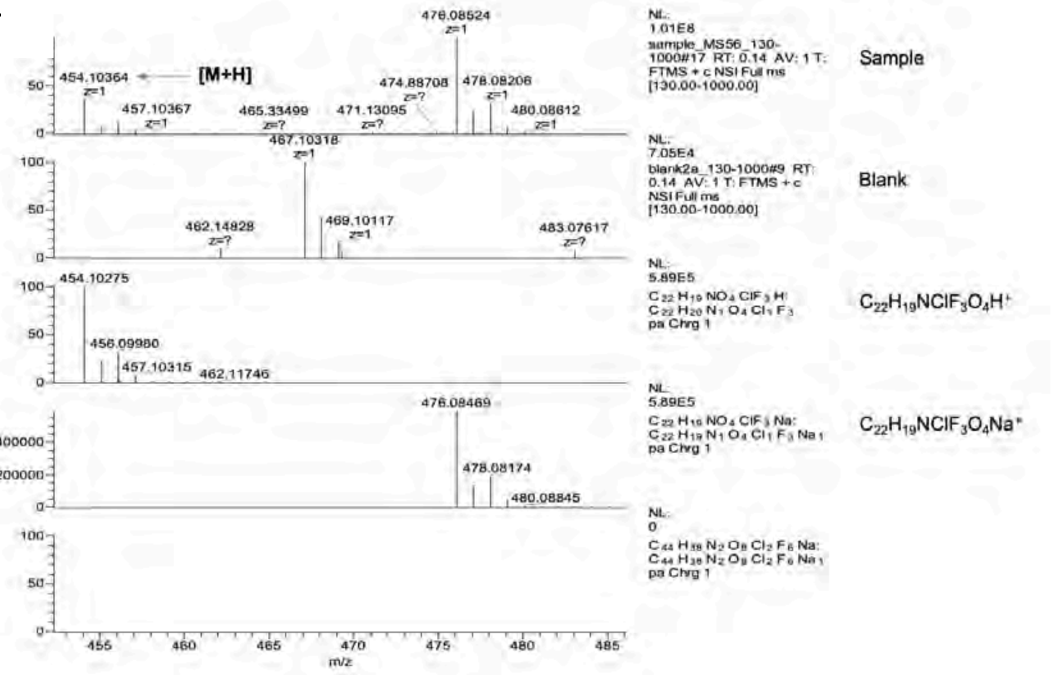

b.

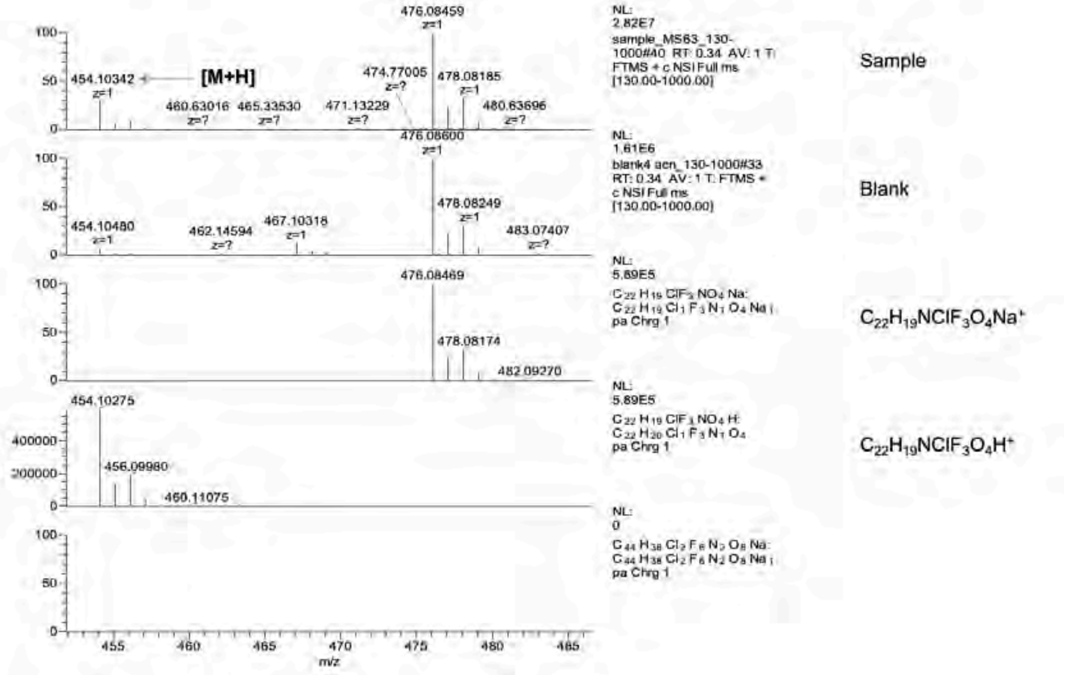

**Figure S9. High Resolution Mass Spectrometry (HRMS) in positive ion mode of compound 13c (a) compound 13d (b).**

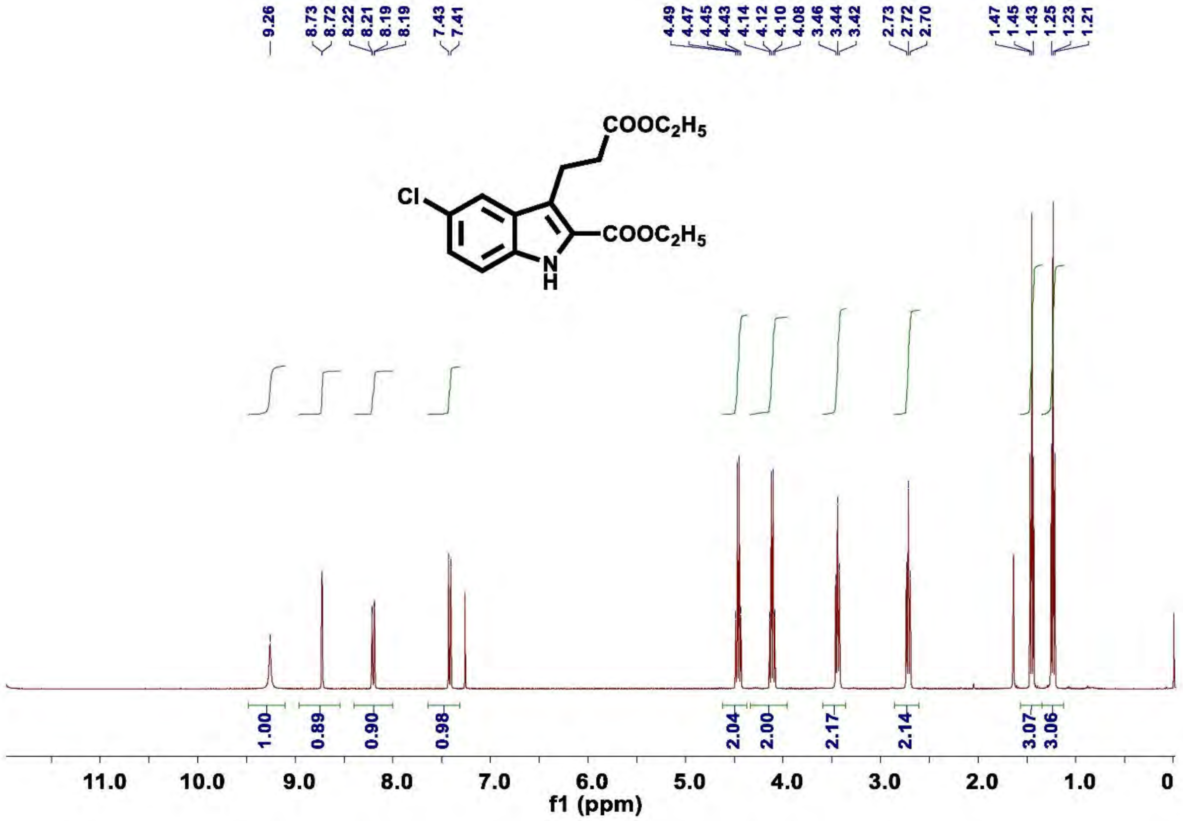

**Figure S10.** $^{1}$**H NMR of compound 3a**

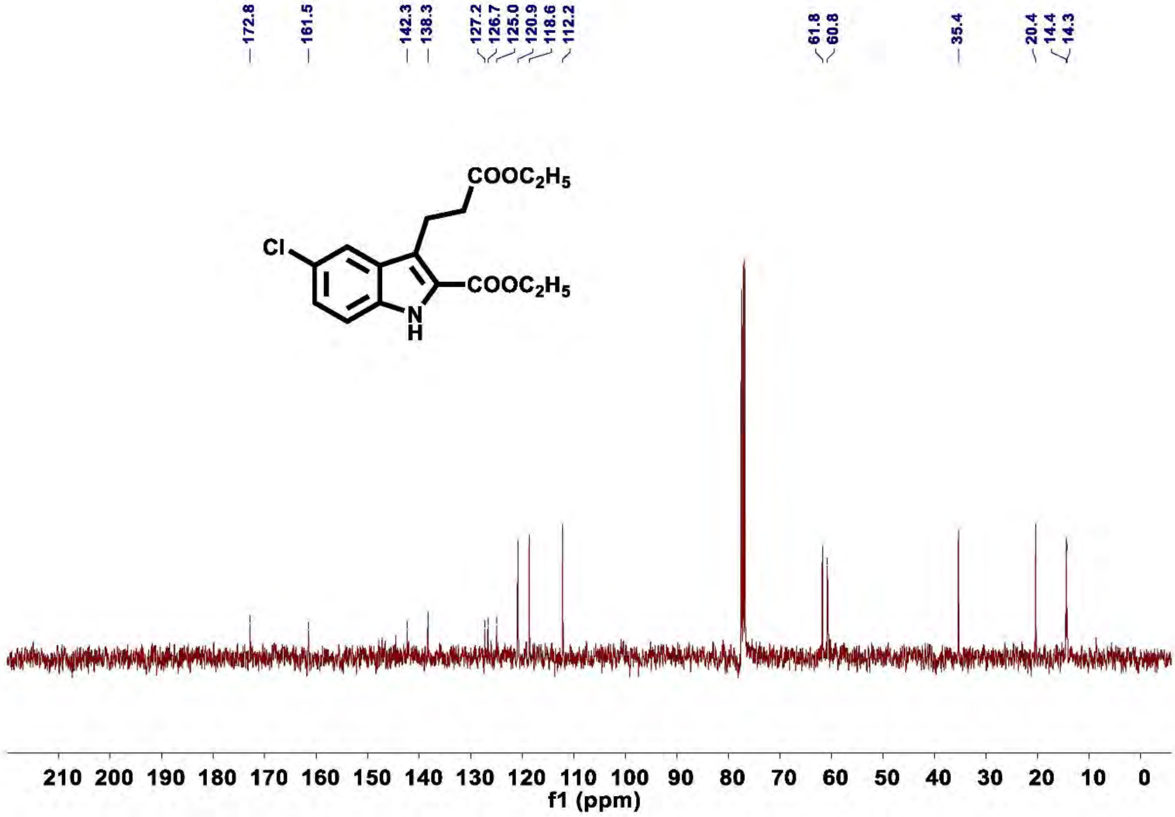

**Figure S11.** $^{13}$**C NMR of compound 3a**

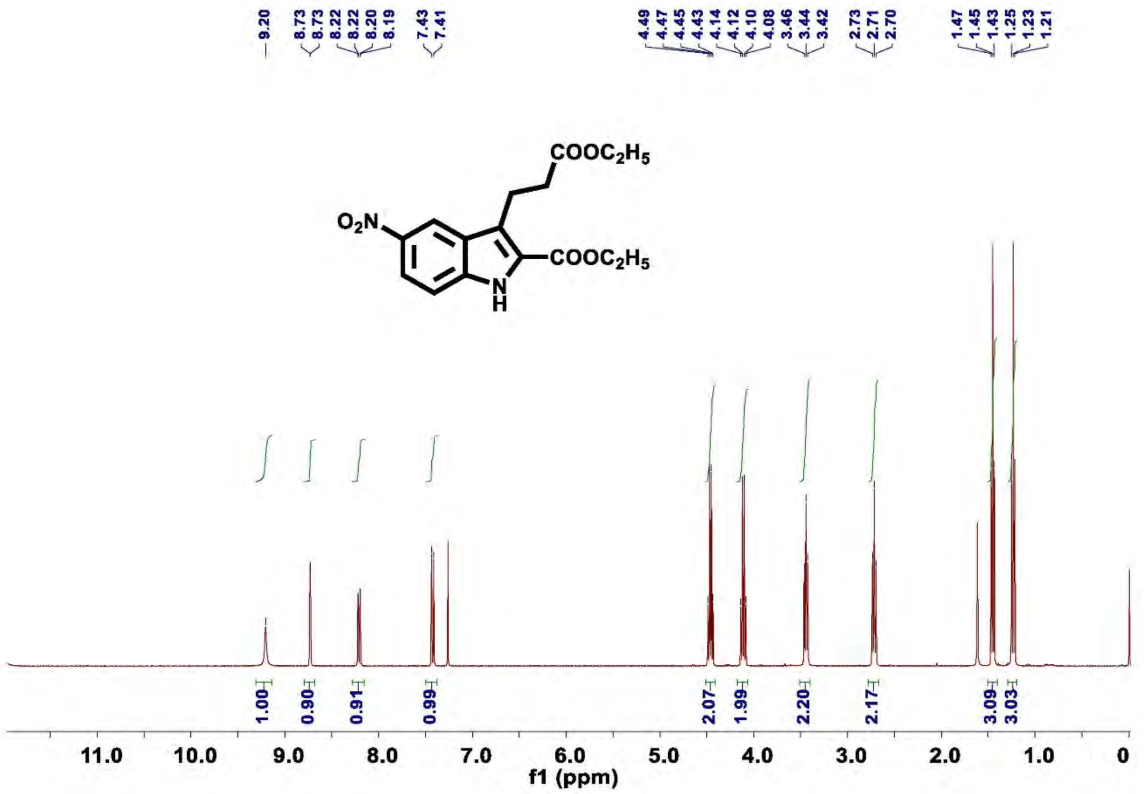

**Figure S12. $^1$H NMR of compound 3b**

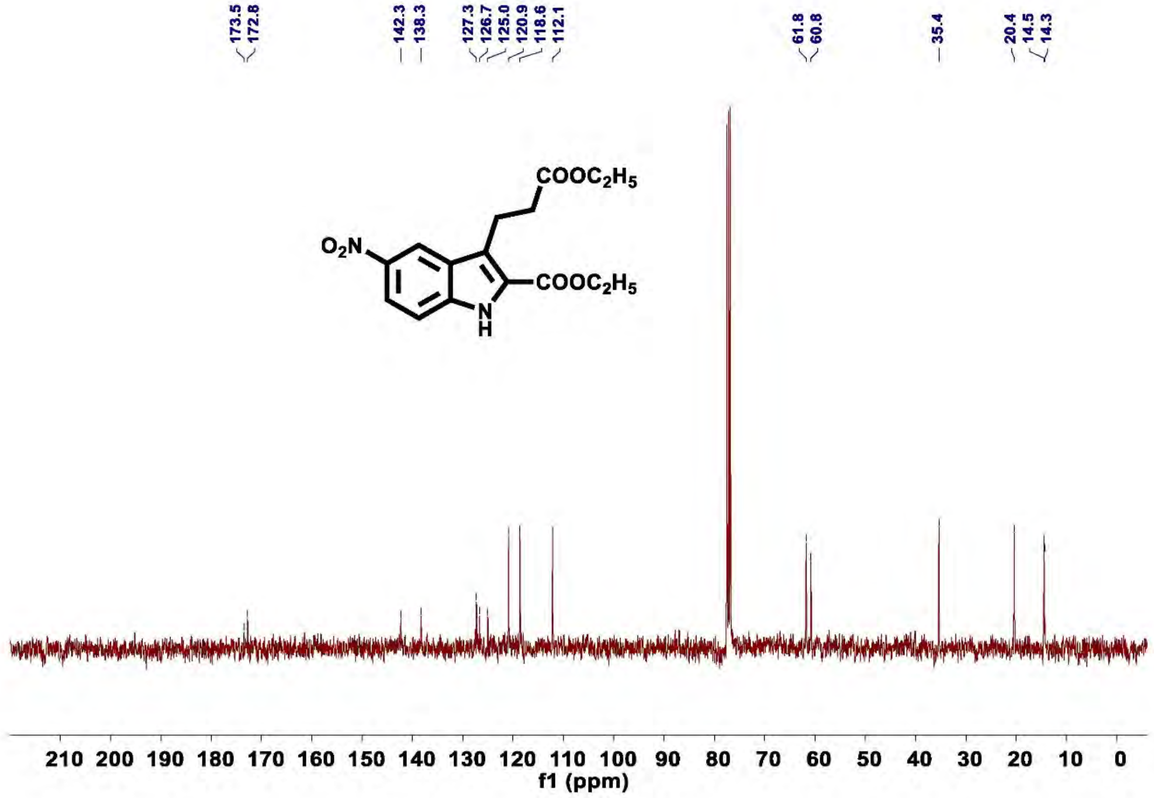

**Figure S13.** $^{13}$**C NMR of compound 3b**

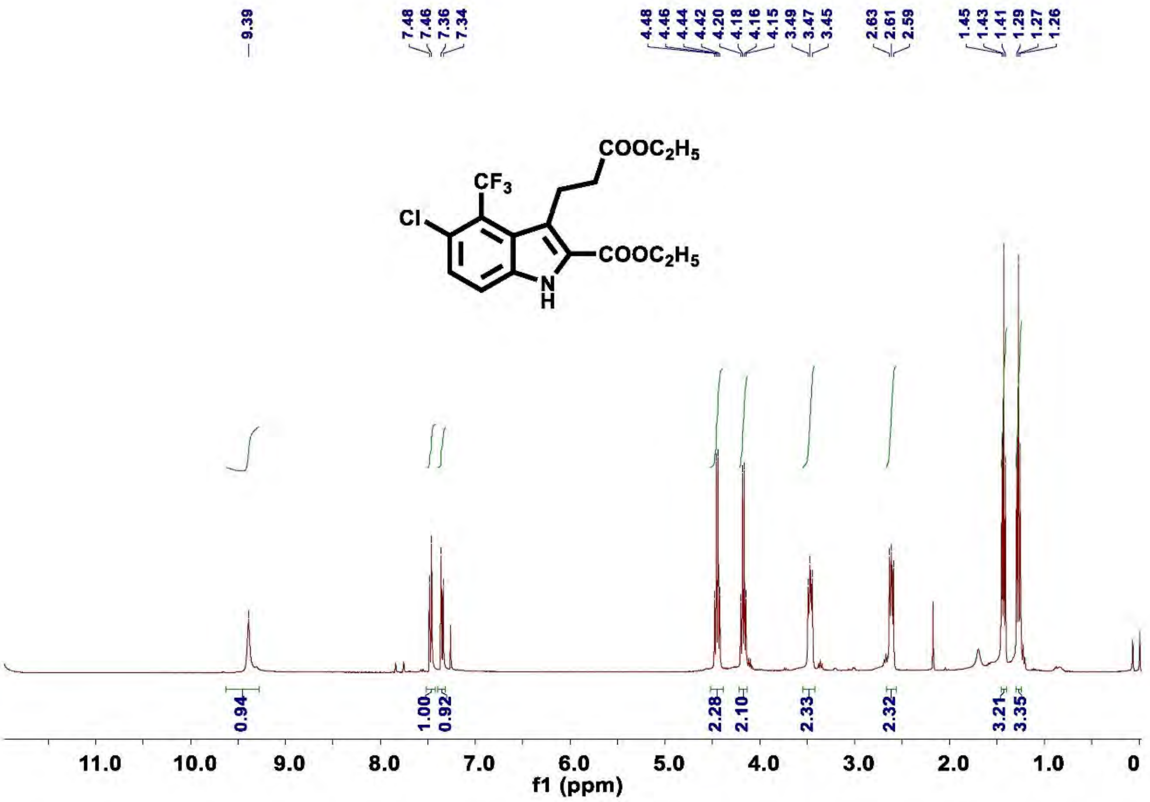

**Figure S14.** $^{1}$**H NMR of compound 3c**

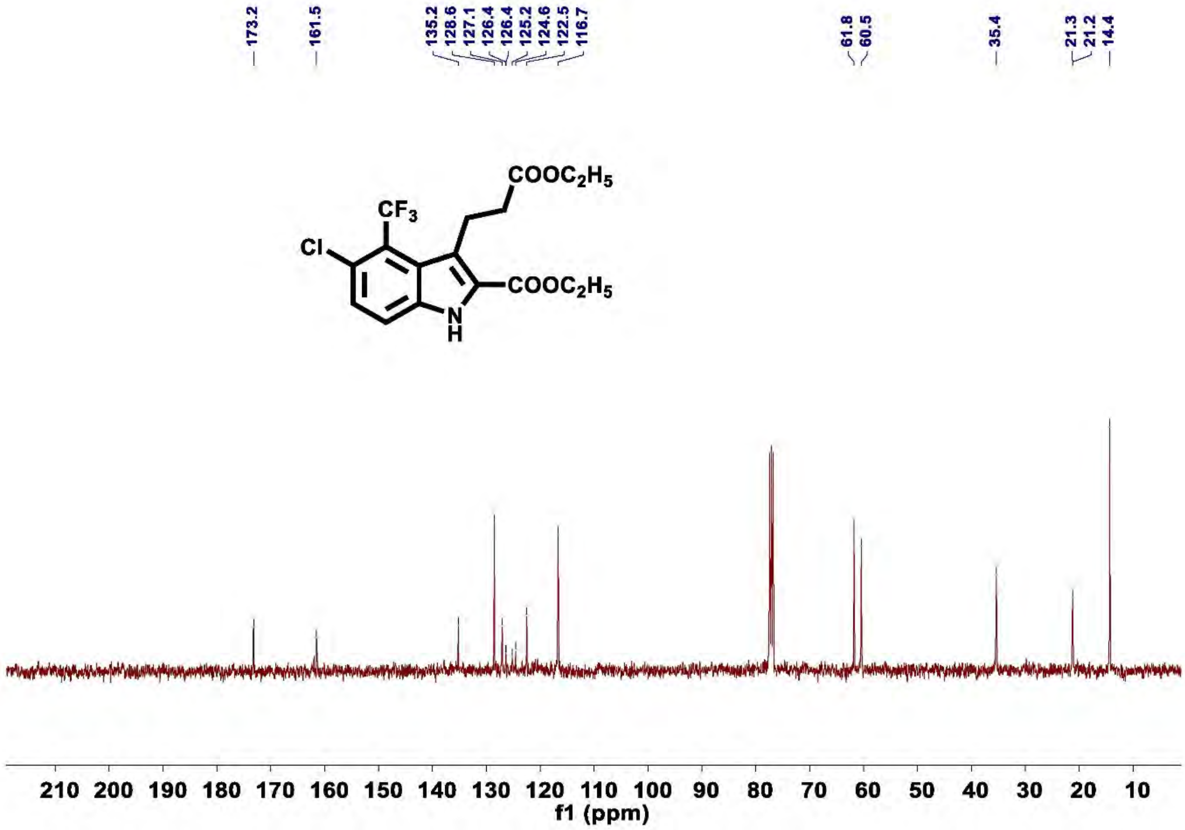

Figure S15. $^{13}$C NMR of compound 3c

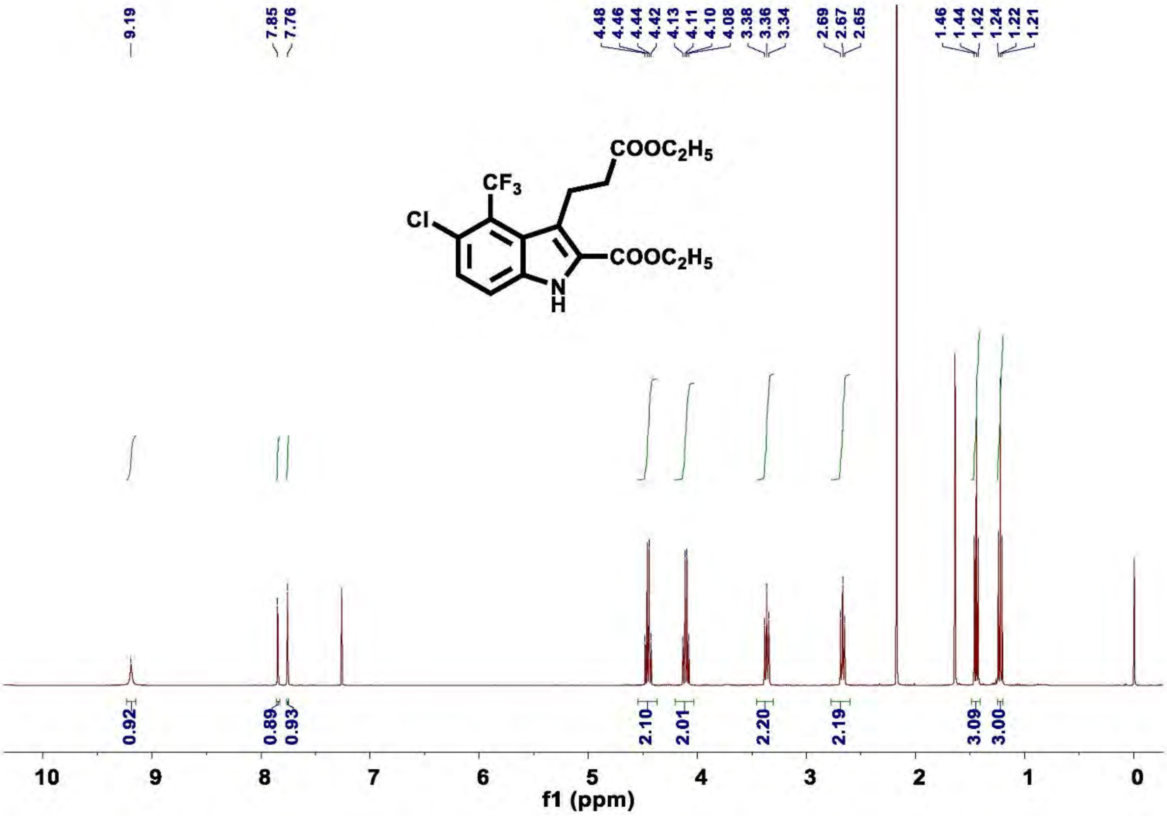

Figure S16. $^{1}$H NMR of compound 3c’

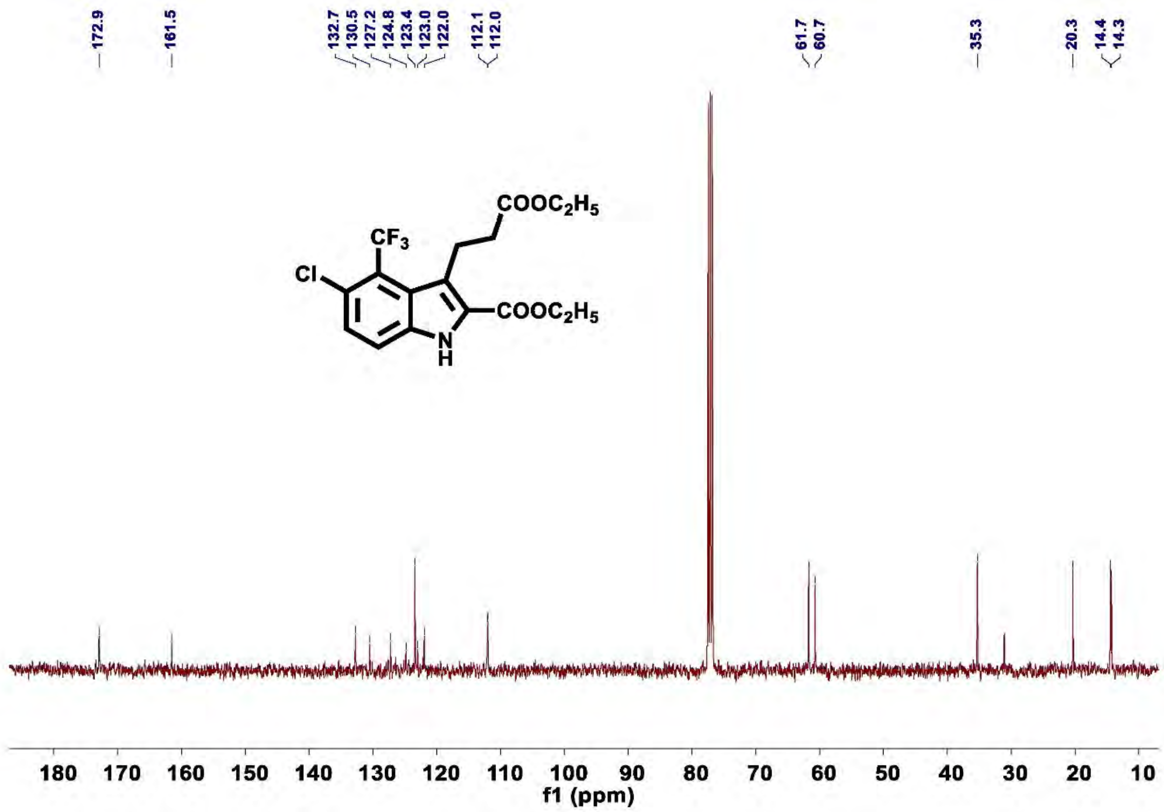

**Figure S17.** $^{13}$**C NMR of compound 3c'**

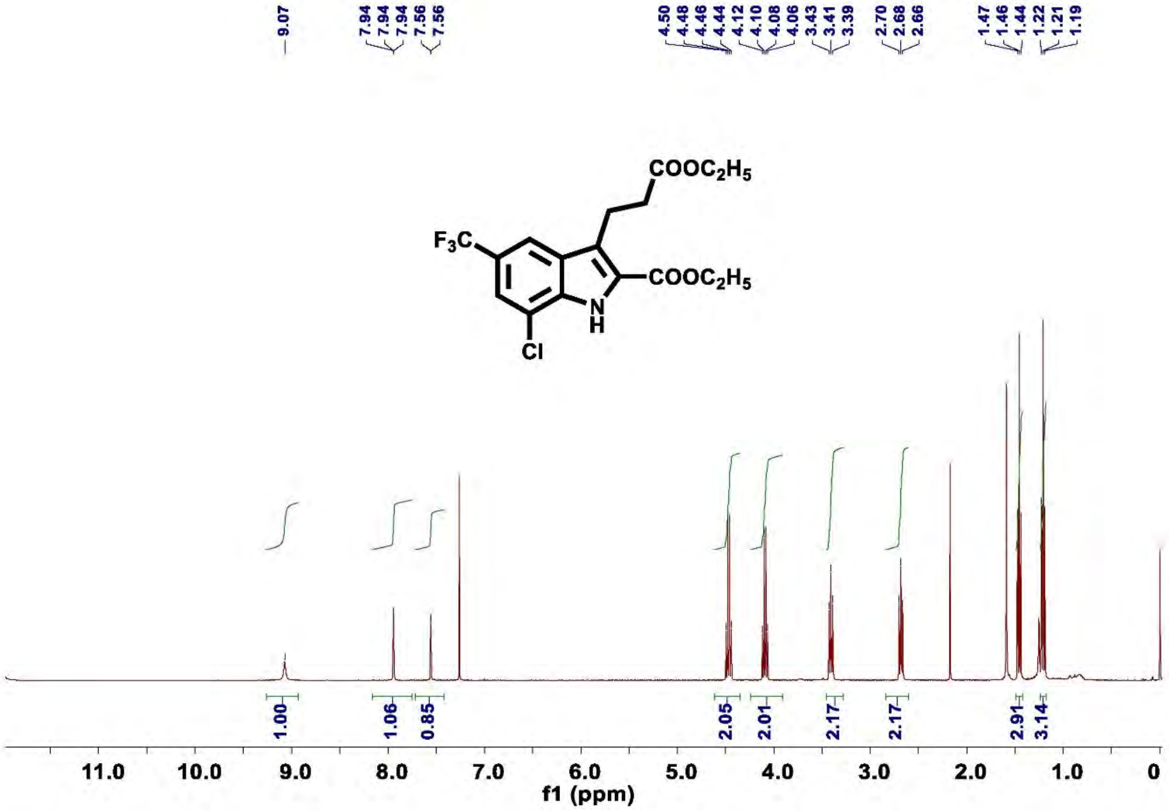

**Figure S18.** $^{1}$**H NMR of compound 3d**

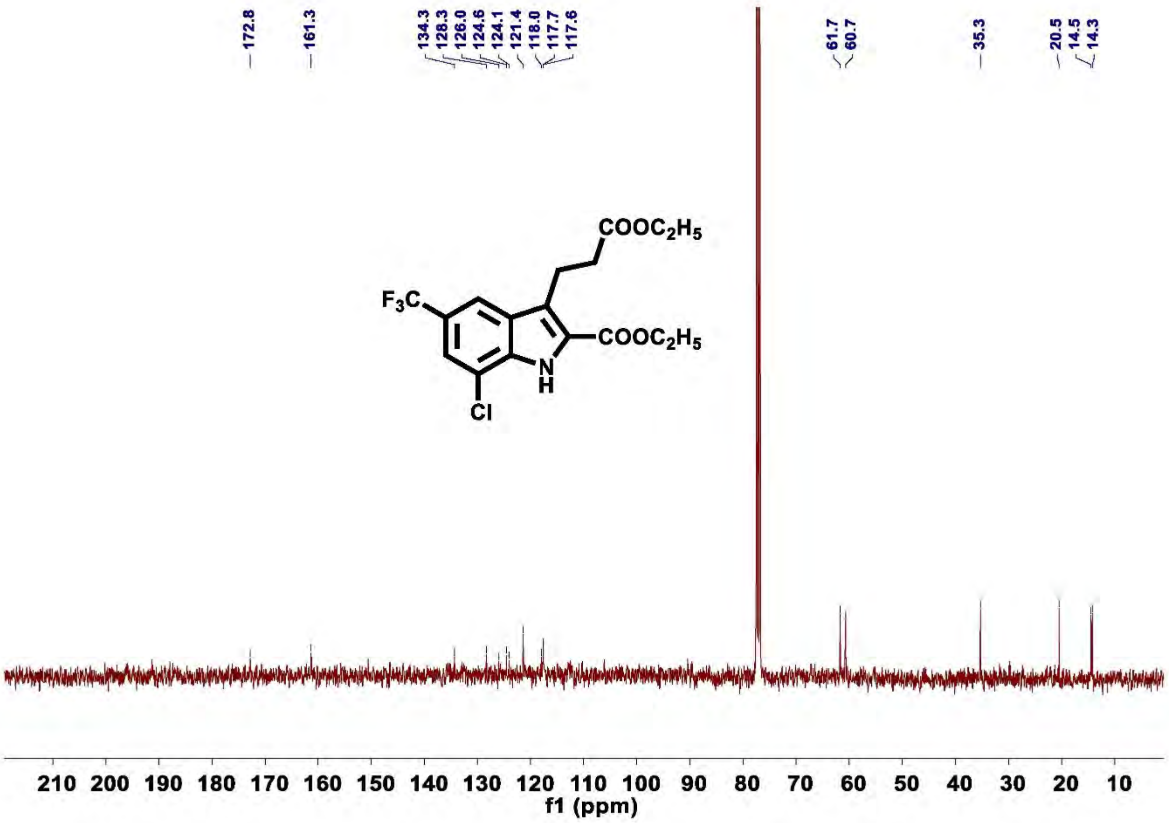

Figure S19. $^{13}$C NMR of compound 3d

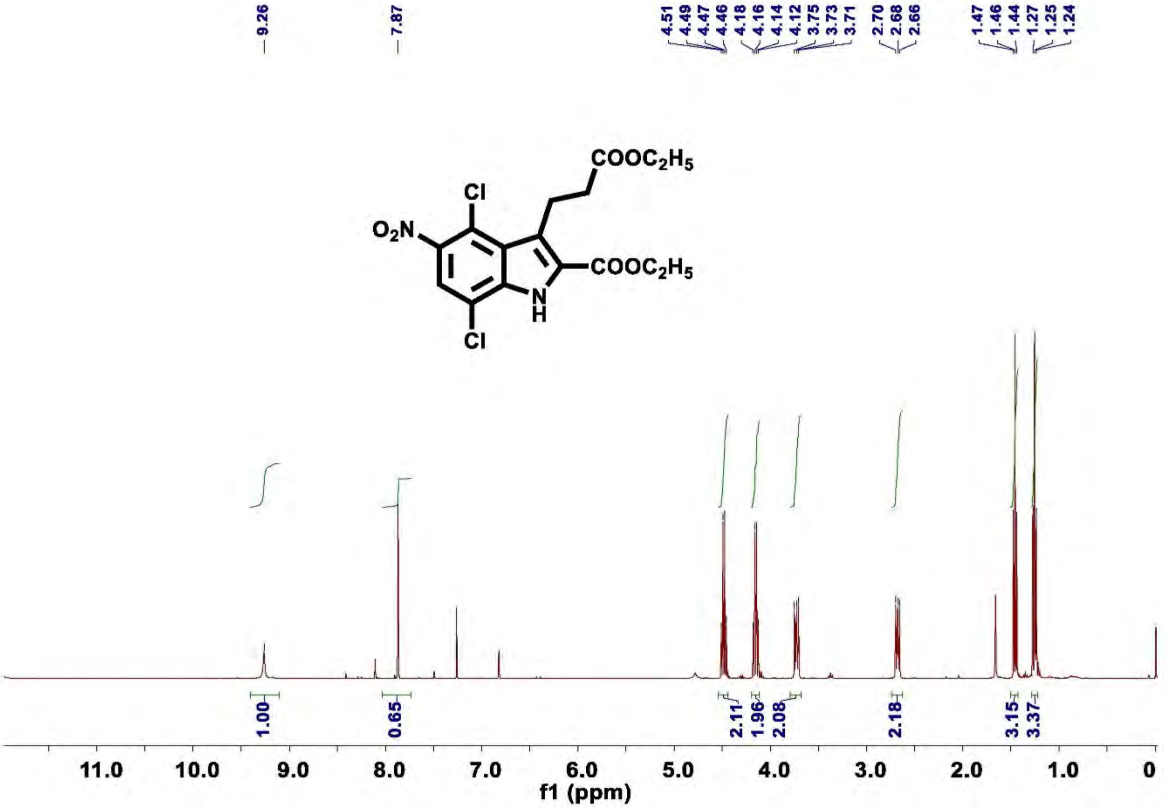

Figure S20. $^{1}$H NMR of compound 3e

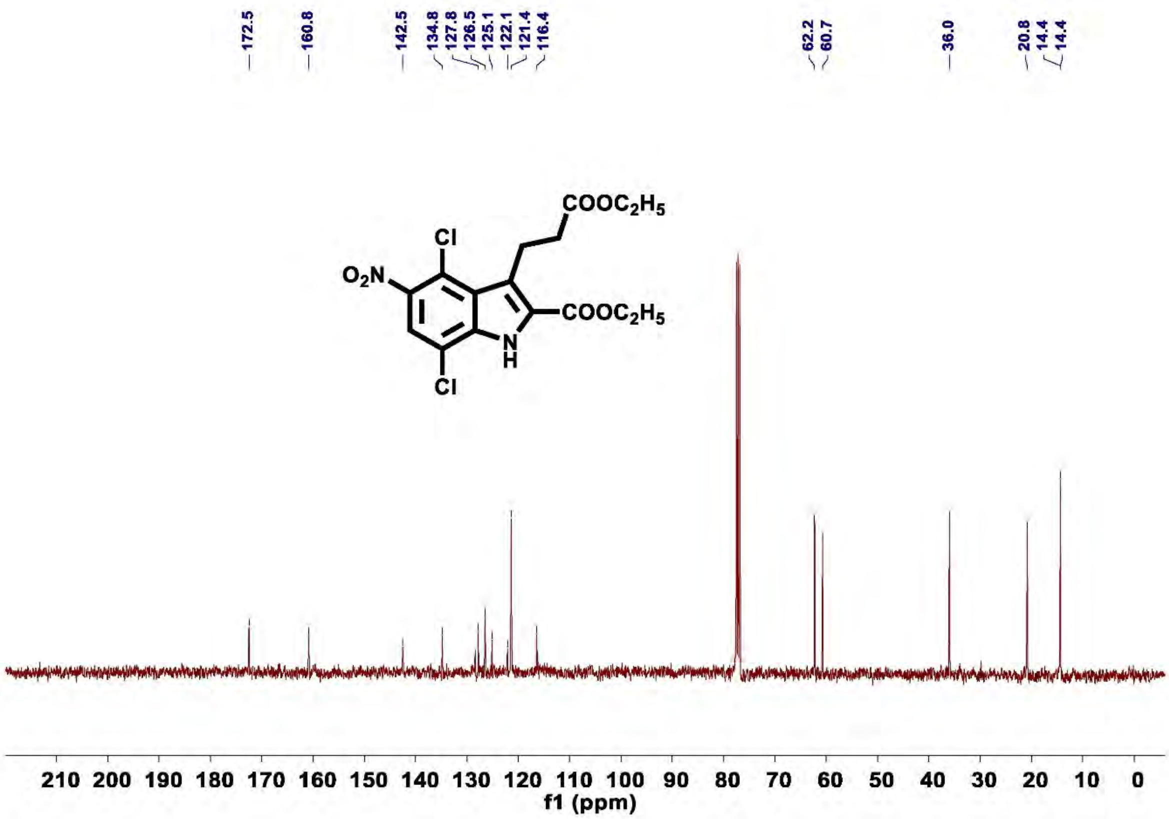

Figure S21. $^{13}$C NMR of compound 3e

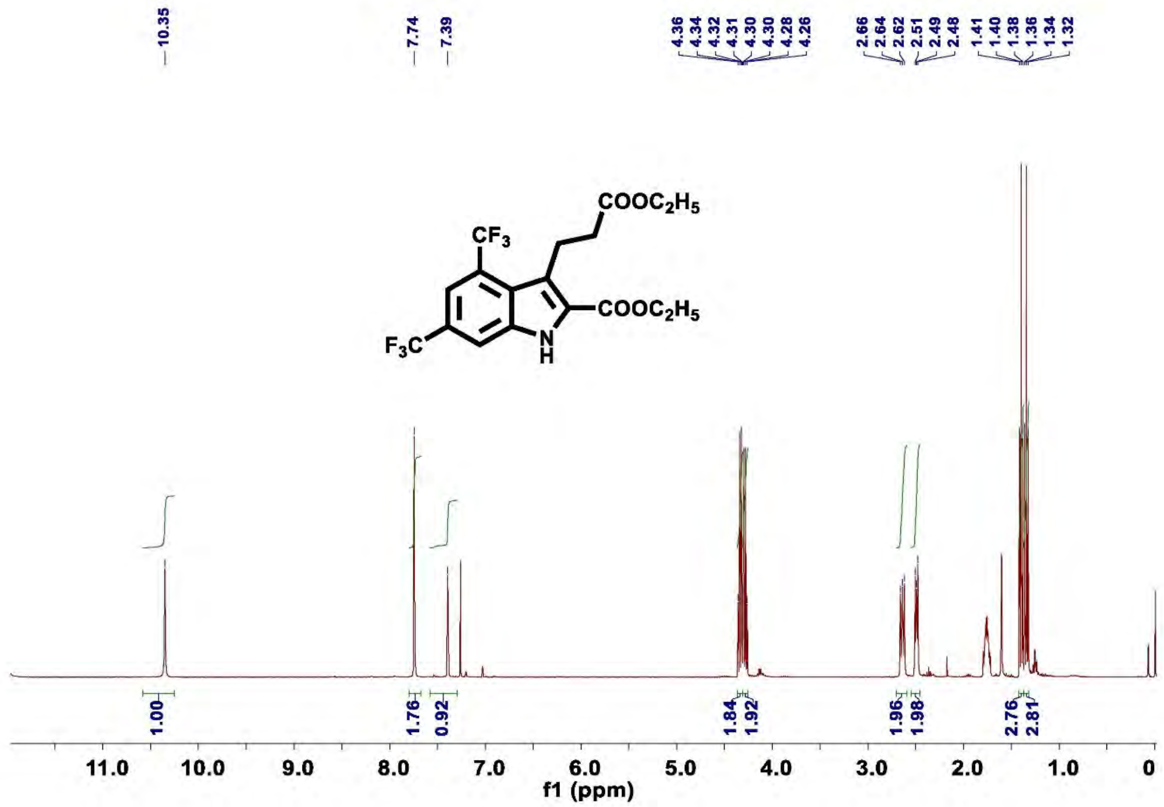

Figure S22. $^{1}$H NMR of compound 3f

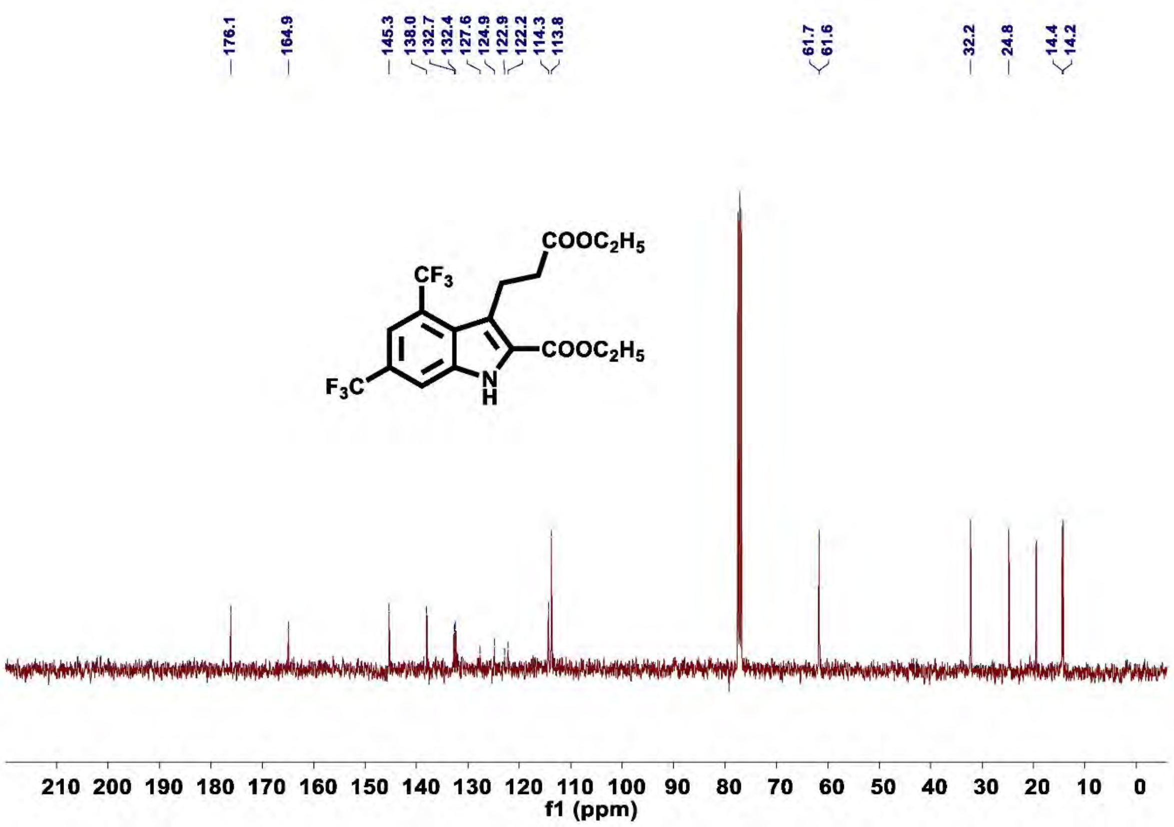

**Figure S23.** $^{13}$**C NMR of compound 3f**

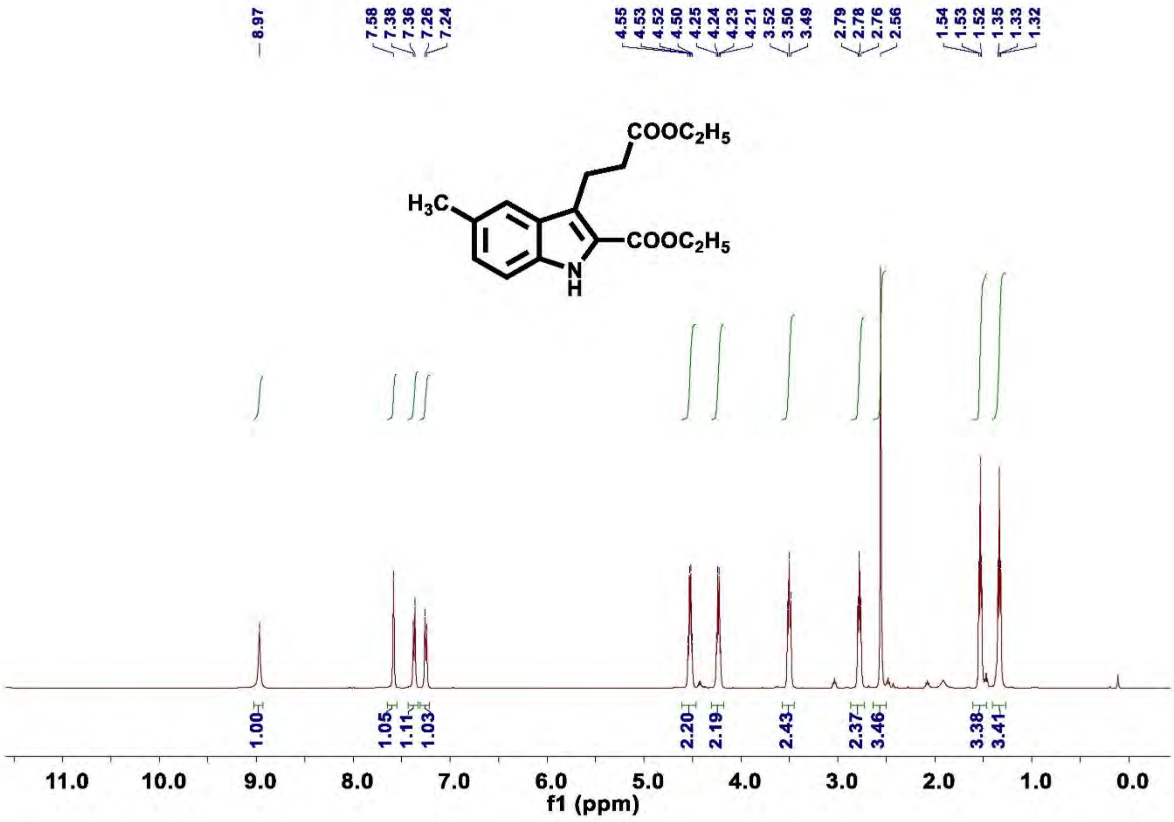

**Figure S24.** $^{1}$**H NMR of compound 3g**

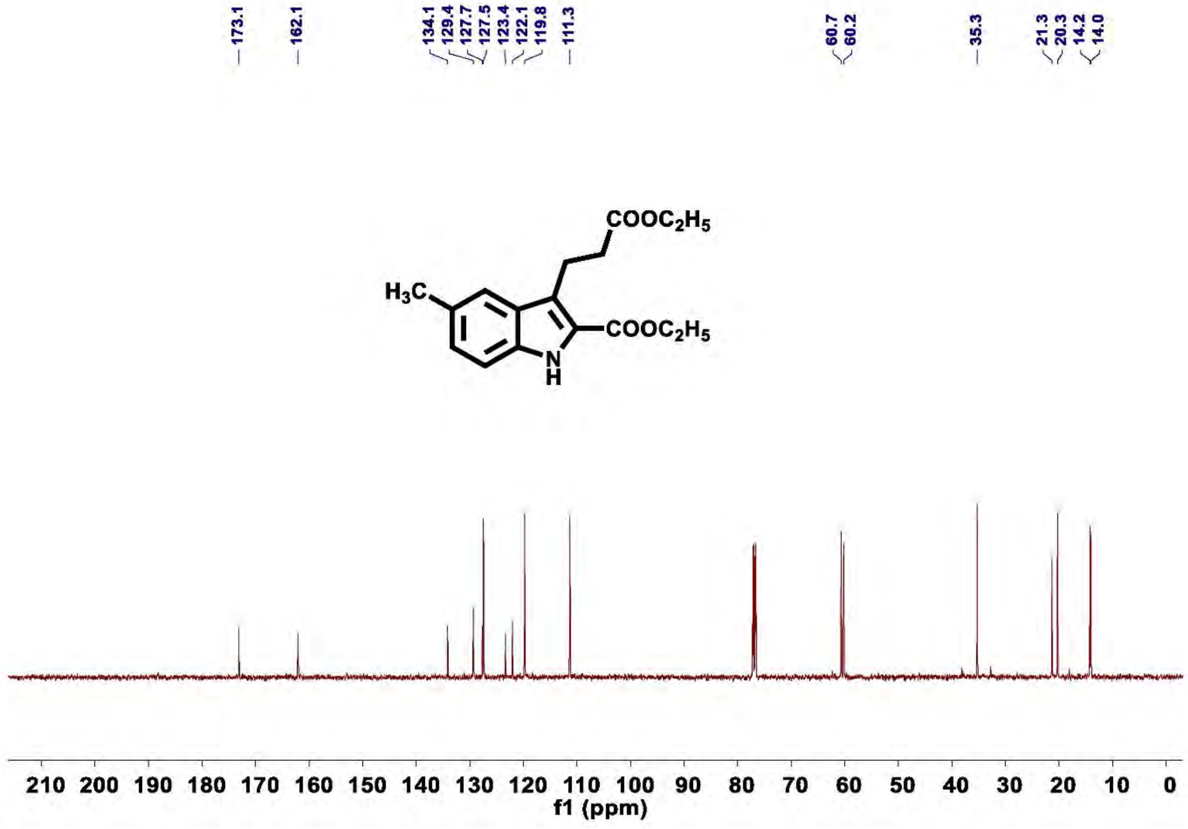

**Figure S25.** $^{13}$**C NMR of compound 3g**

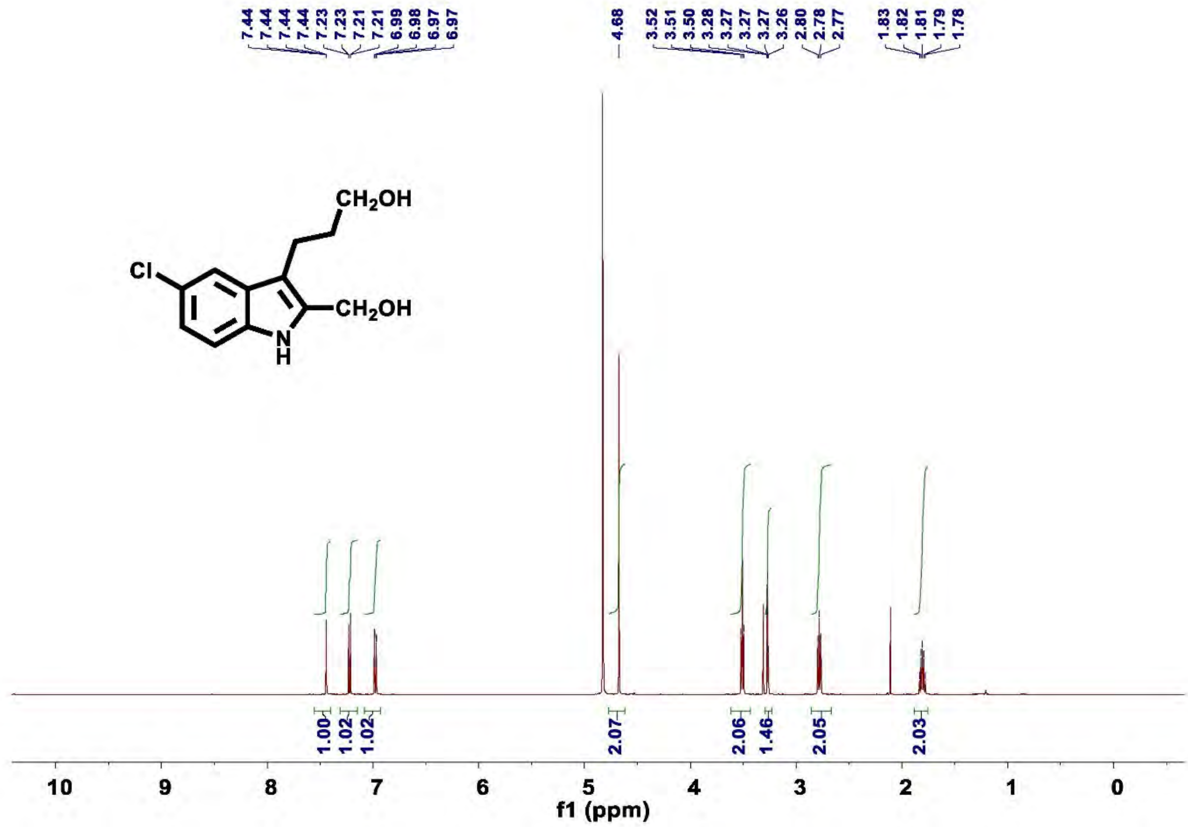

**Figure S26.** $^{1}$**H NMR of compound 4a**

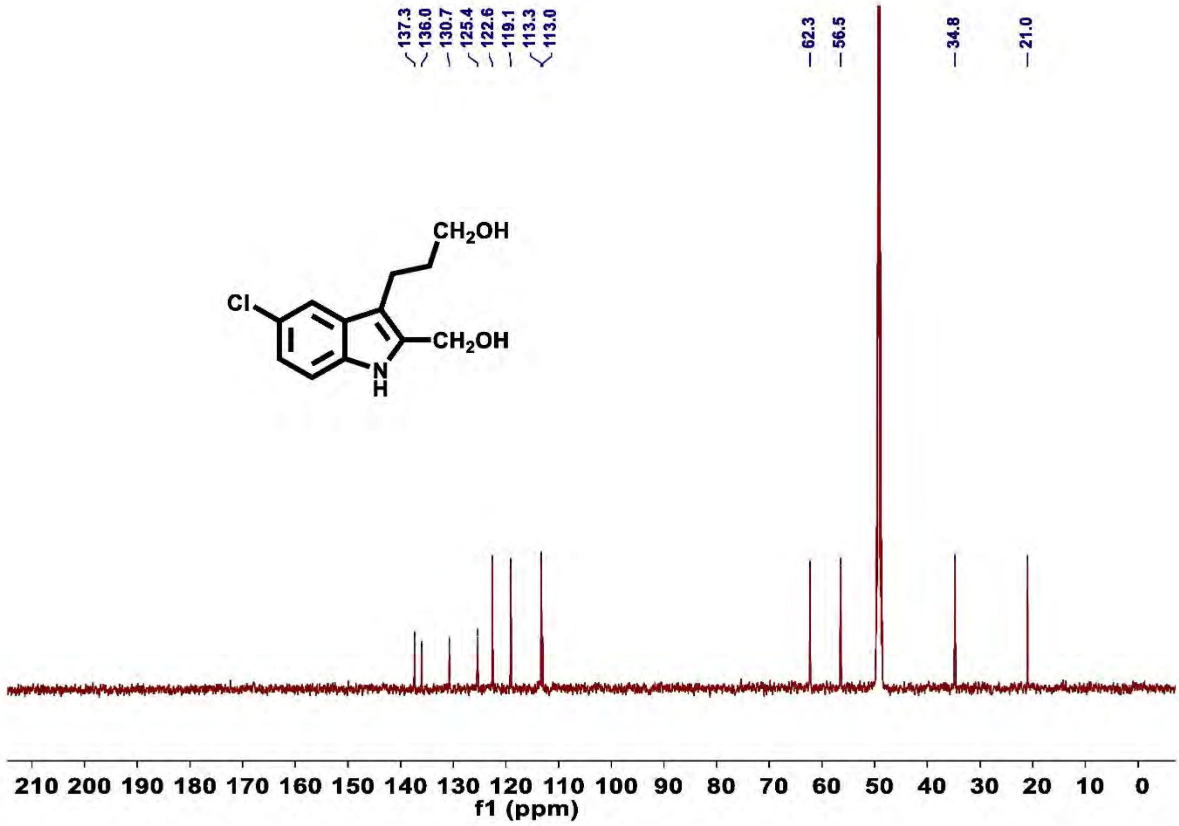

**Figure S27.** $^{13}C$ **NMR of compound 4a**

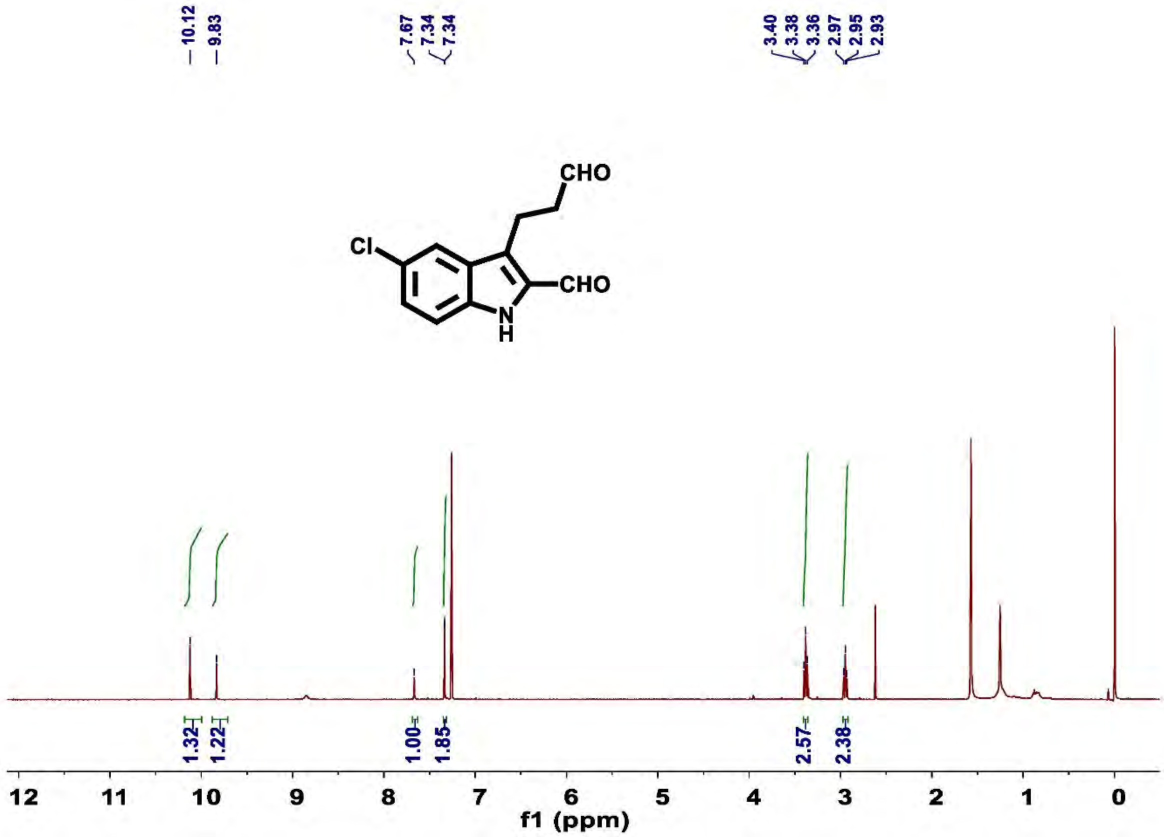

Figure S28. $^{1}H$ NMR of compound 5a

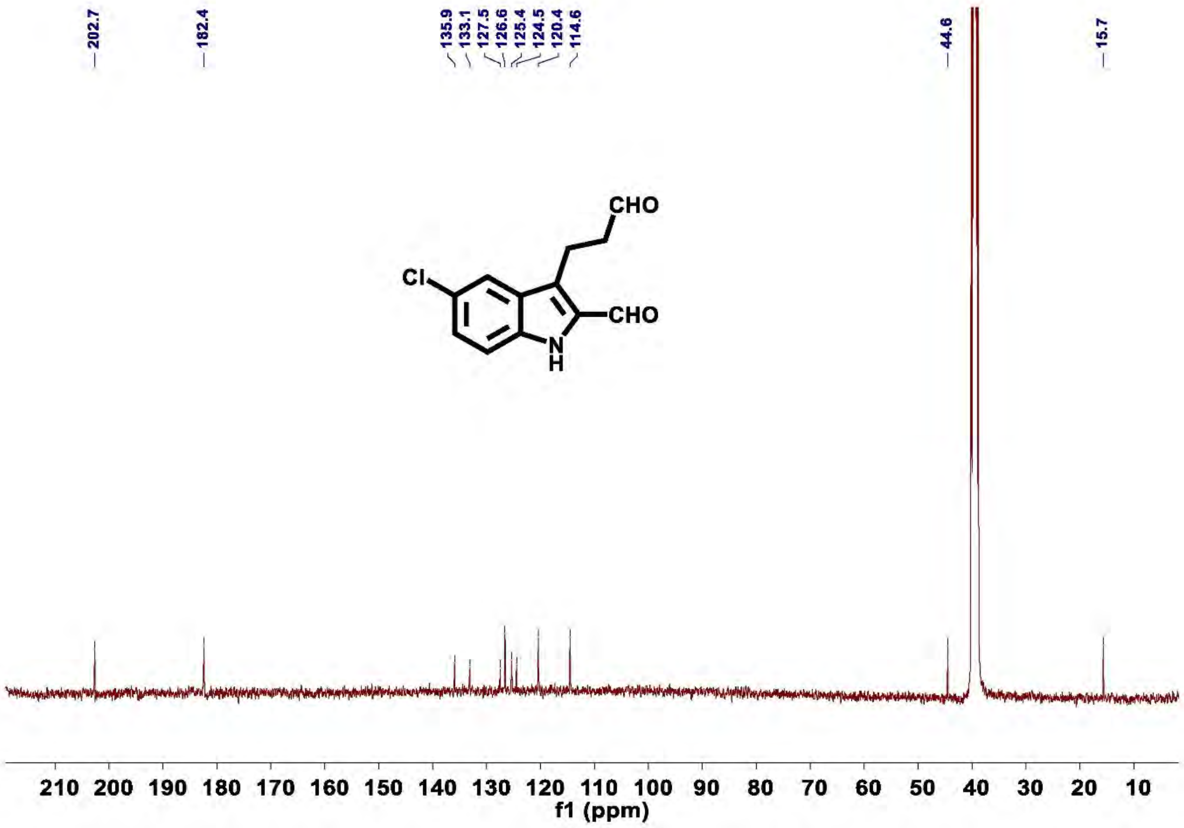

Figure S29. $^{13}C$ NMR of compound 5a

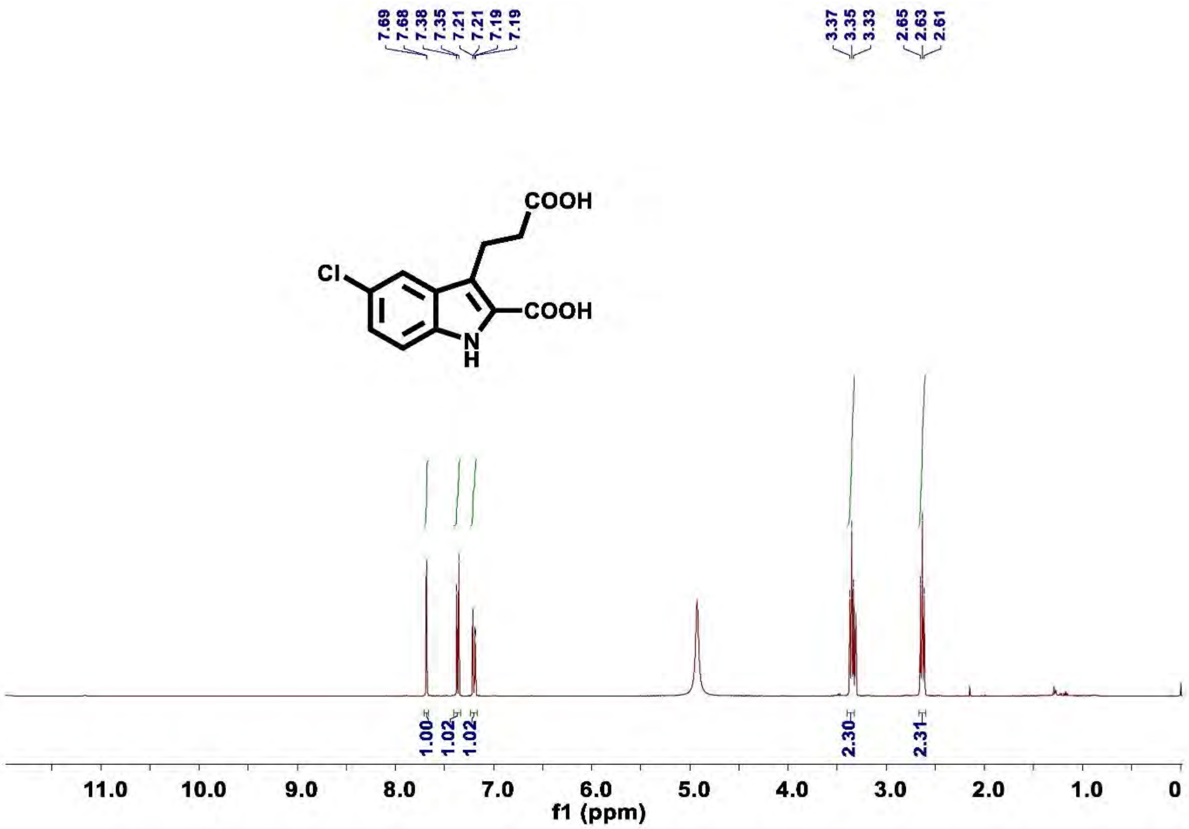

**Figure S30. $^{1}$H NMR of compound 6a**

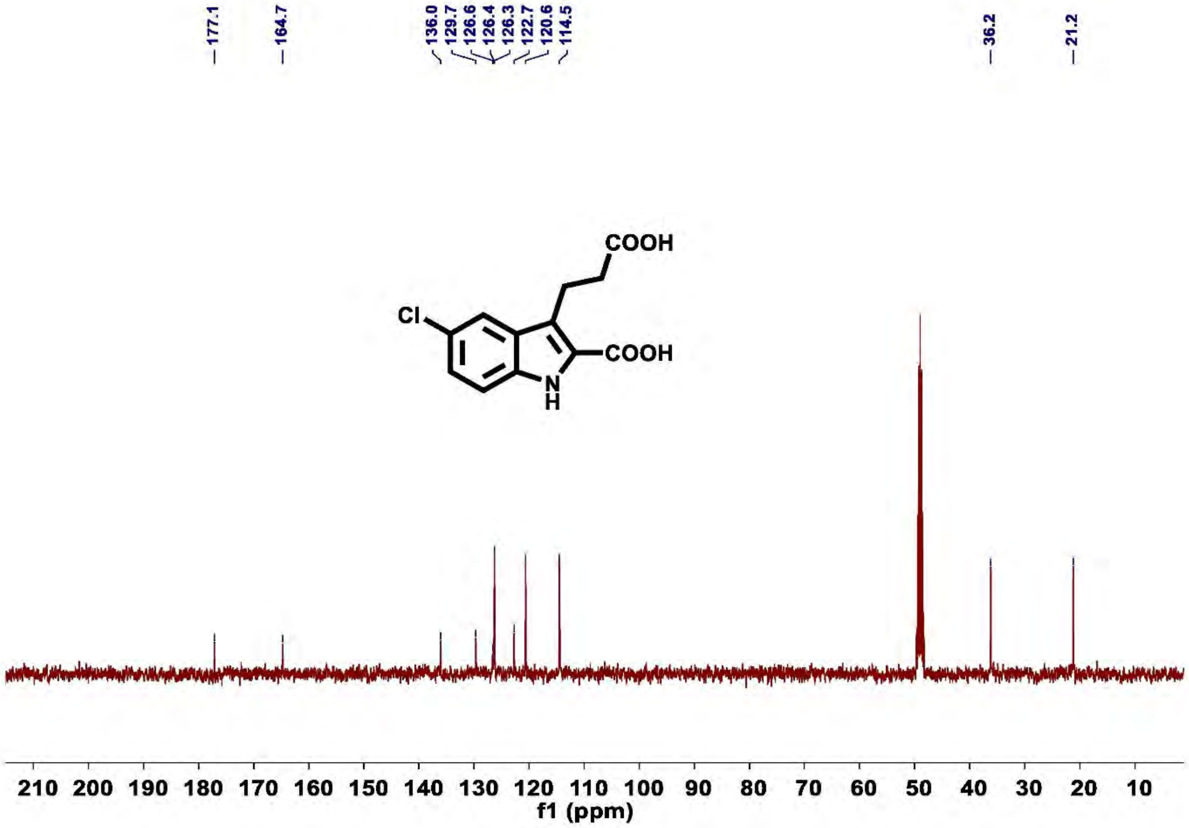

**Figure S31. $^{13}$C NMR of compound 6a**

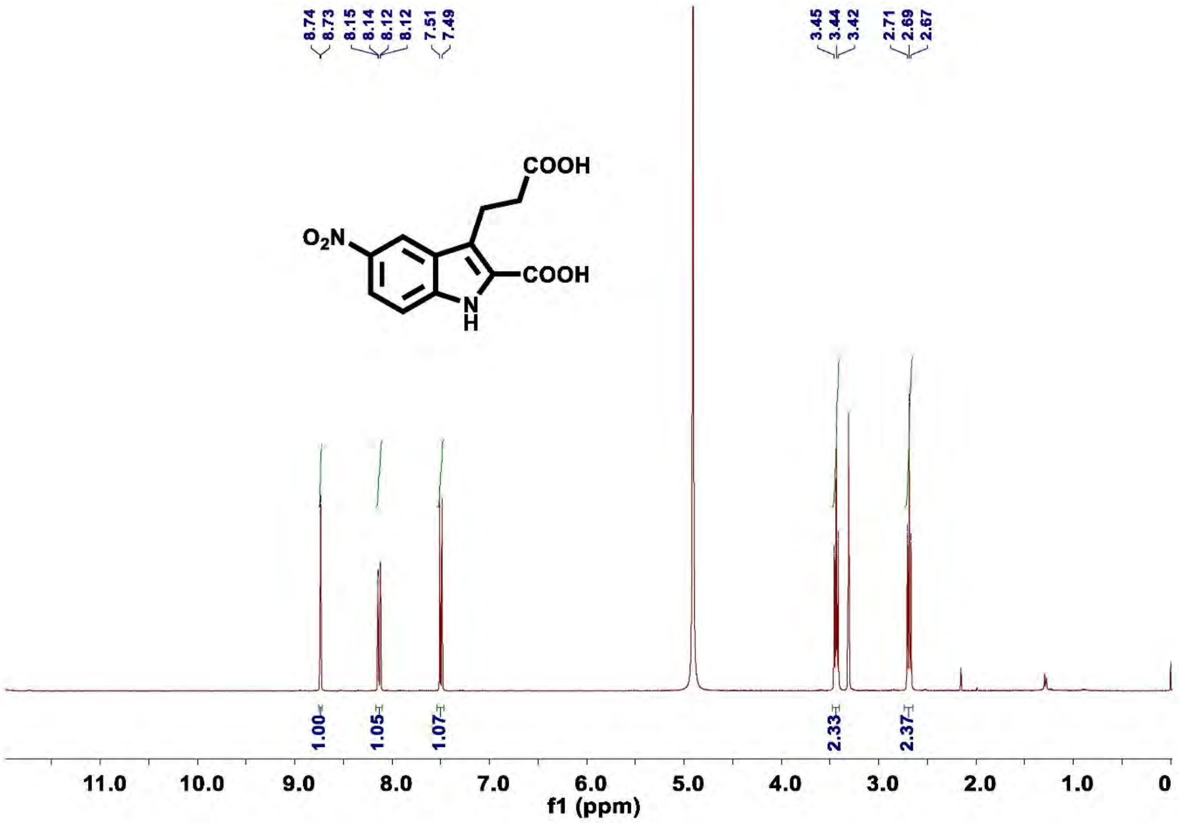

**Figure S32. $^{1}$H NMR of compound 6b**

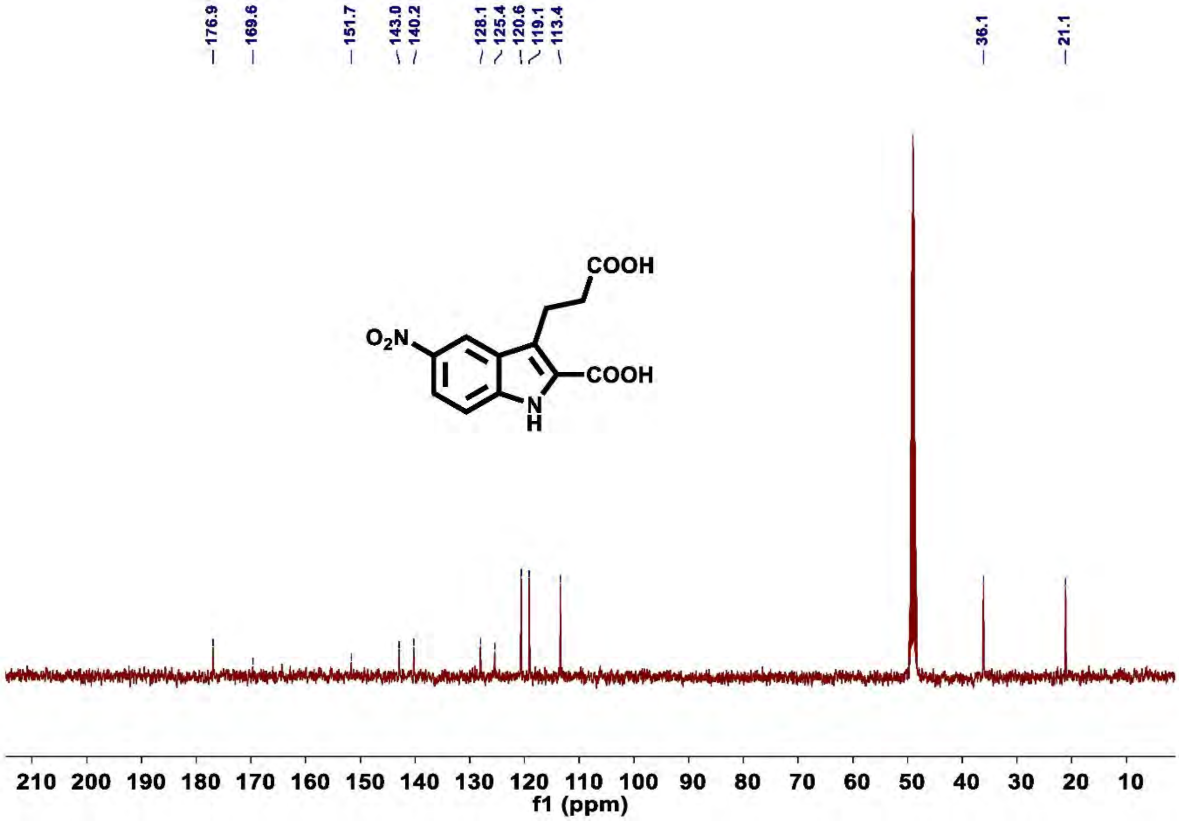

**Figure S33. $^{13}$C NMR of compound 6b**

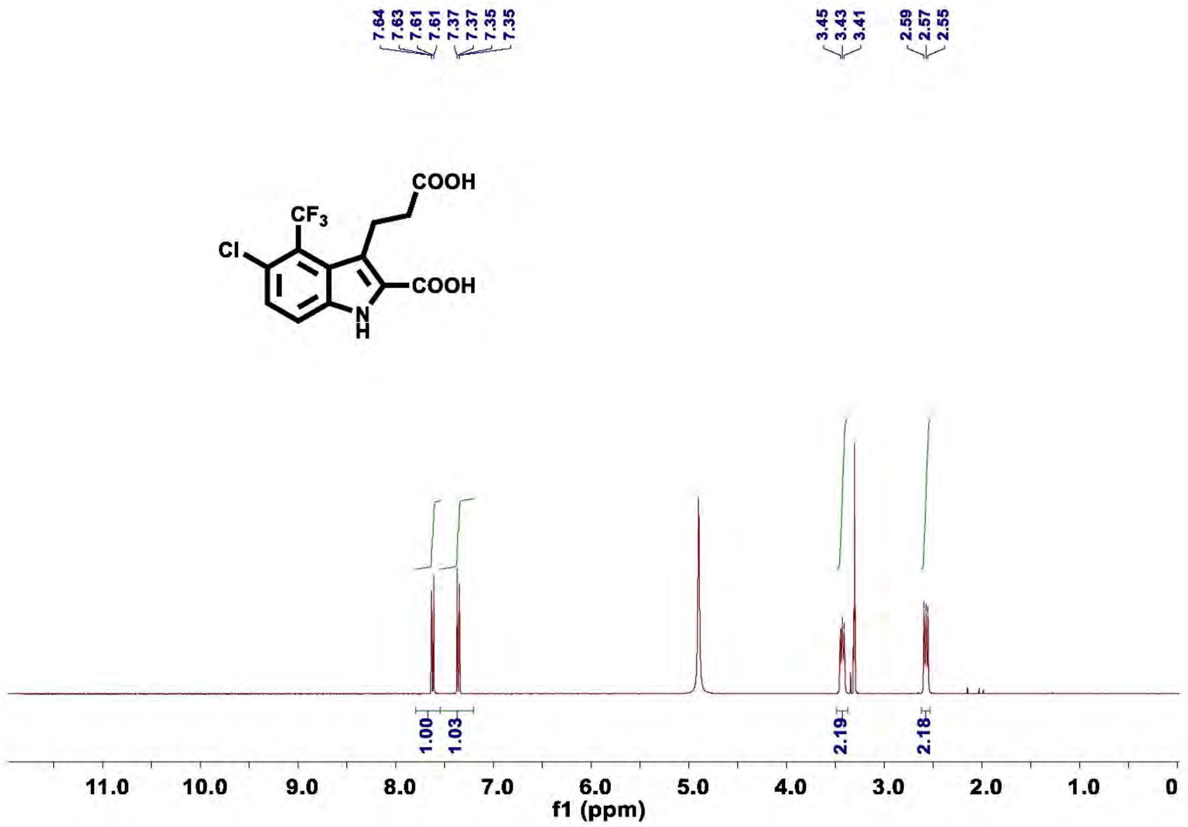

**Figure S34.** [1]**H NMR of compound 6c**

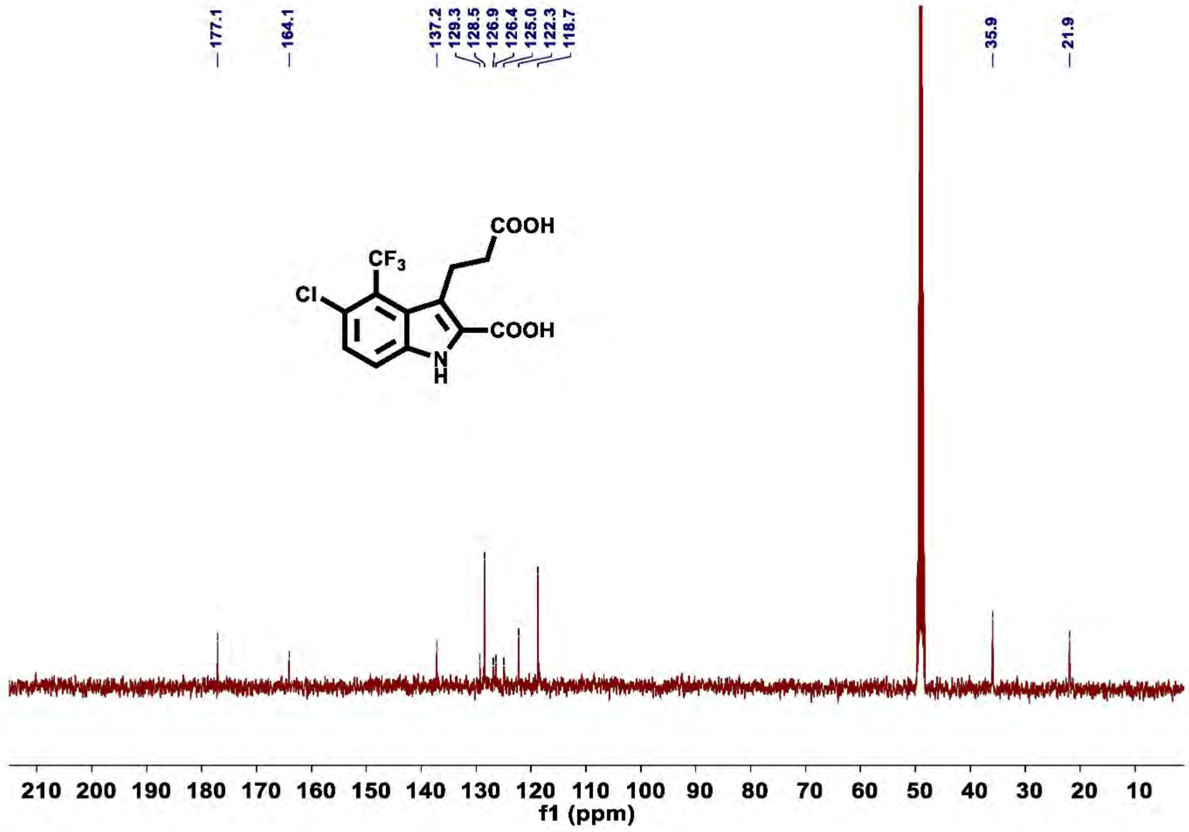

**Figure S35.** [13]**C NMR of compound 6c**

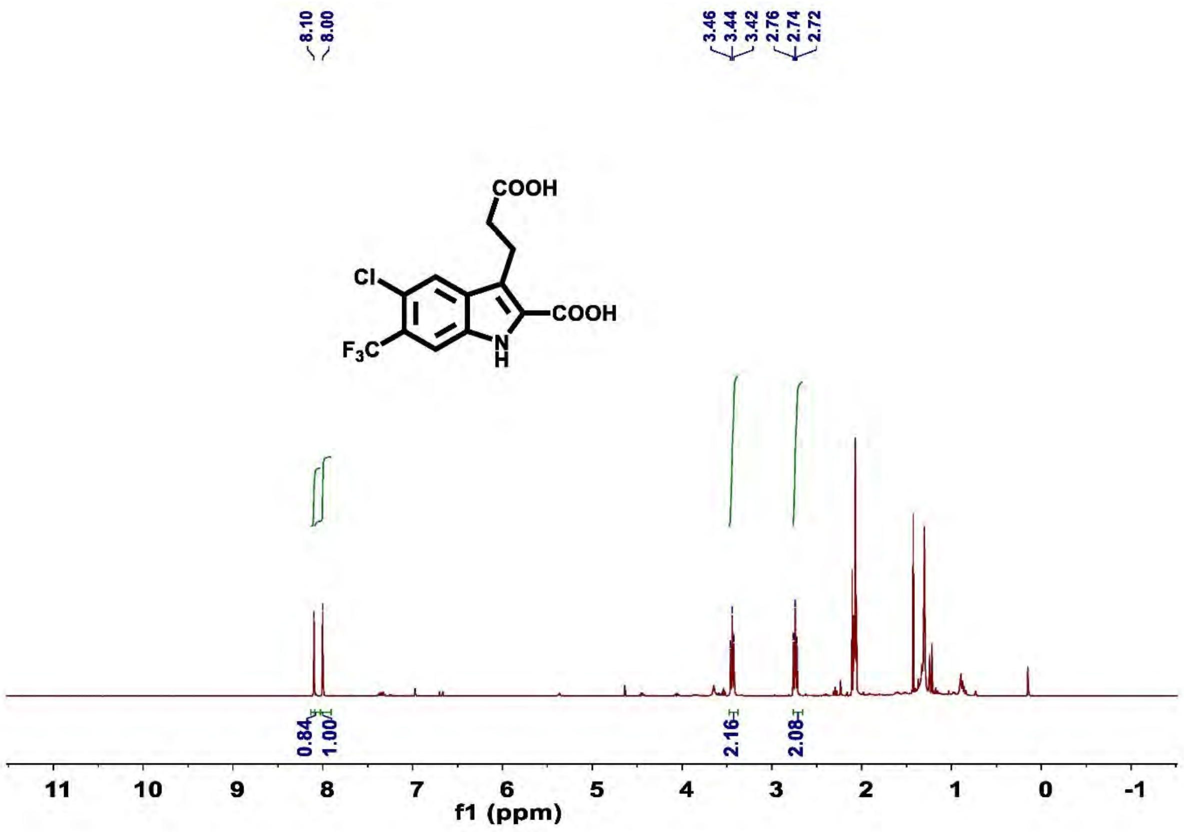

**Figure S36.** $^1$**H NMR of compound 6c'**

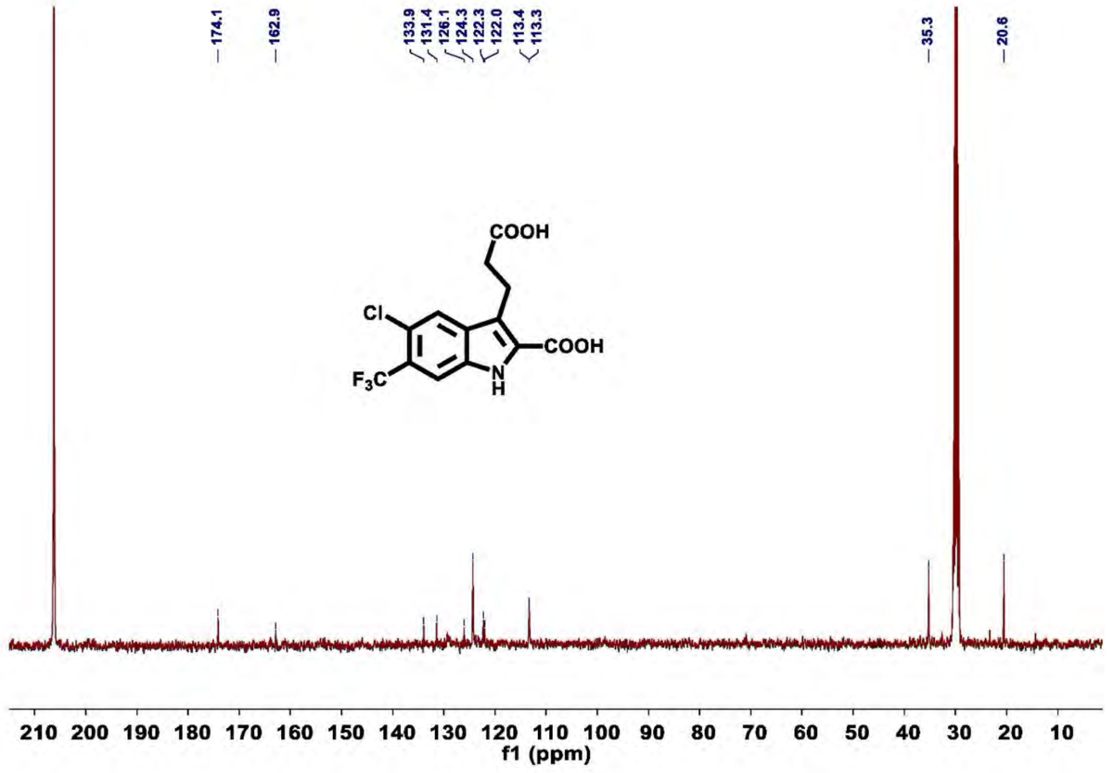

**Figure S37.** $^{13}$**C NMR of compound 6c'**

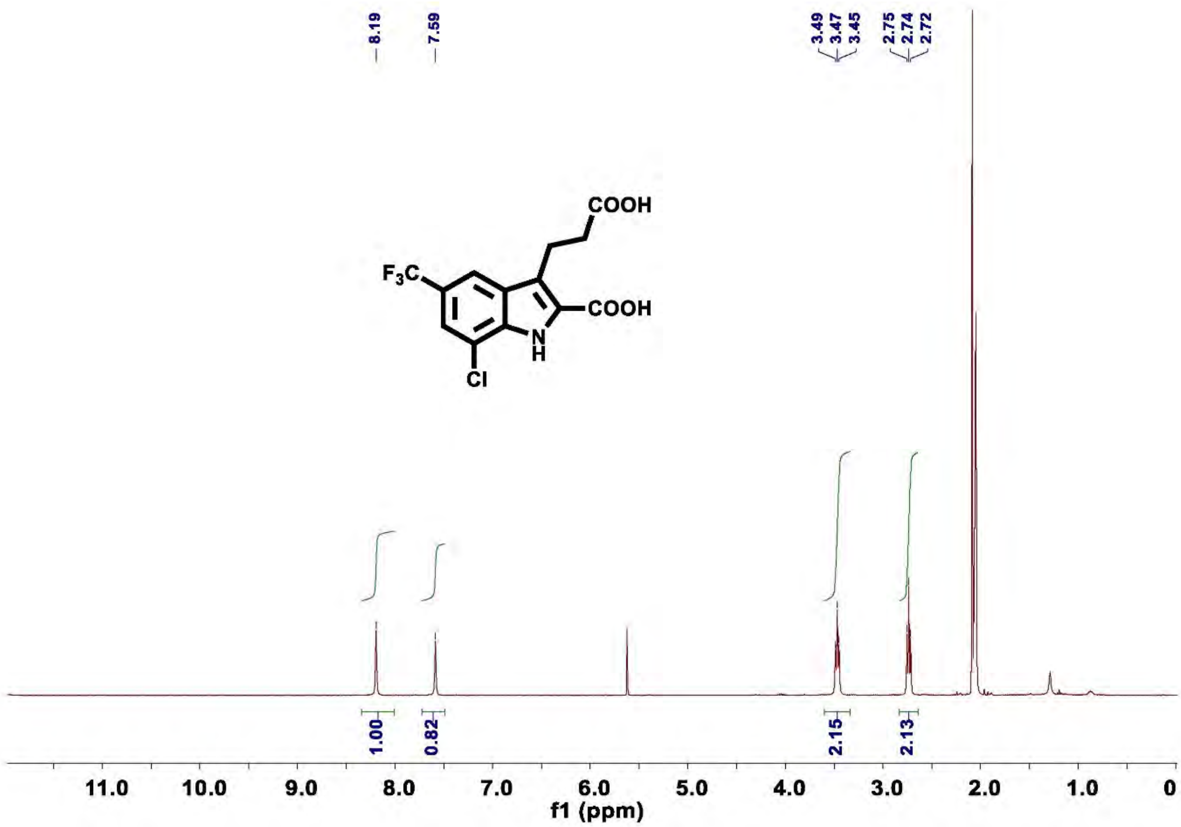

**Figure S38. $^{1}$H NMR of compound 6d**

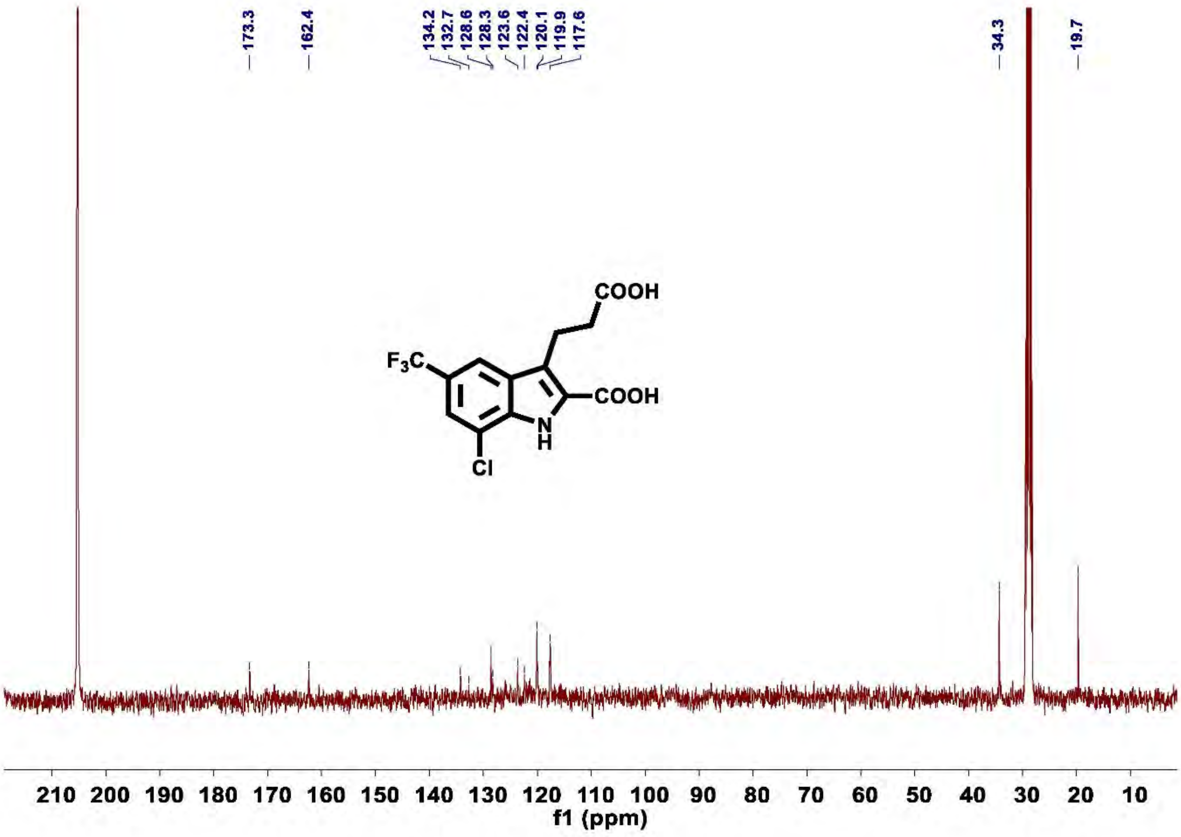

**Figure S39. $^{13}$C NMR of compound 6d**

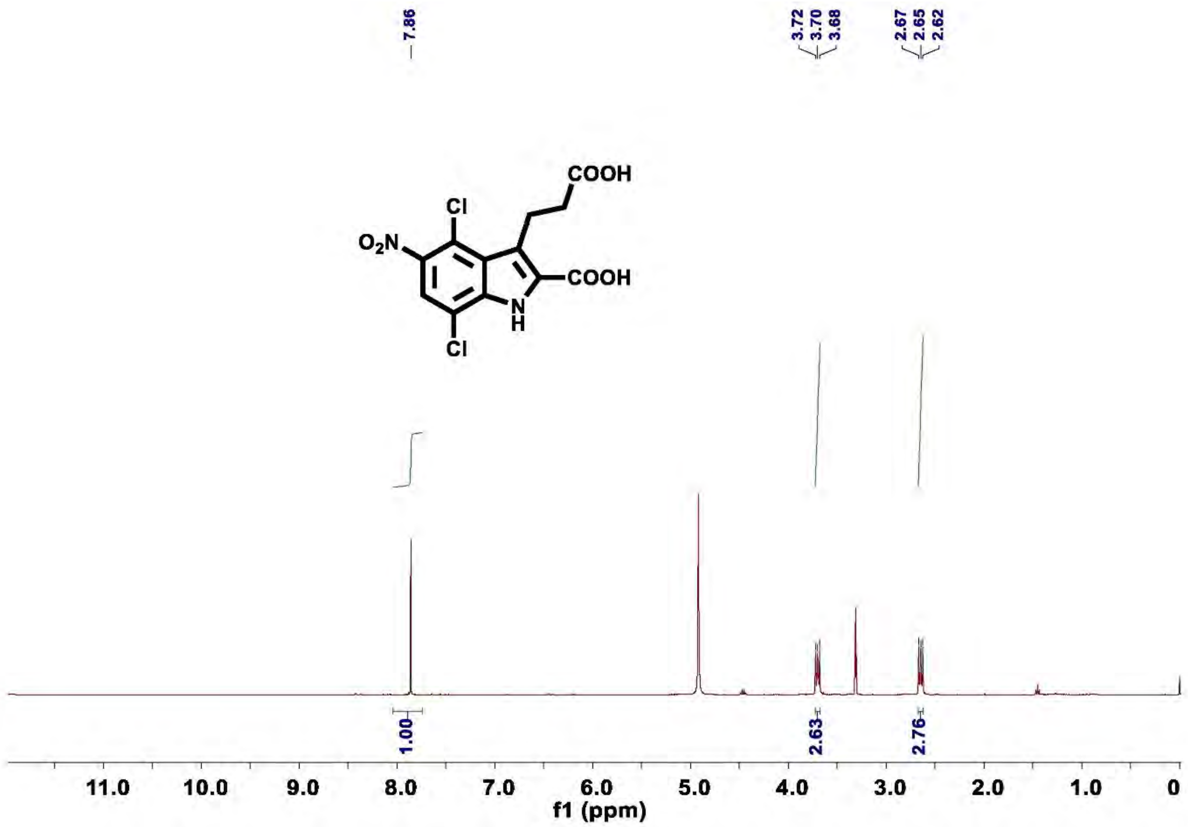

Figure 40. $^{1}$H NMR of compound 6e

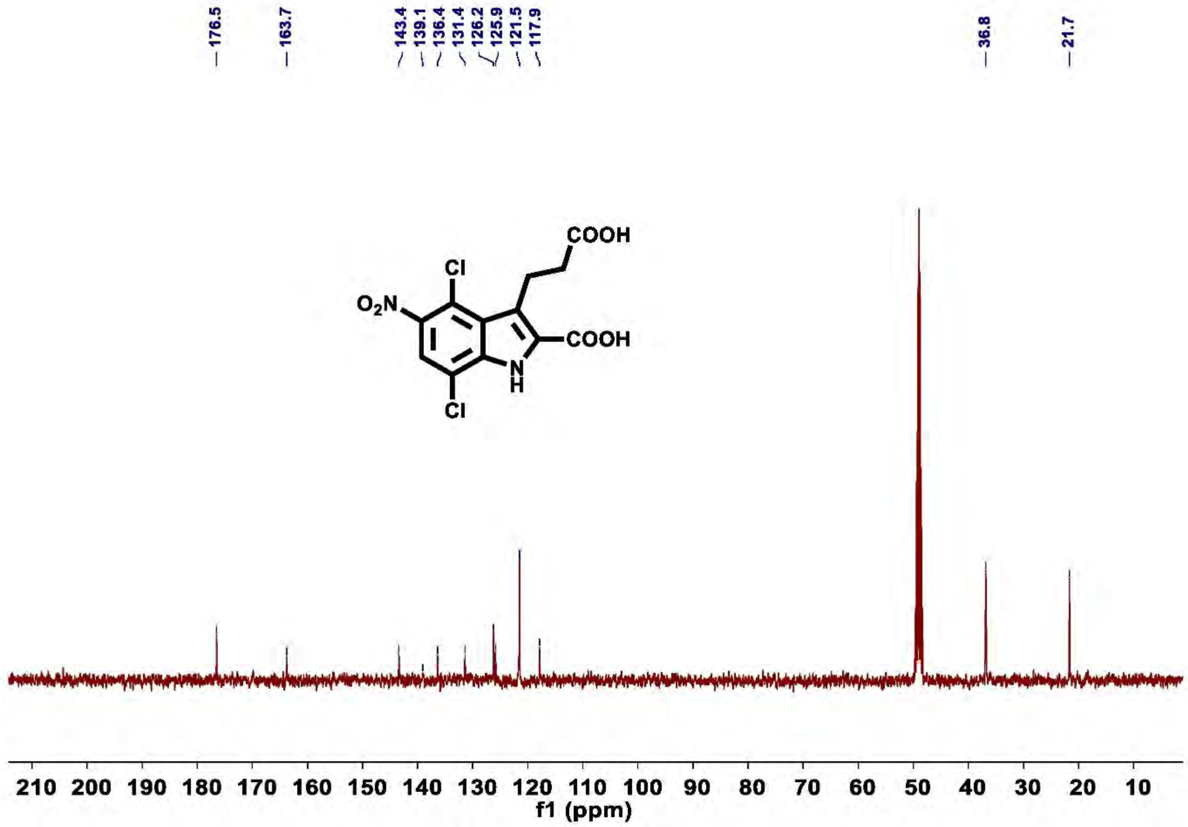

Figure S41. $^{13}$C NMR of compound 6e

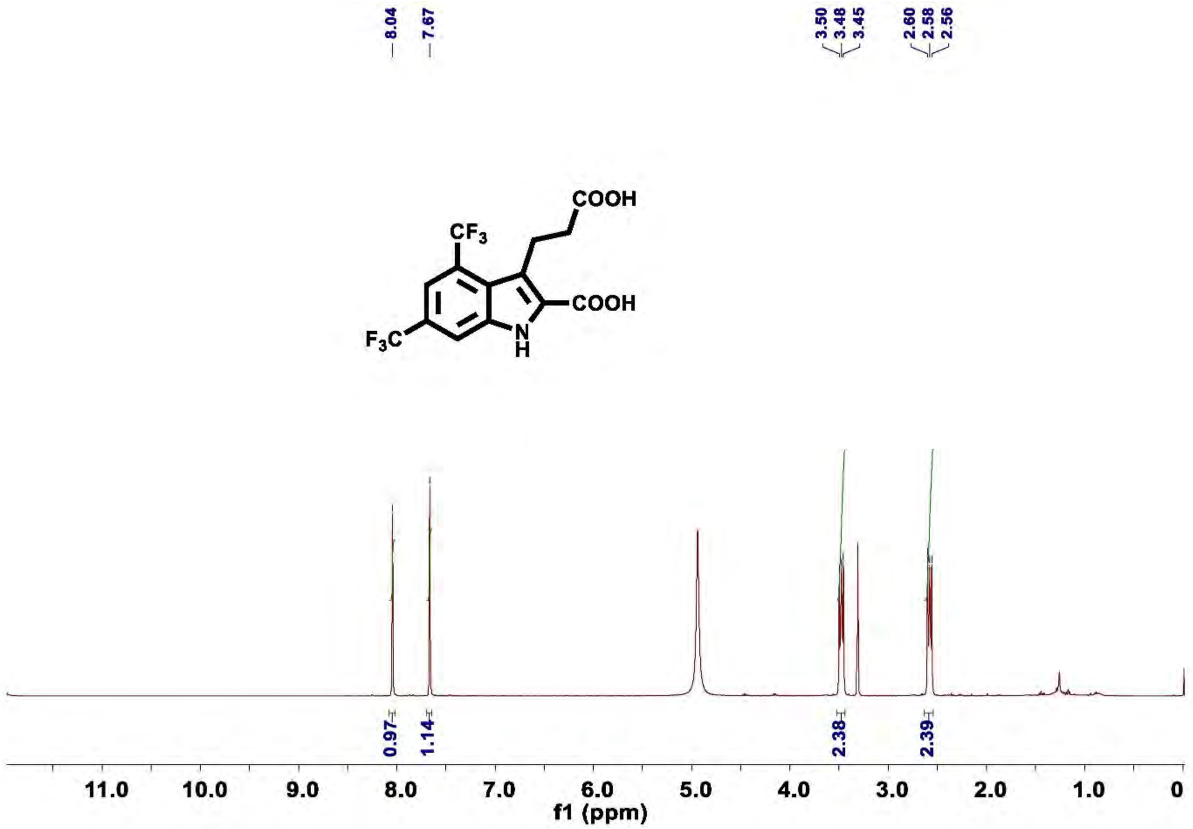

**Figure S42.** $^{1}$**H NMR of compound 6f**

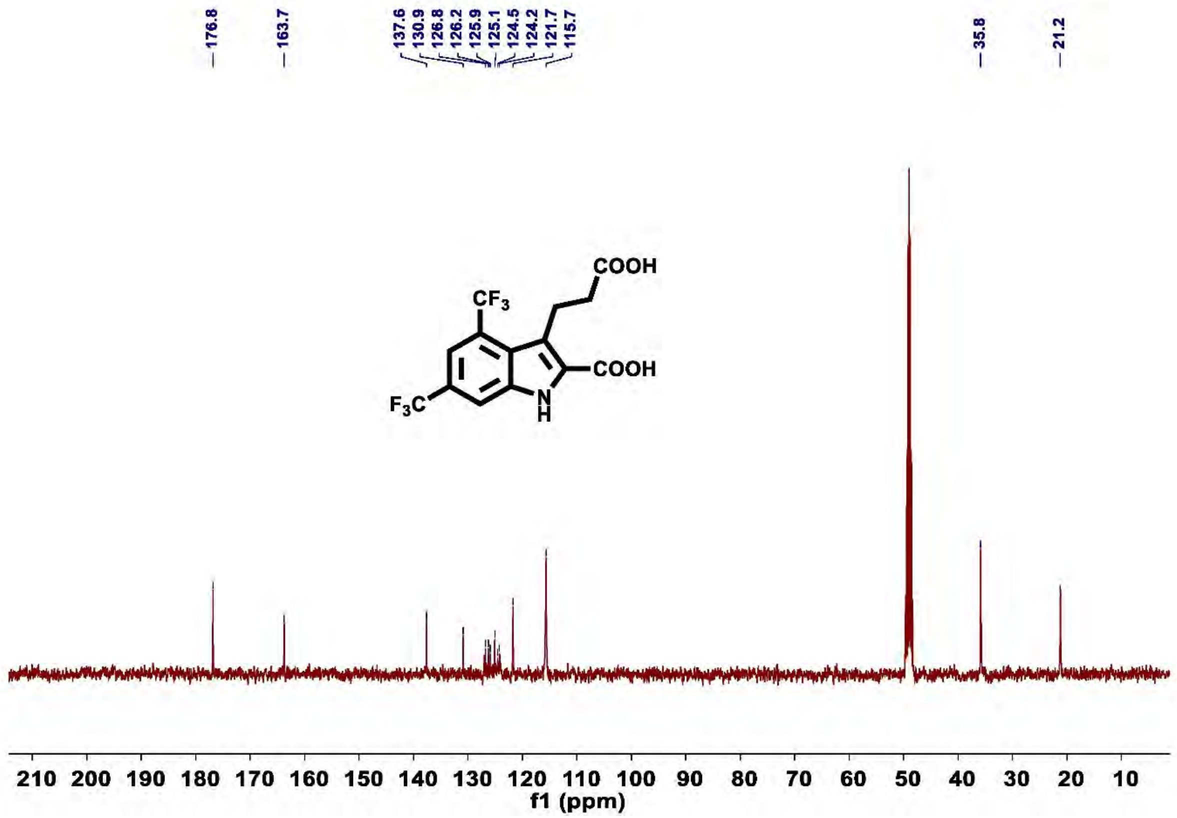

**Figure S43.** $^{13}$**C NMR of compound 6f**

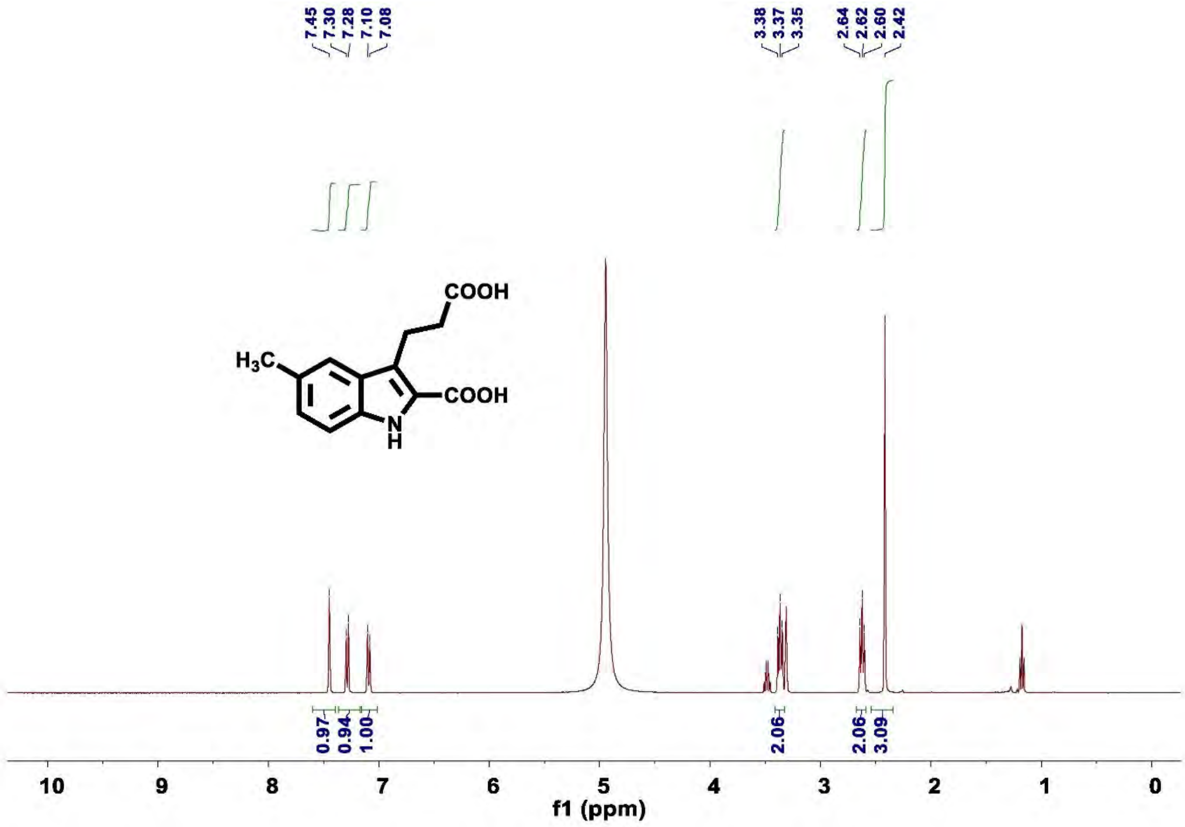

Figure S44. [1]H NMR of compound 6g

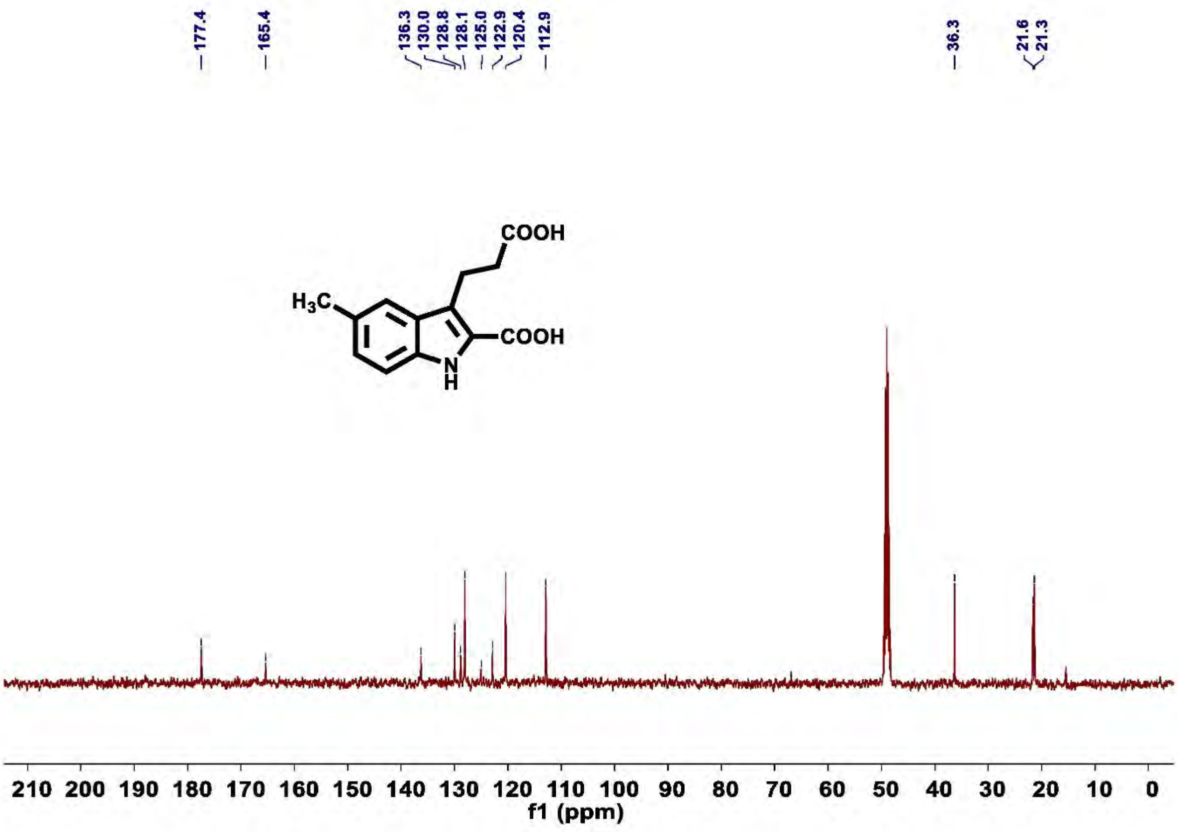

Figure S45. [13]C NMR of compound 6g

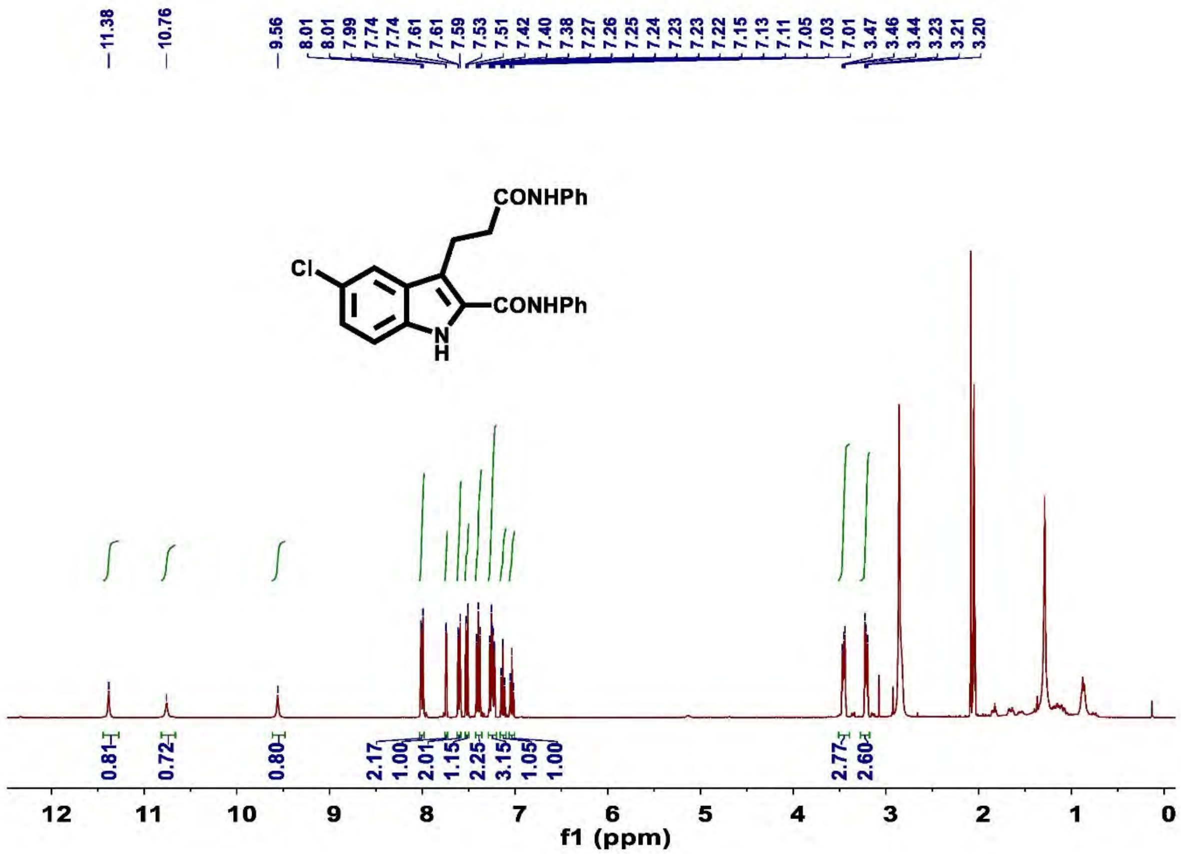

**Figure S46. $^1$H NMR of compound 7a**

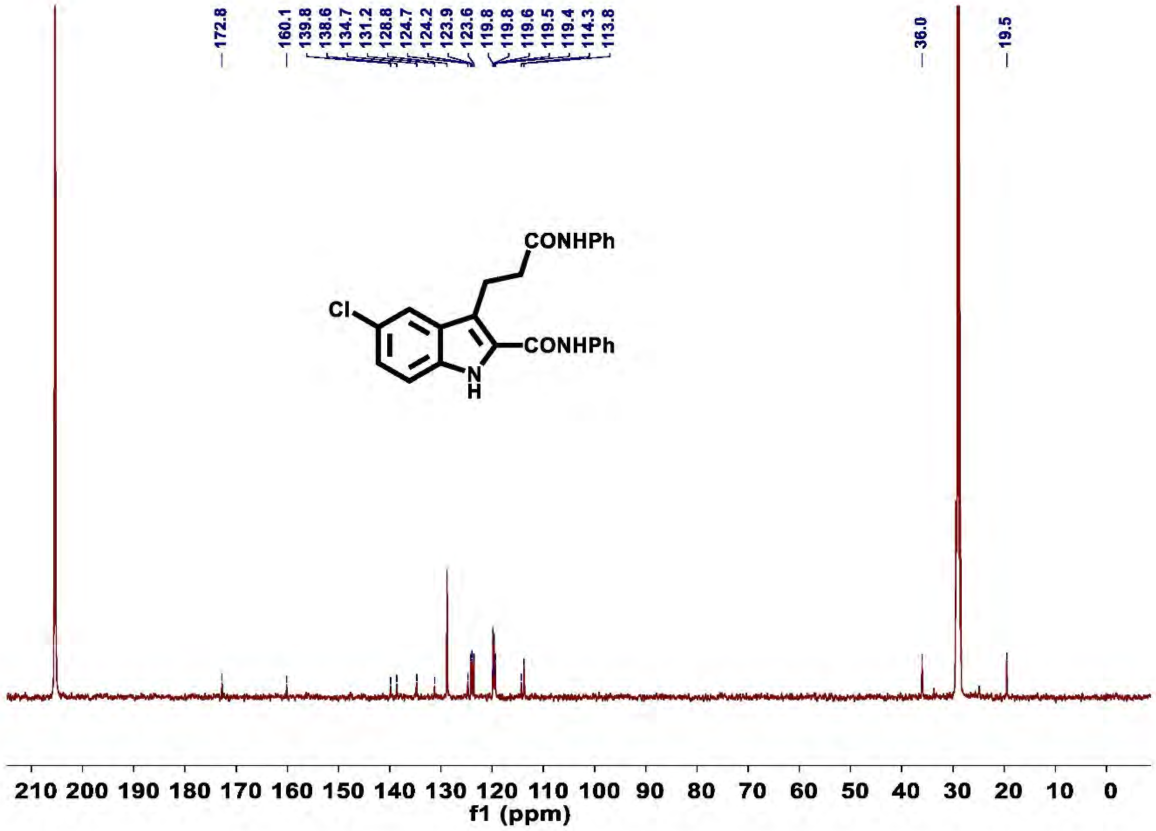

**Figure S47.** $^{13}$C NMR of compound 7a

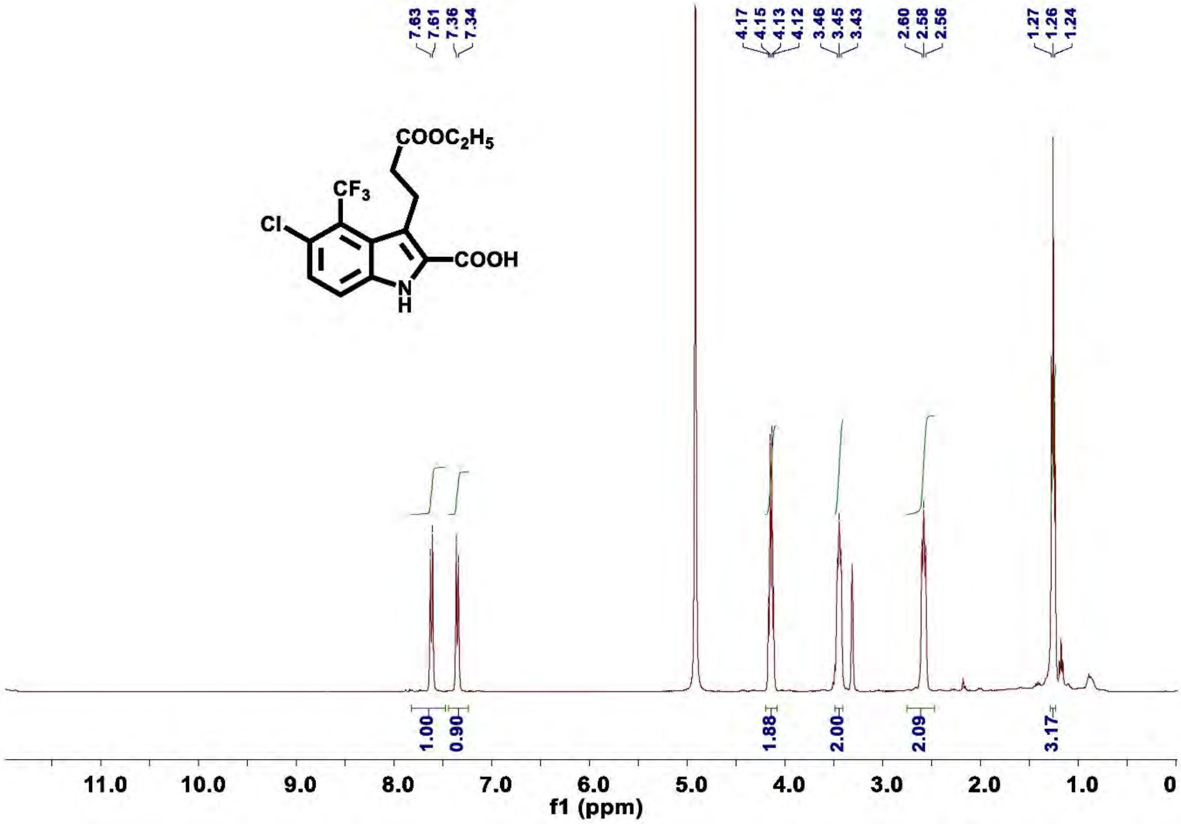

**Figure S48.** $^{1}$H NMR of compound 8c

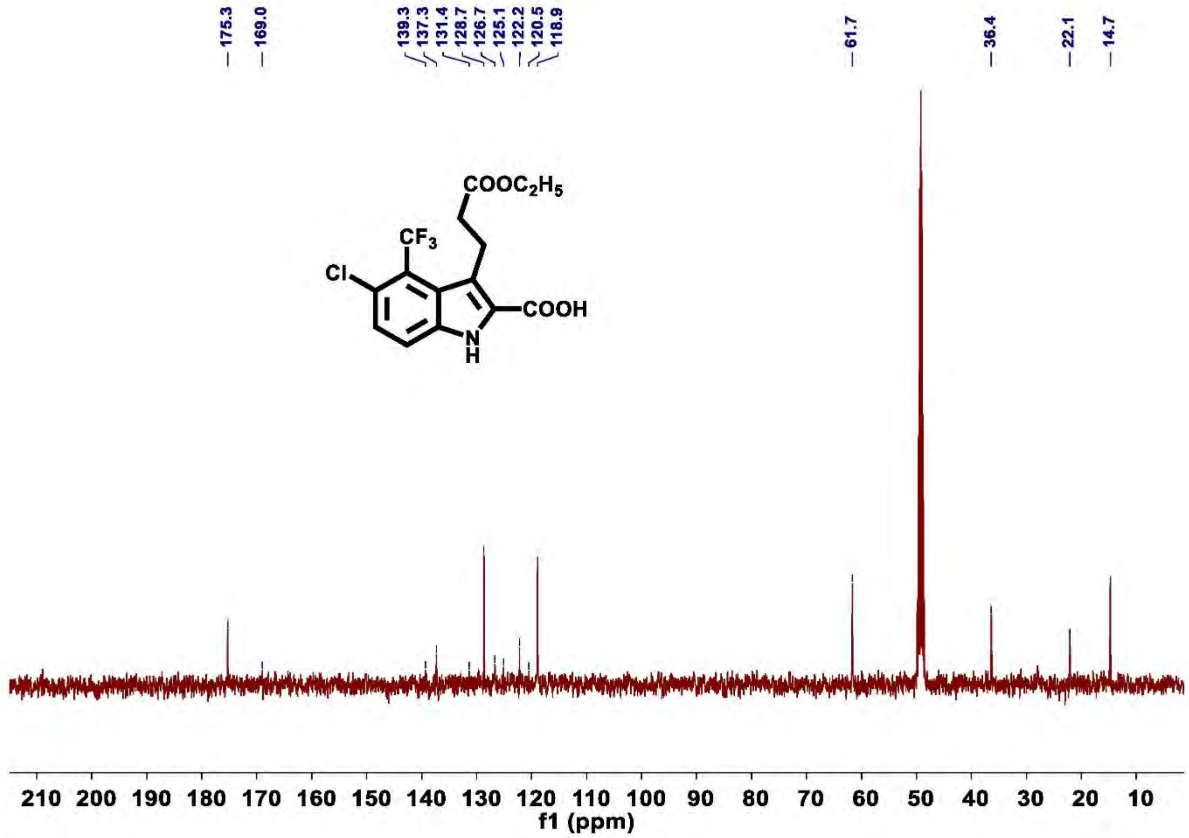

**Figure S49.** $^{13}$**C NMR of compound 8c**

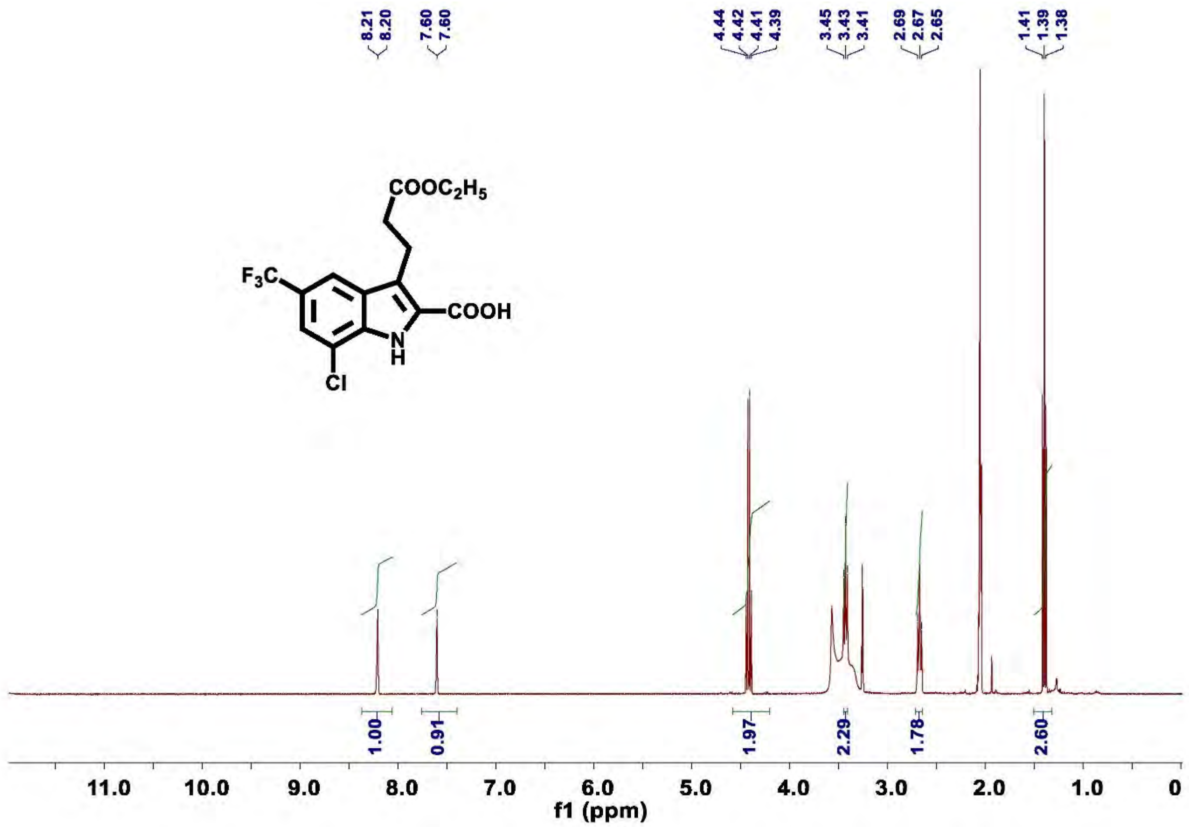

**Figure S50.** $^{1}$H **NMR of compound 8d**

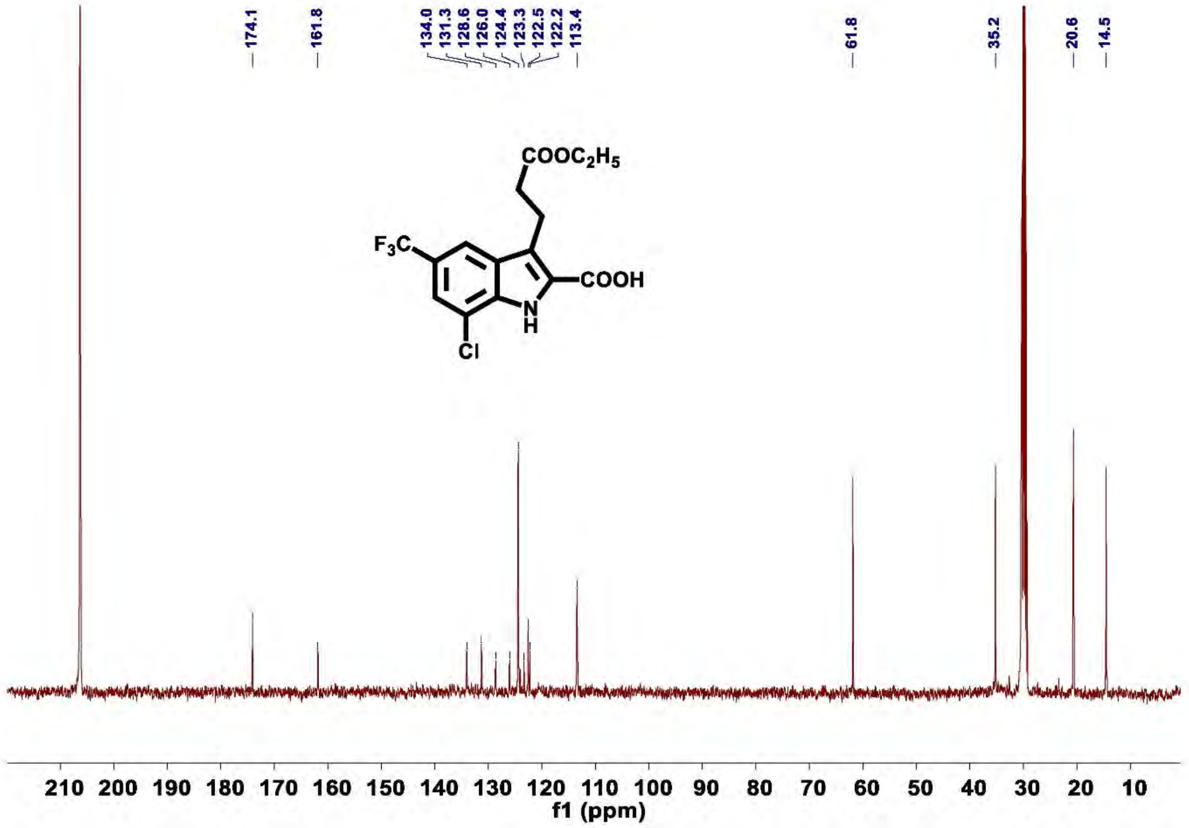

**Figure S51.** $^{13}$C **NMR of compound 8d**

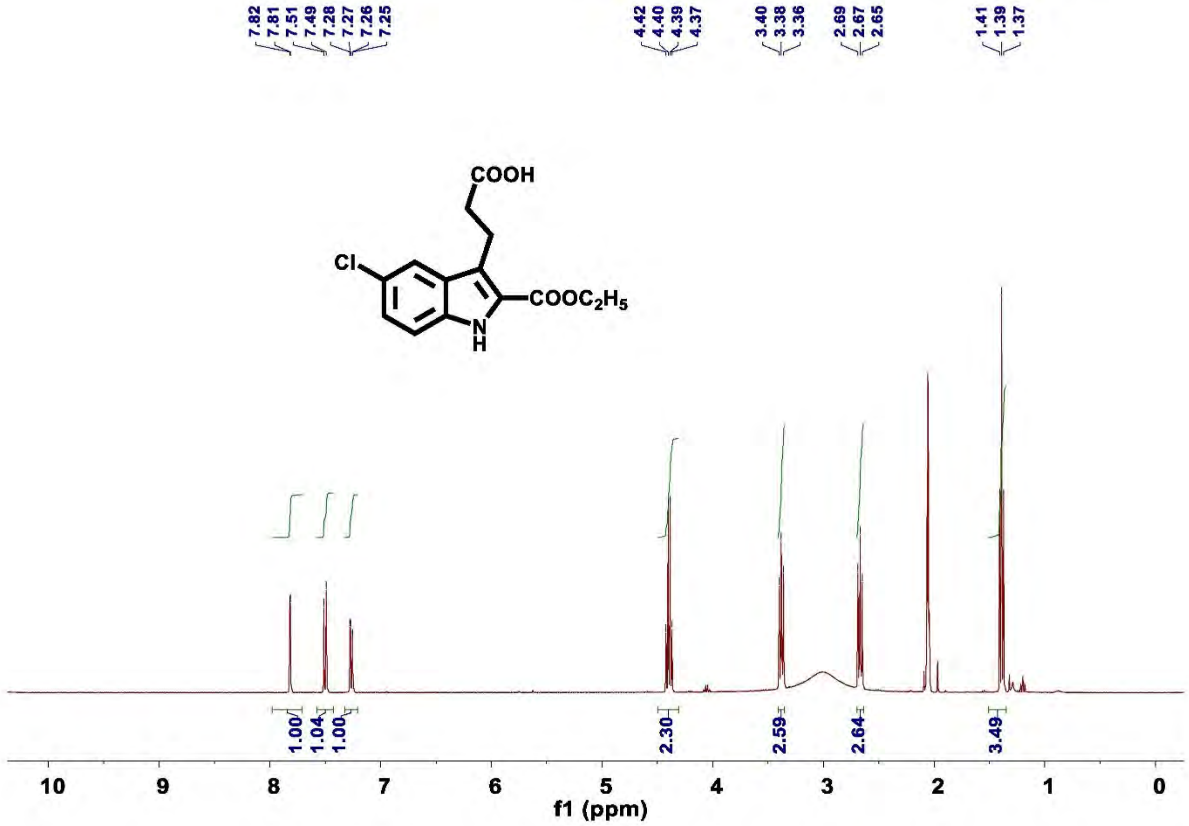

**Figure S52. [1]H NMR of compound 9a**

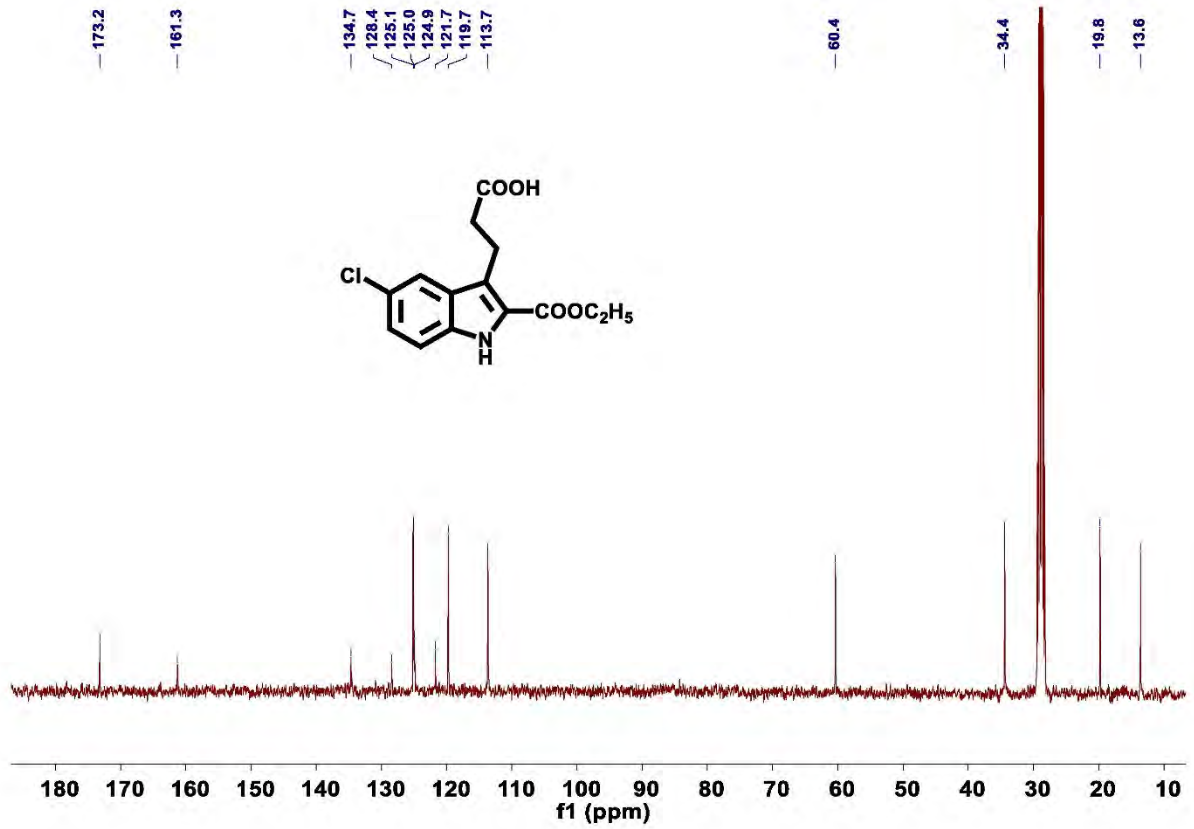

**Figure S53.** $^{13}$**C NMR of compound 9a**

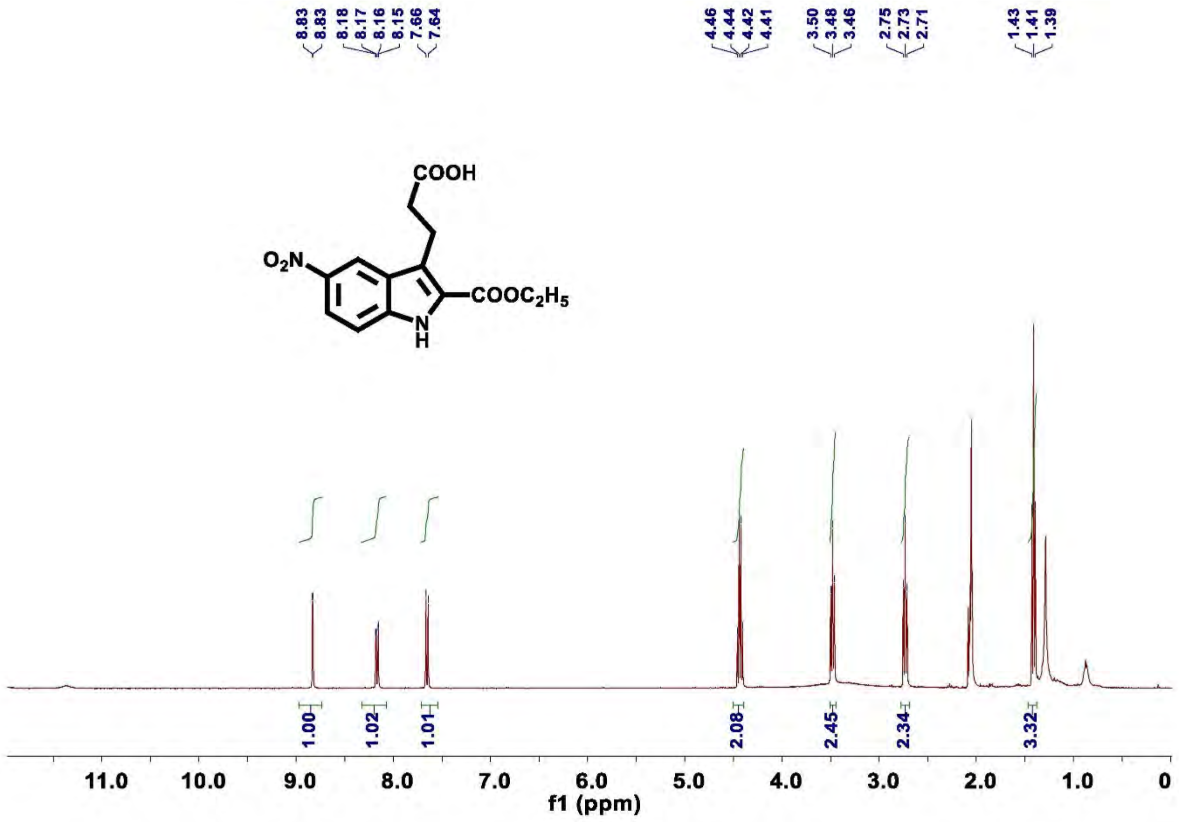

**Figure S54.** $^{1}$**H NMR of compound 9b**

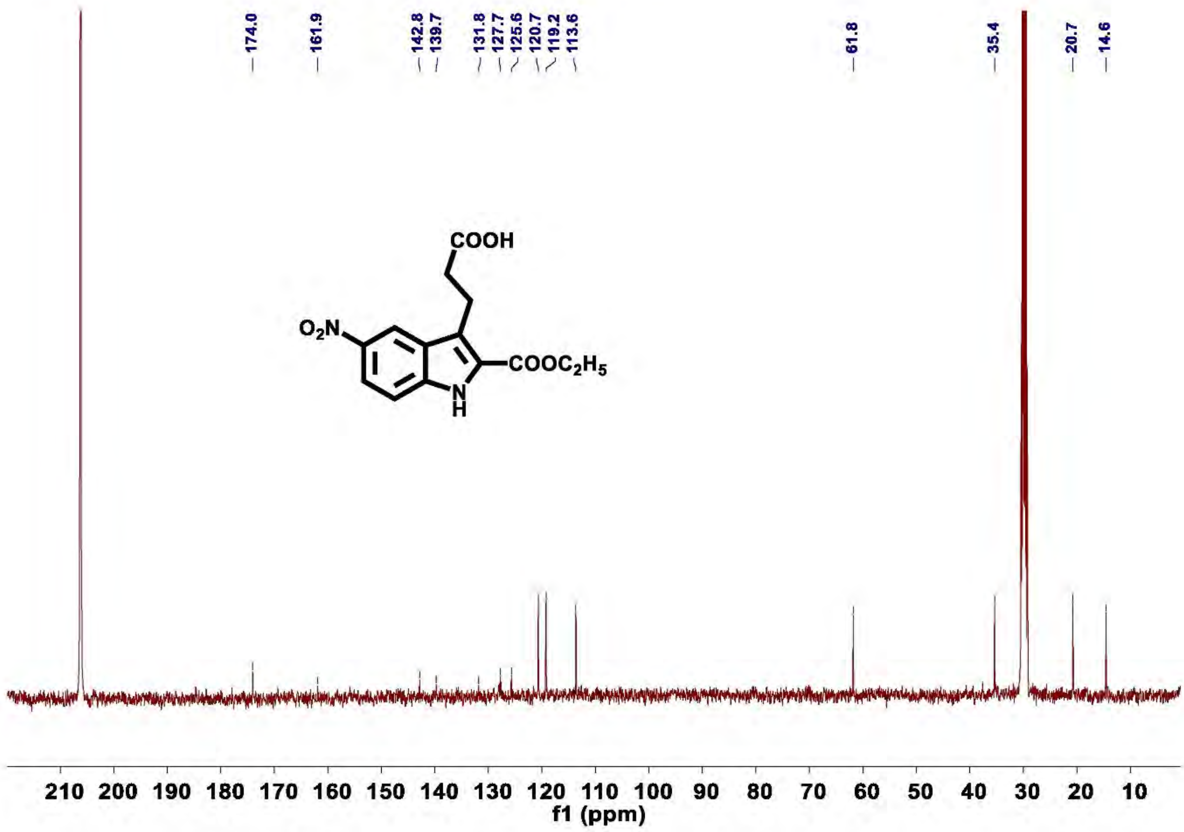

Figure S55. $^{13}C$ NMR of compound 9b

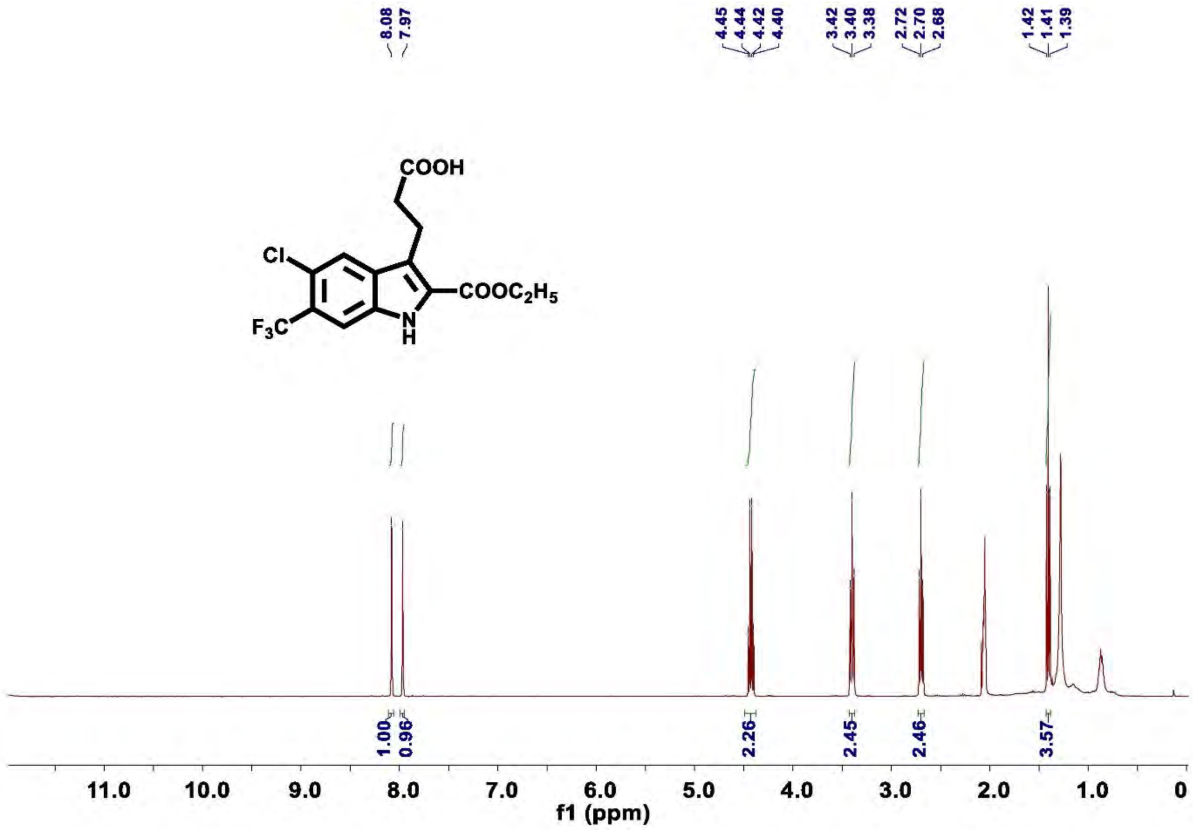

**Figure S56.** $^1$**H NMR of compound 9c'**

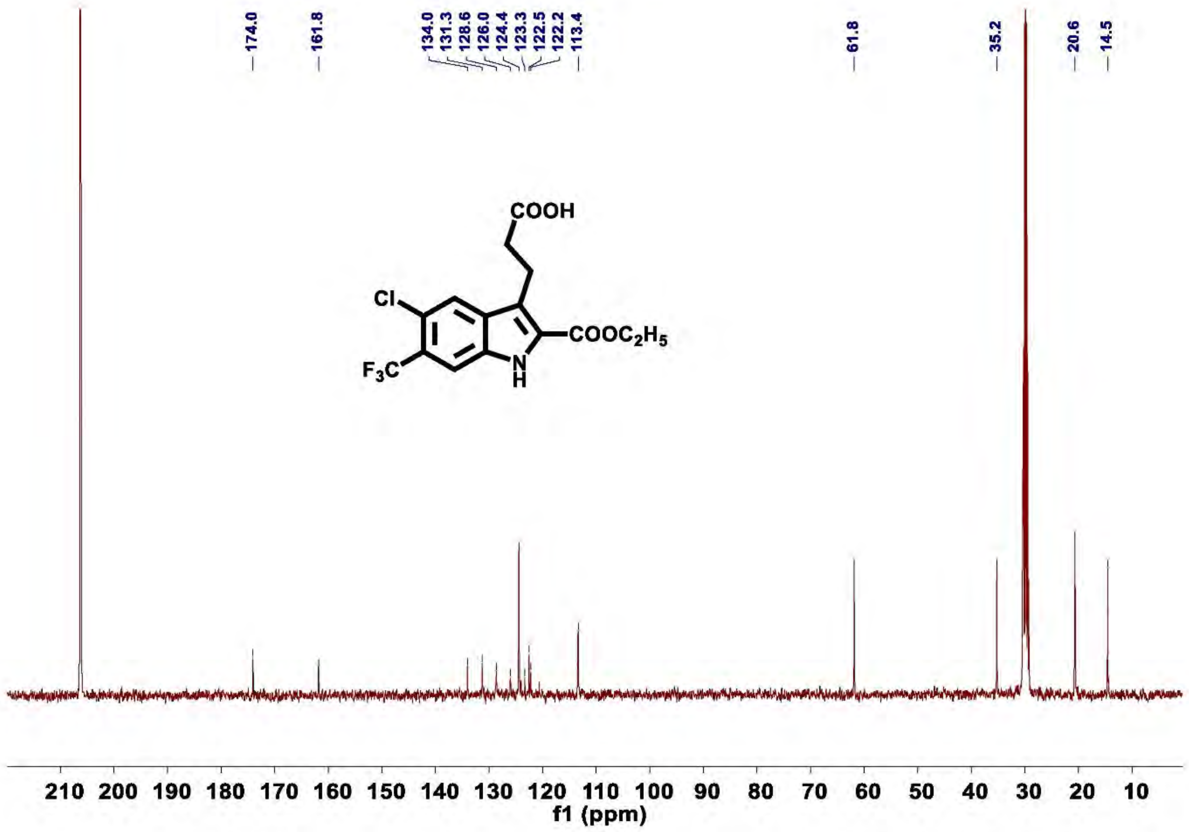

**Figure S57.** $^{13}$**C NMR of compound 9c'**

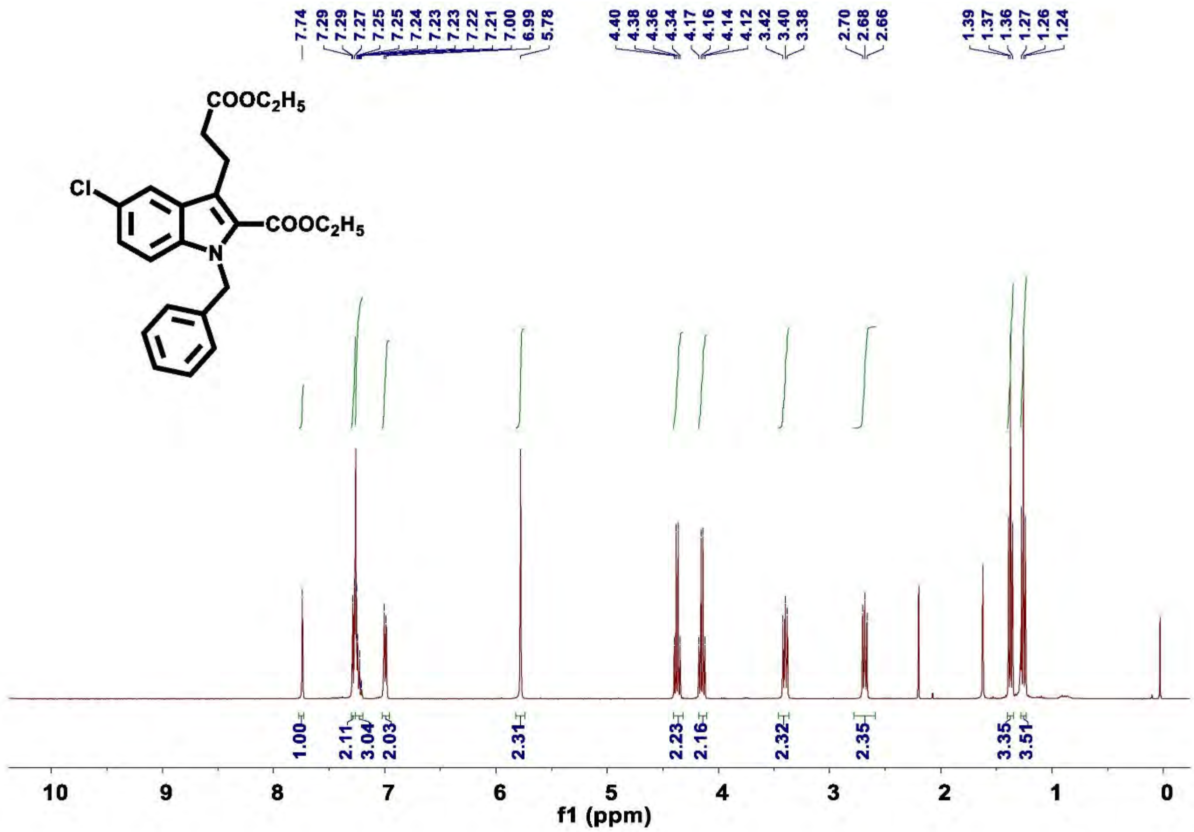

Figure S58. $^{1}$H NMR of compound 10a

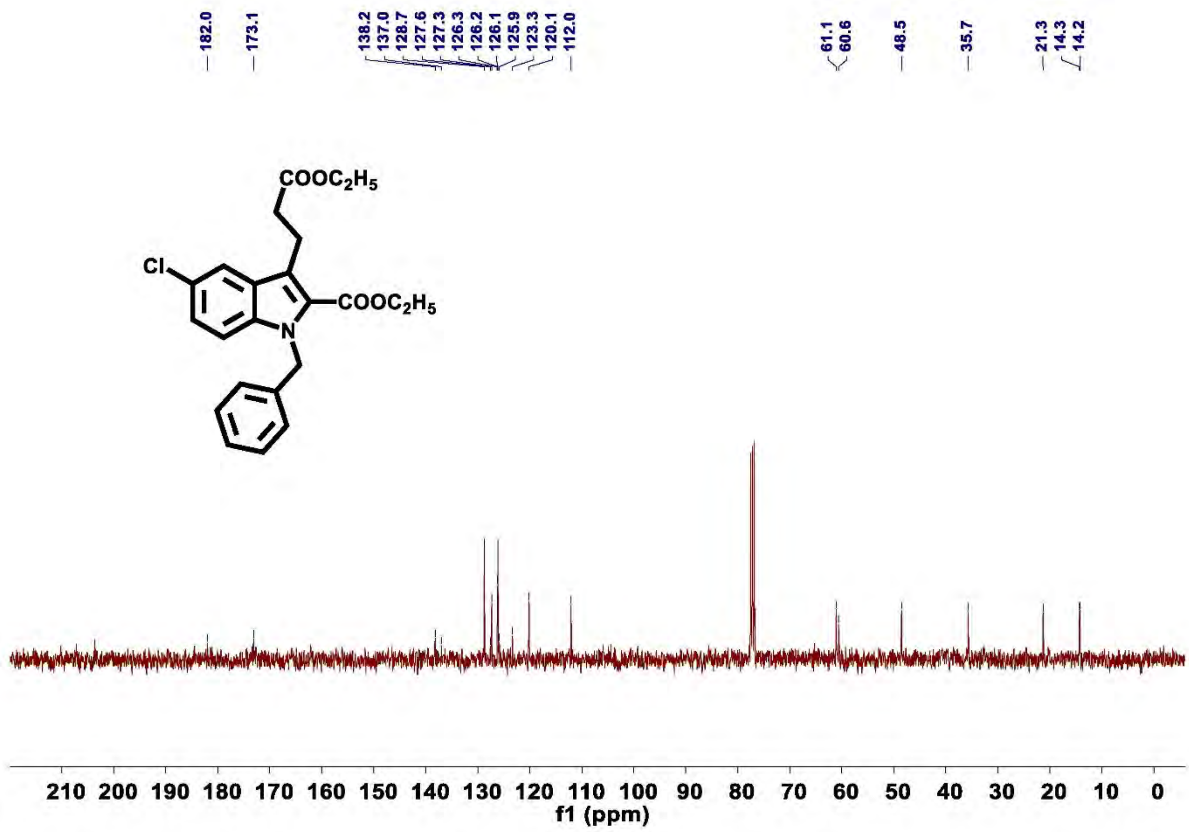

Figure S59. $^{13}$C NMR of compound 10a

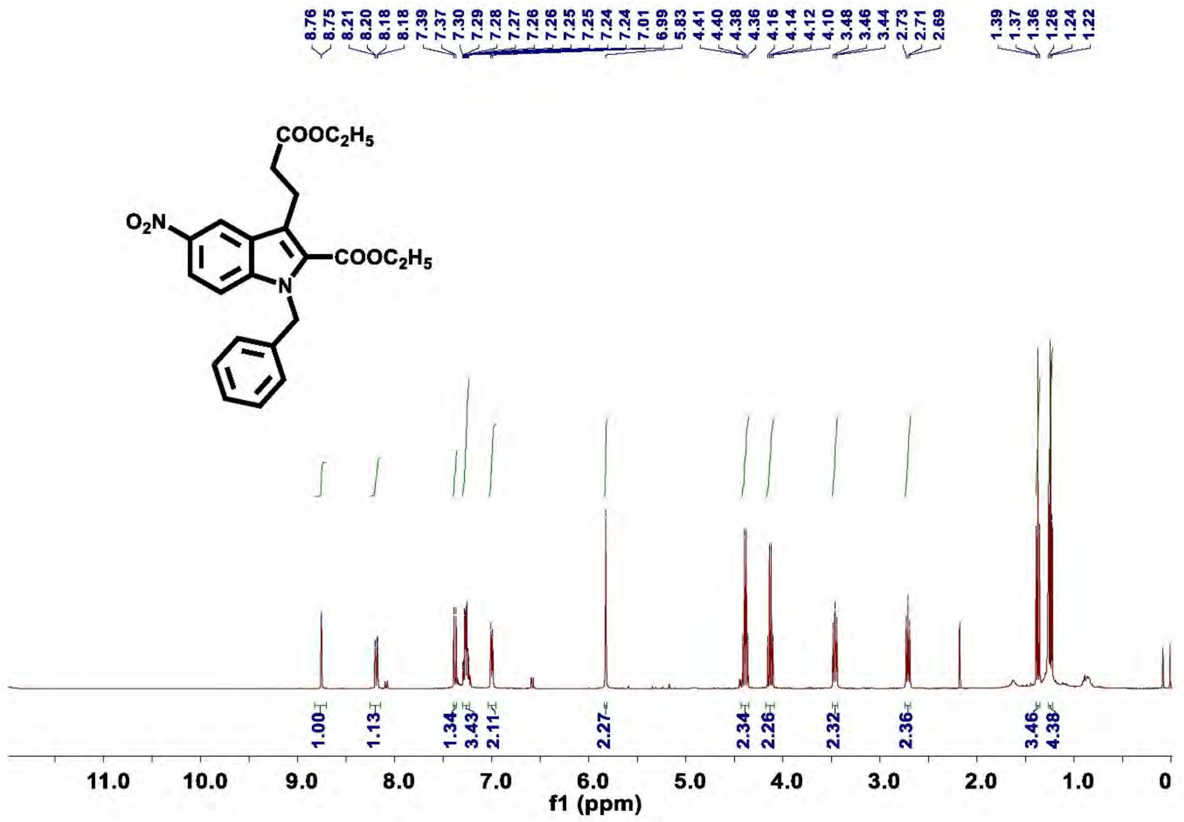

Figure S60. $^{1}$H NMR of compound 10b

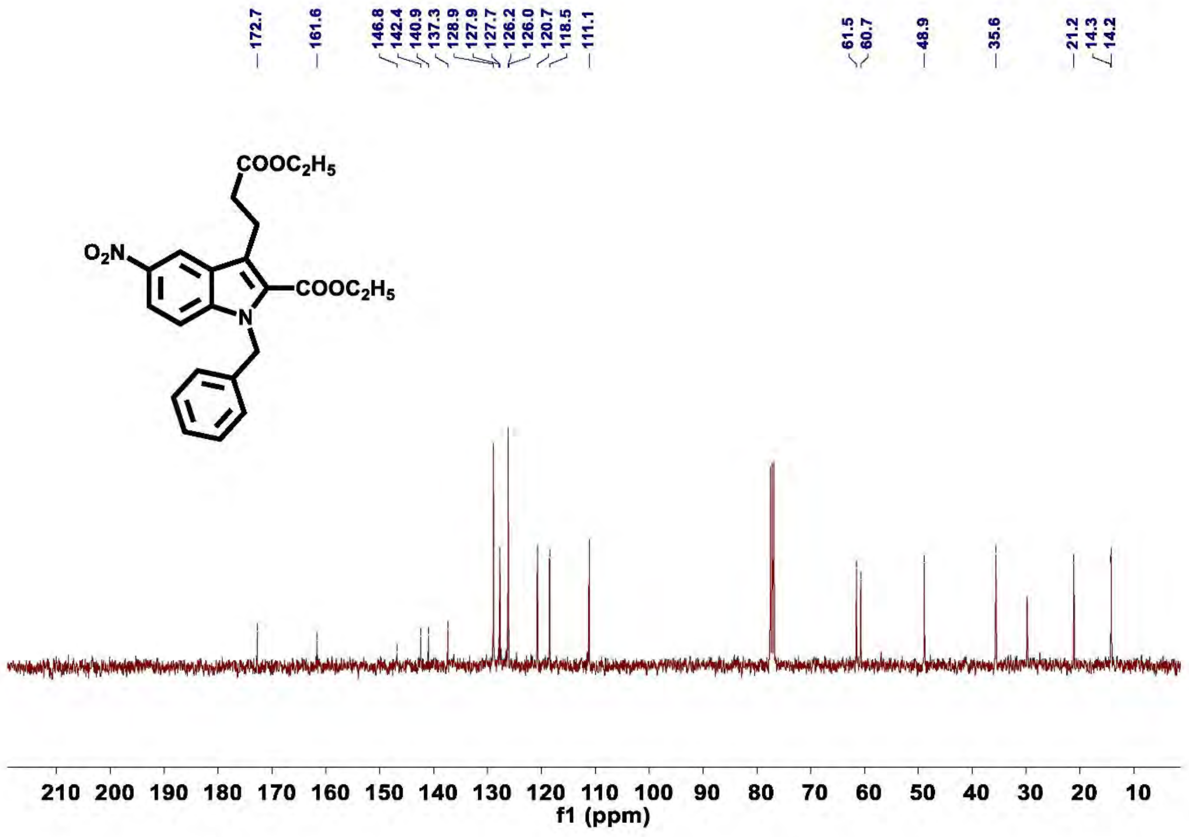

Figure S61. $^{13}$C NMR of compound 10b

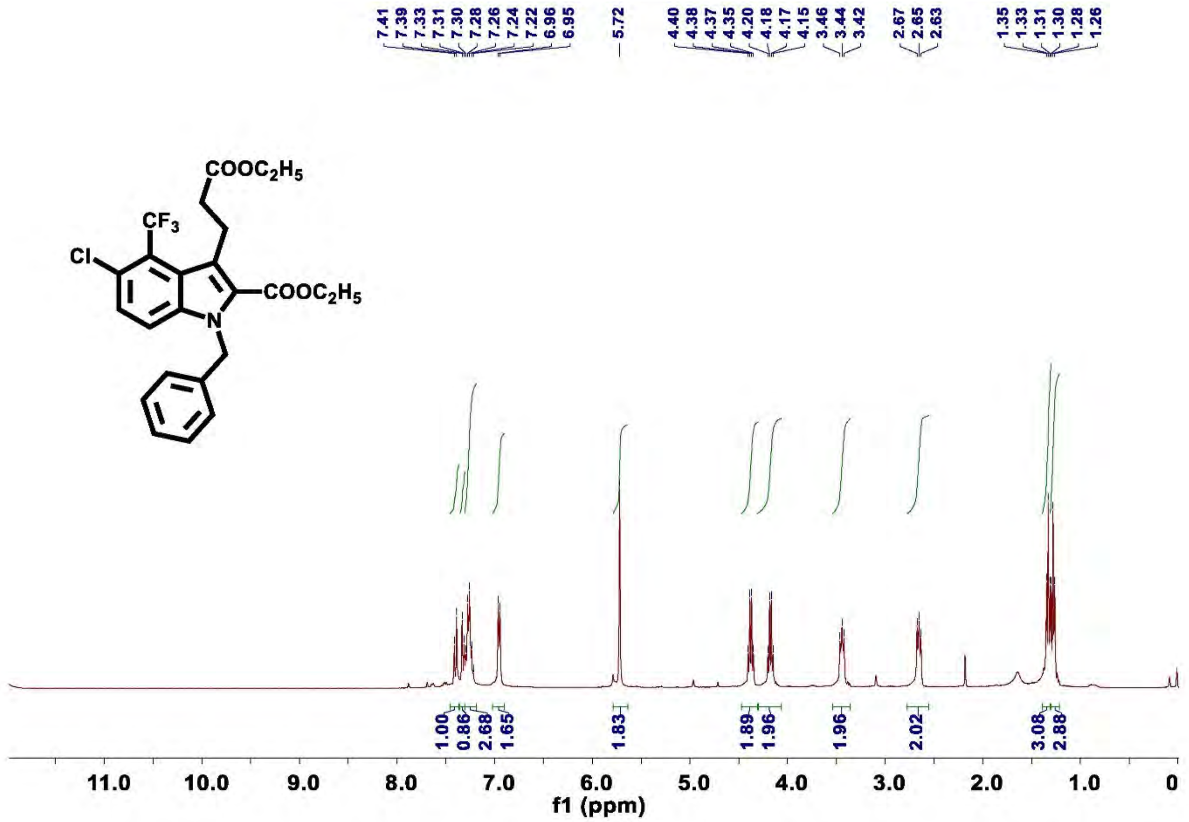

Figure S62. $^{1}H$ NMR of compound 10c

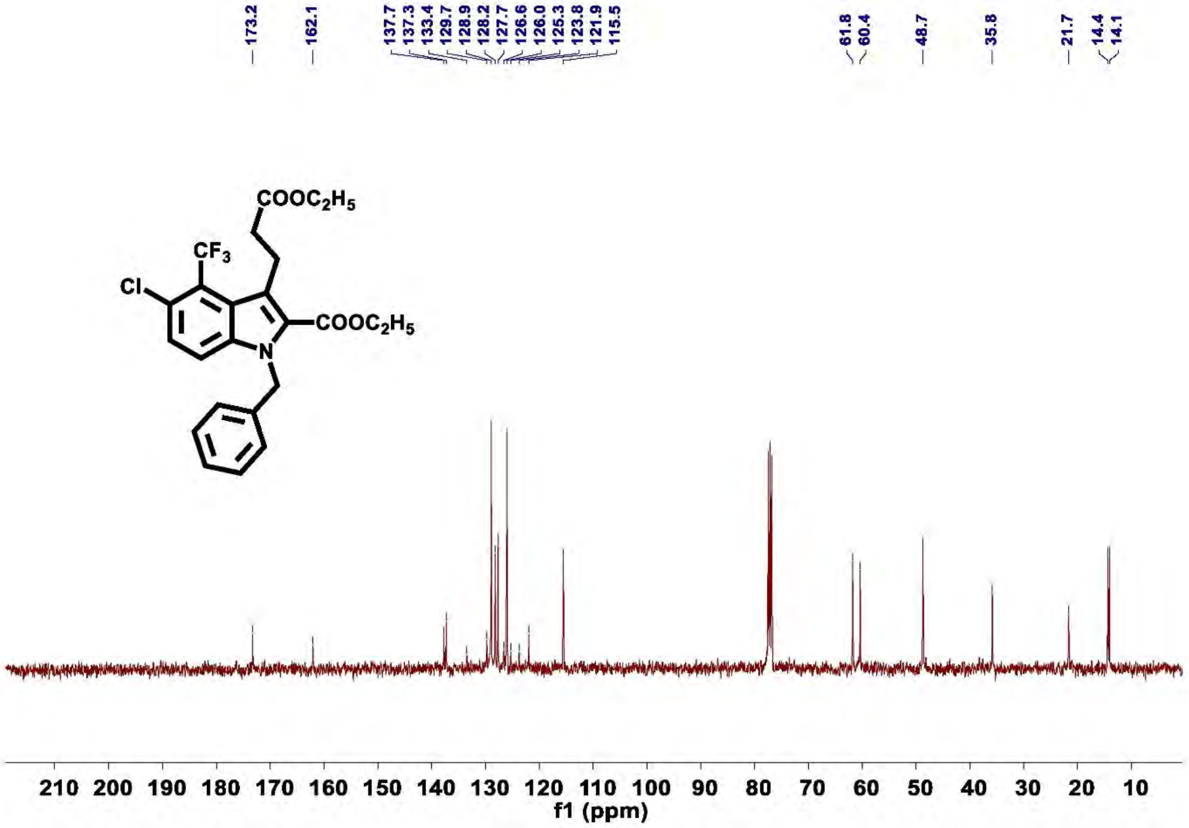

Figure S63. $^{13}C$ NMR of compound 10c

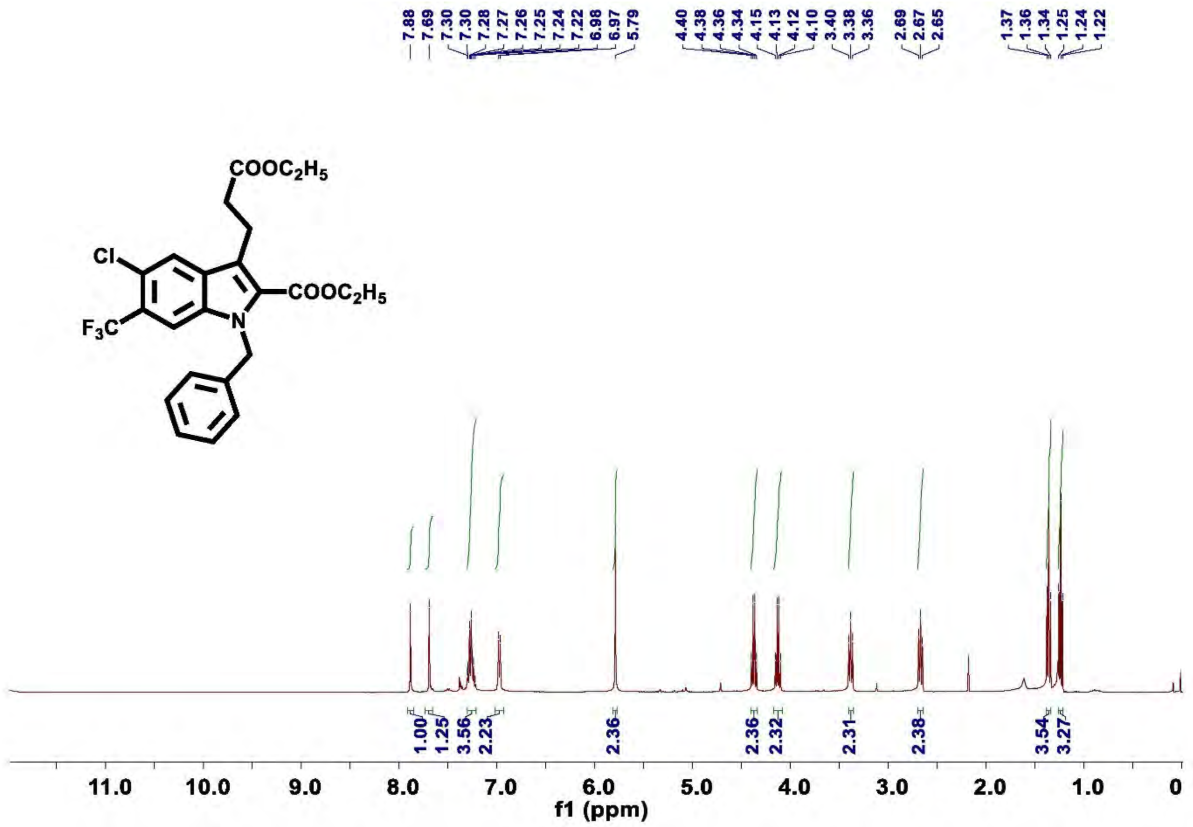

**Figure S64.** $^{1}$**H NMR of compound 10c'**

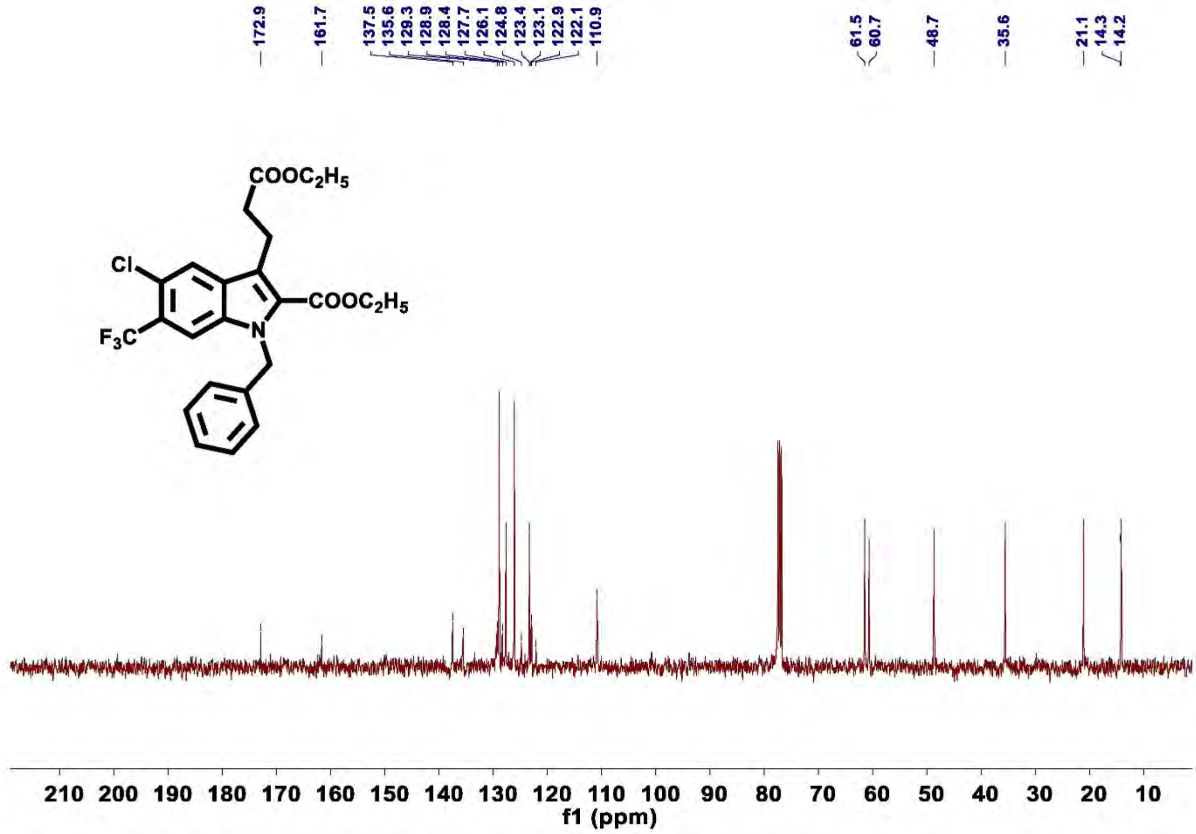

**Figure S65.** $^{13}$**C NMR of compound 10c'**

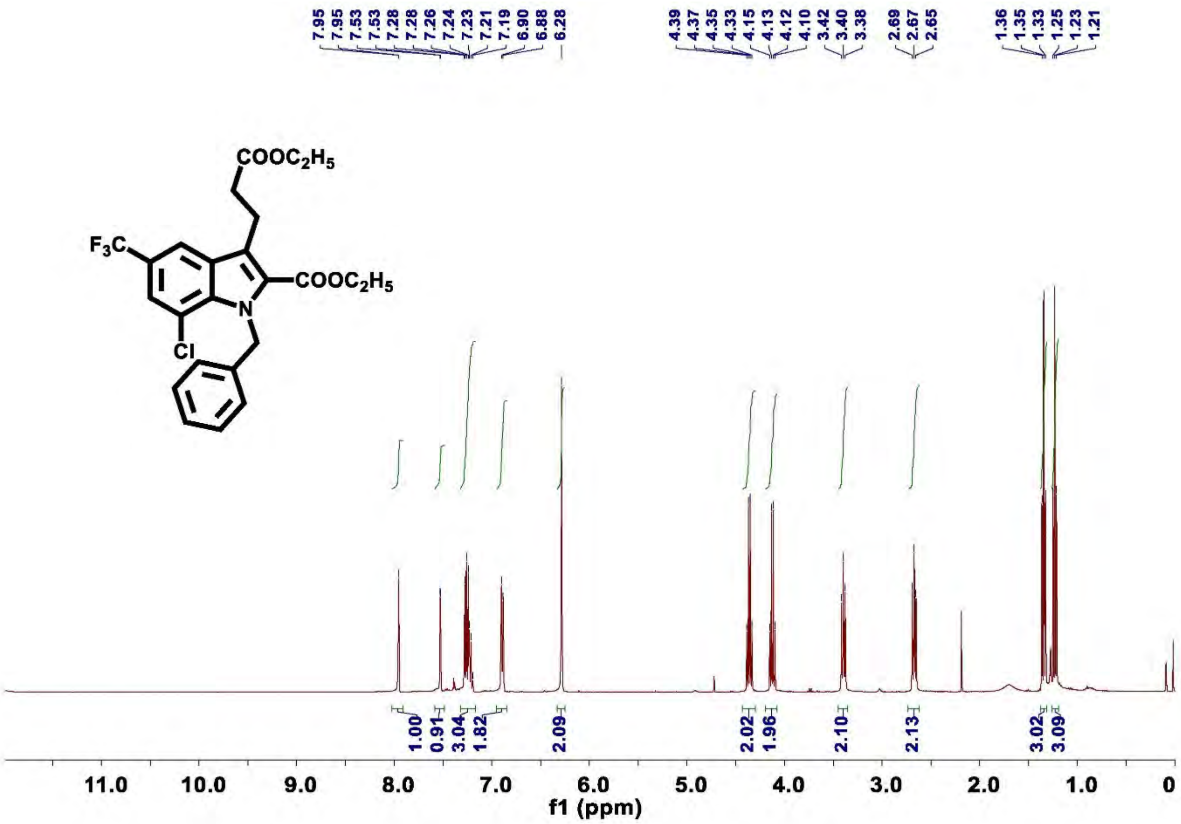

**Figure S66.** $^{1}$**H NMR of compound 10d**

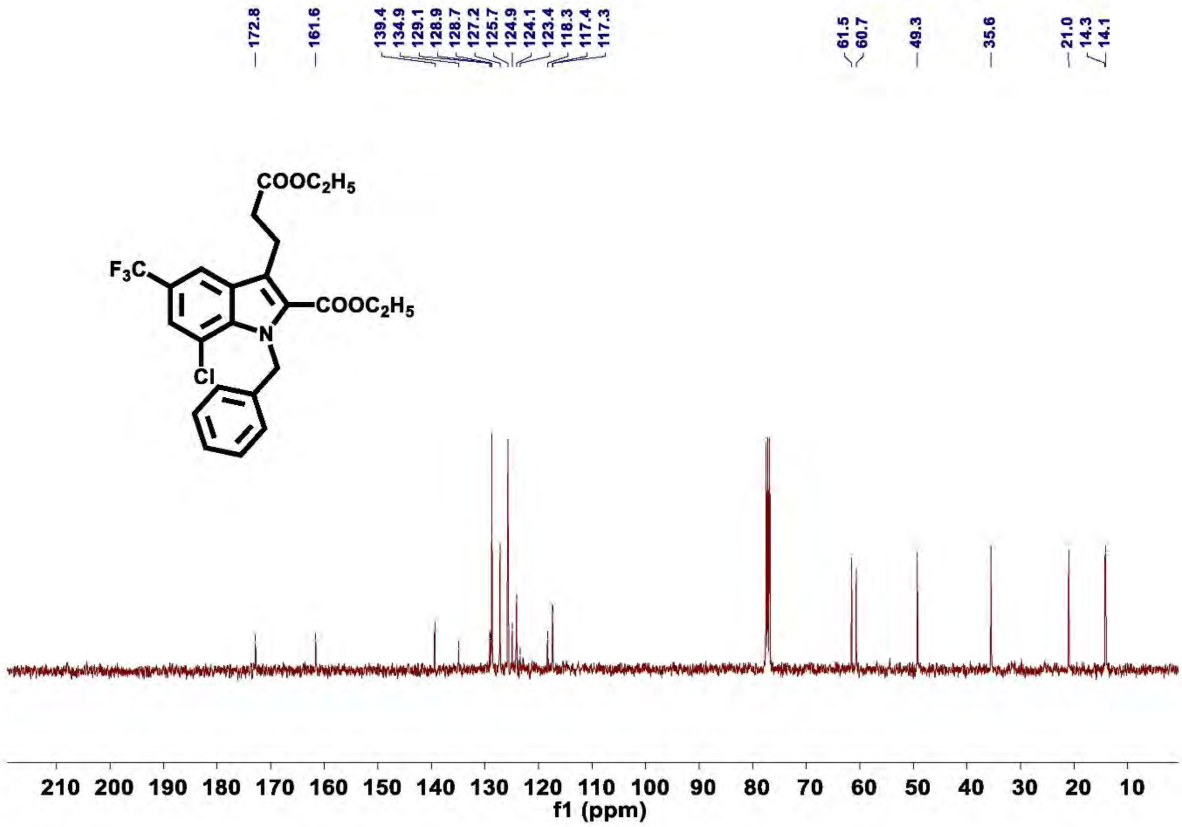

**Figure S67.** $^{13}$**C NMR of compound 10d**

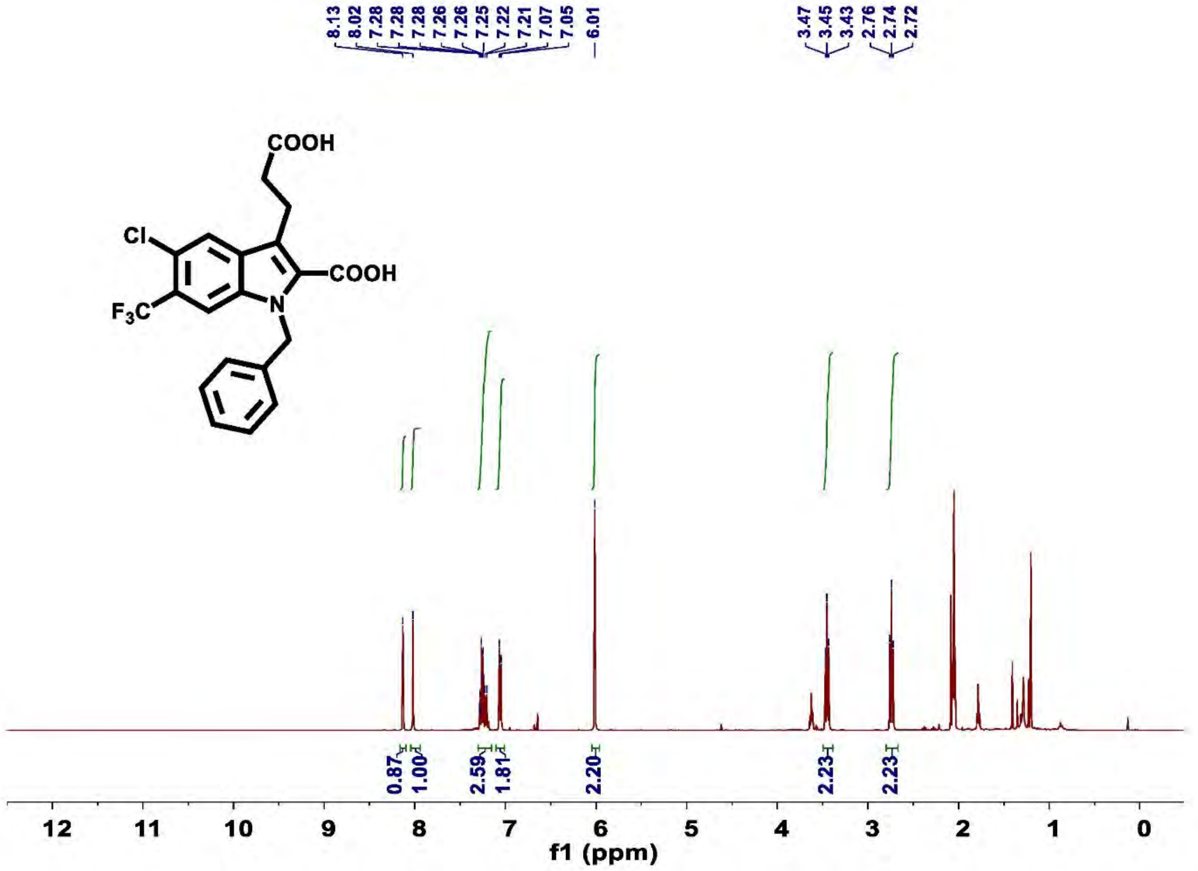

**Figure S68. $^1$H NMR of compound 11c'**

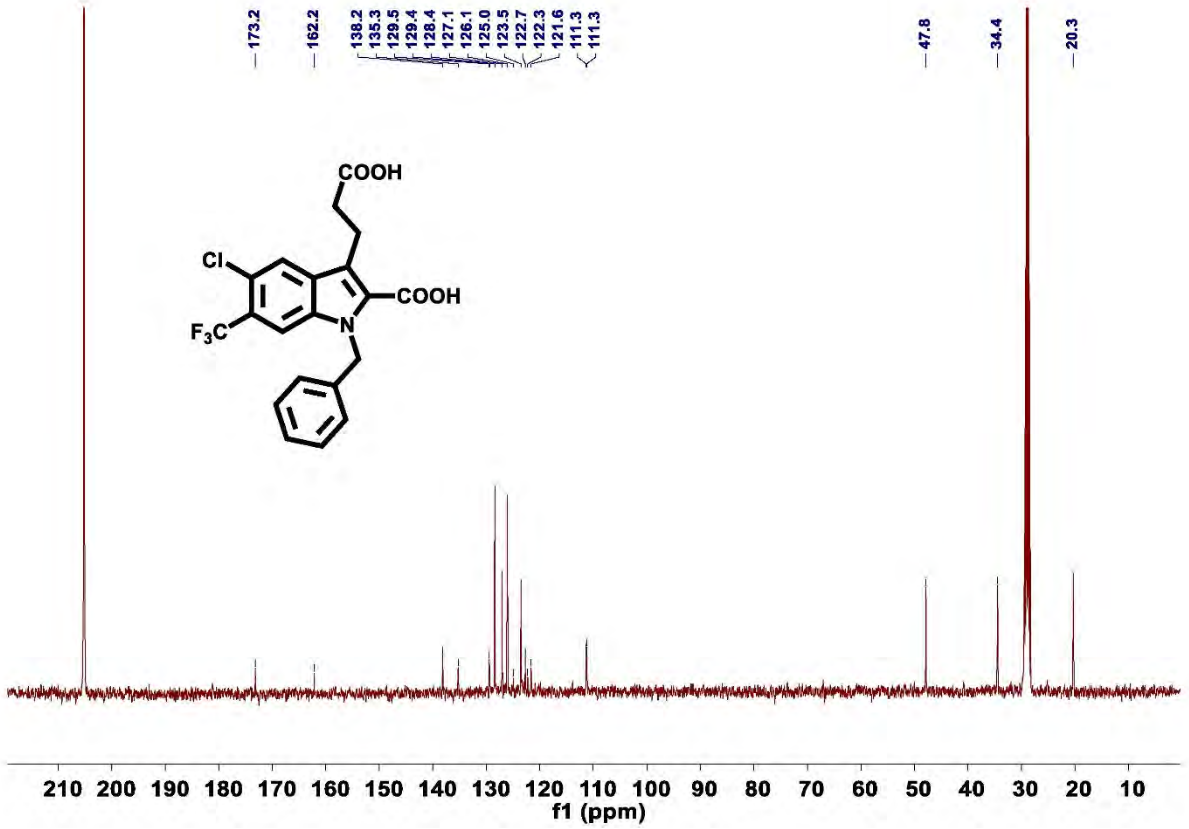

**Figure S69.** $^{13}$**C NMR of compound 11c'**

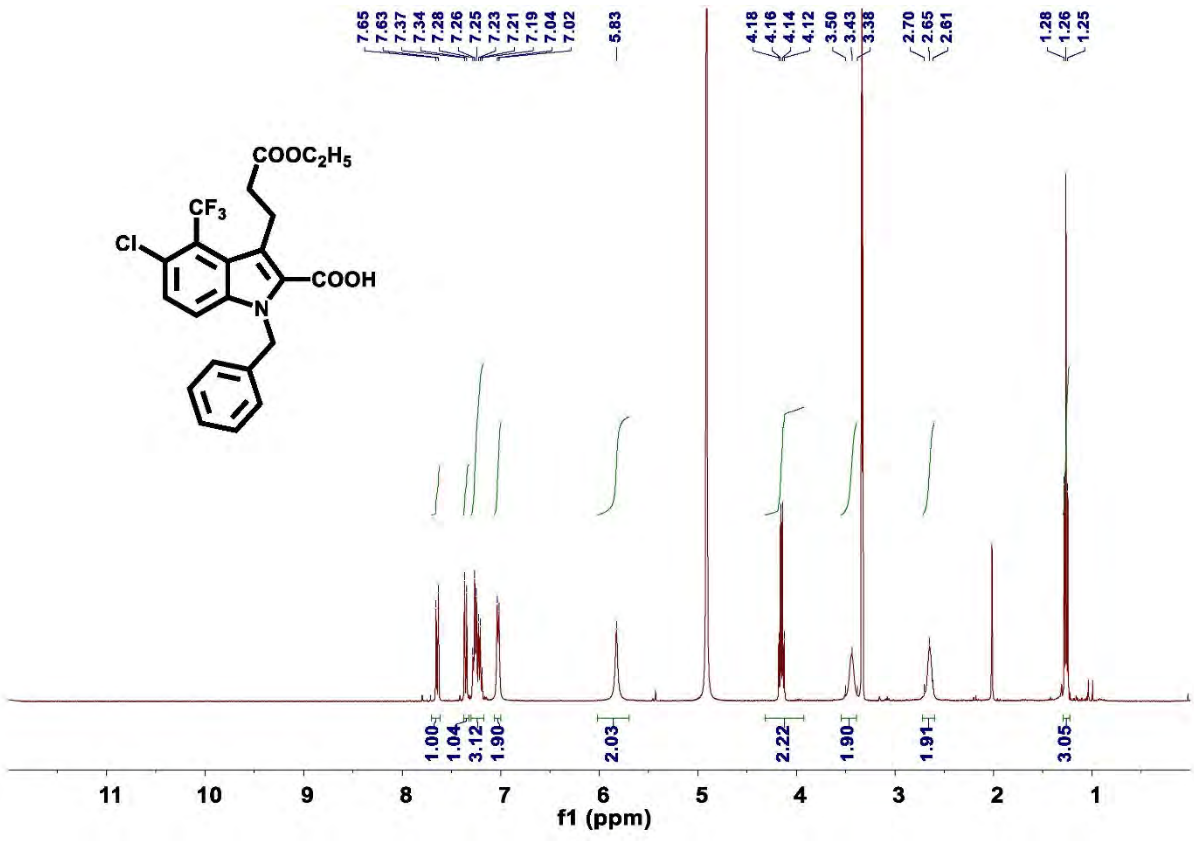

**Figure S70.** $^{1}$**H NMR of compound 12c**

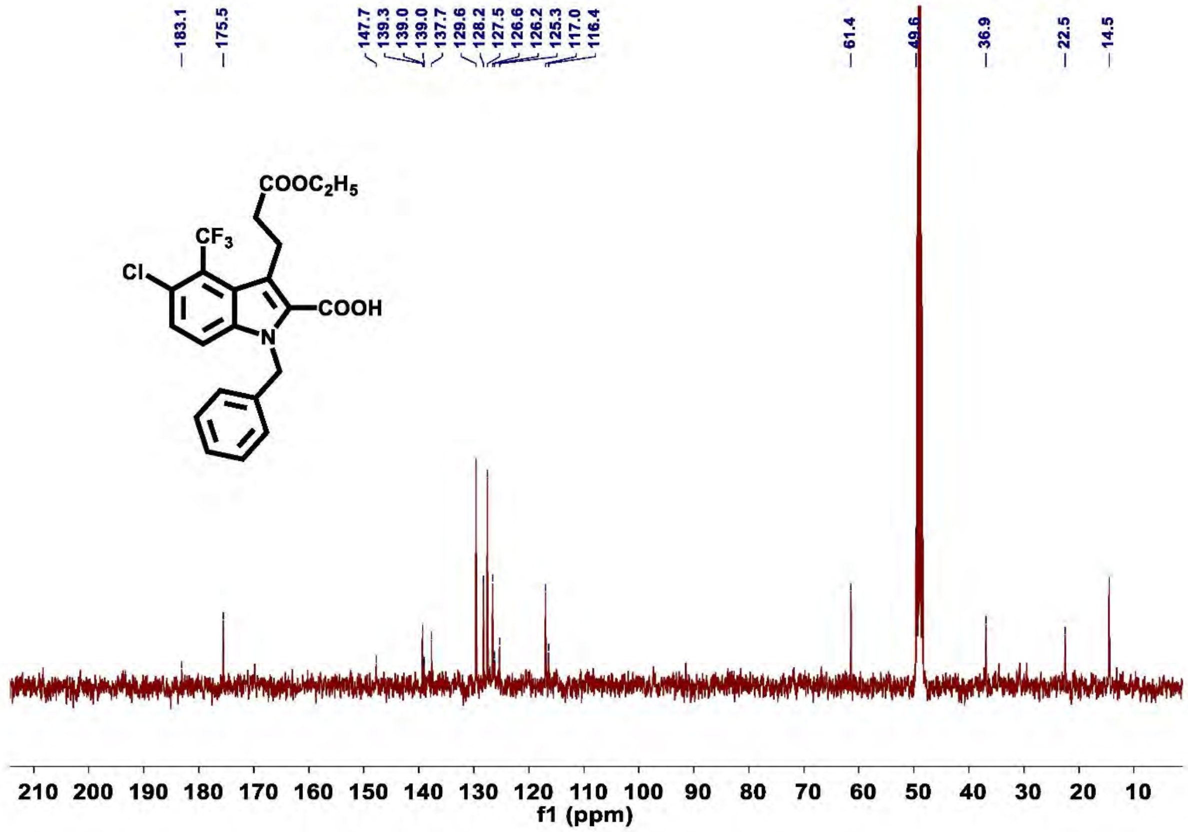

Figure S71. $^{13}$C NMR of compound 12c

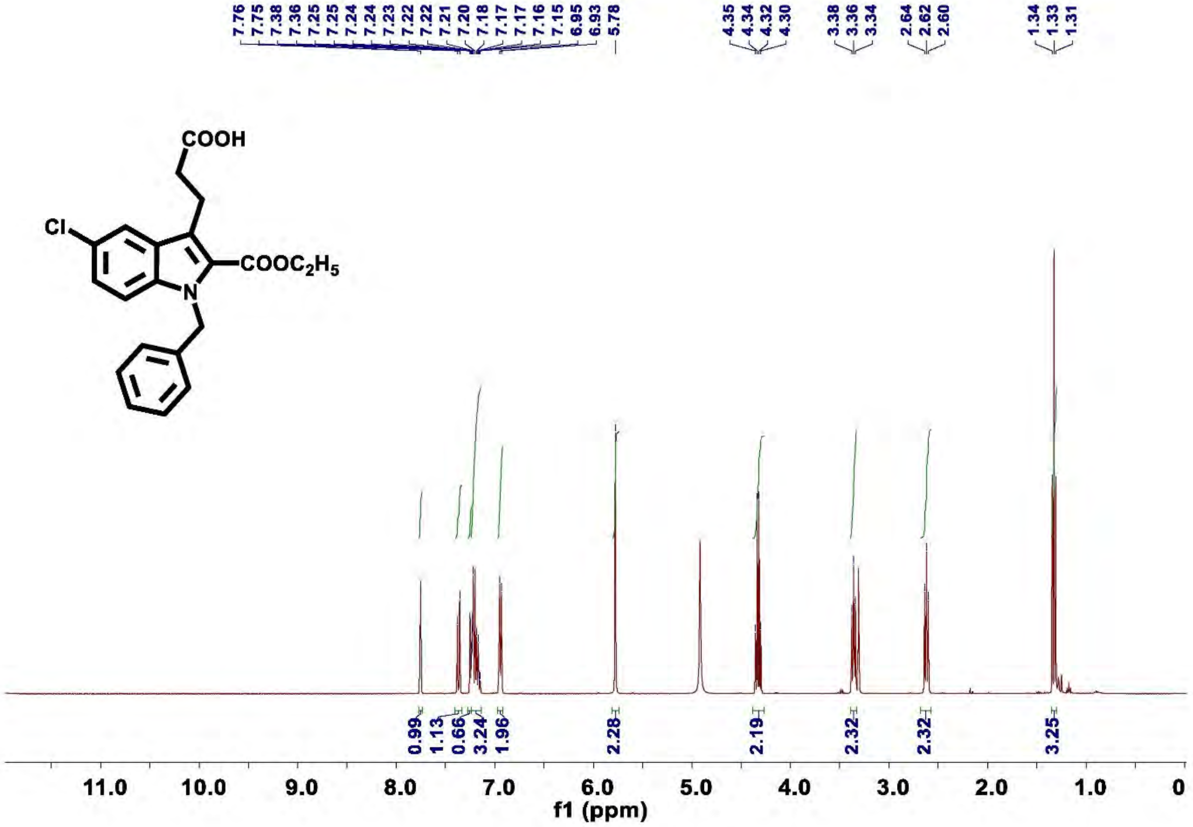

Figure S72. $^{1}$H NMR of compound 13a

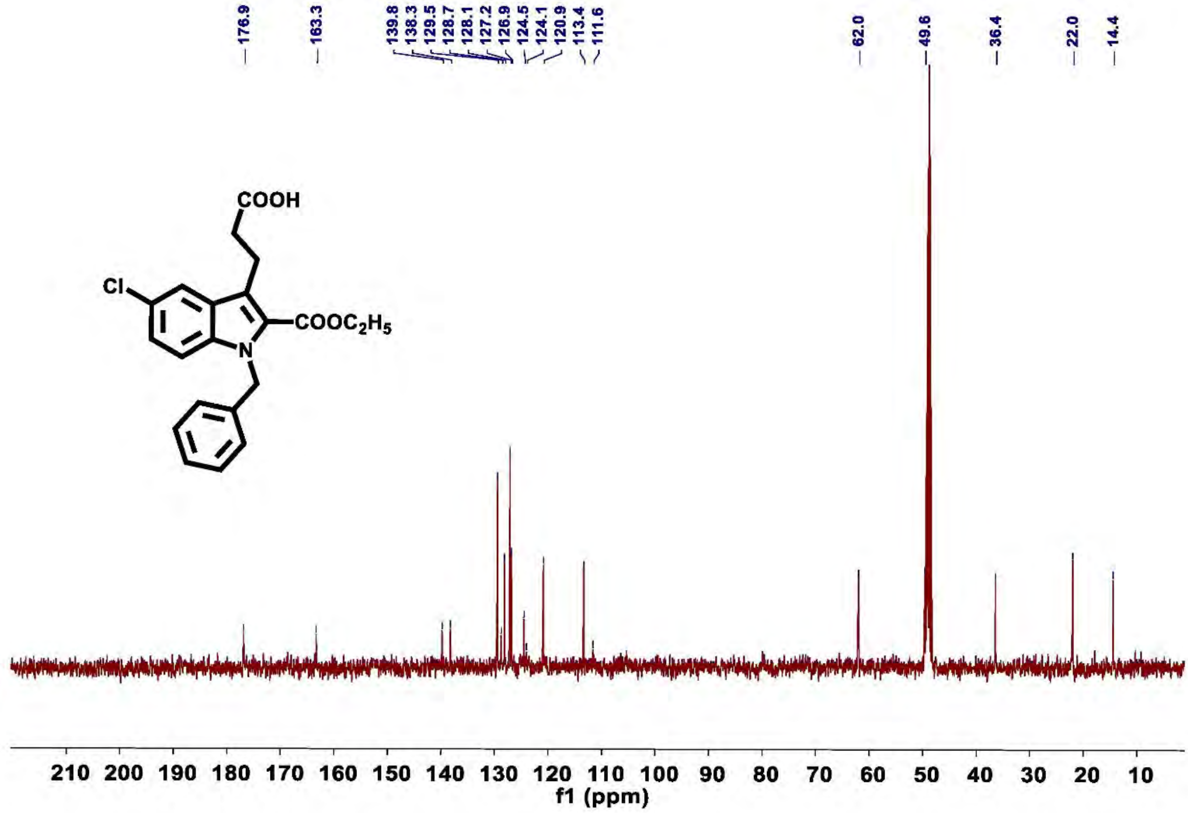

**Figure S73.** $^{13}$C NMR of compound 13a

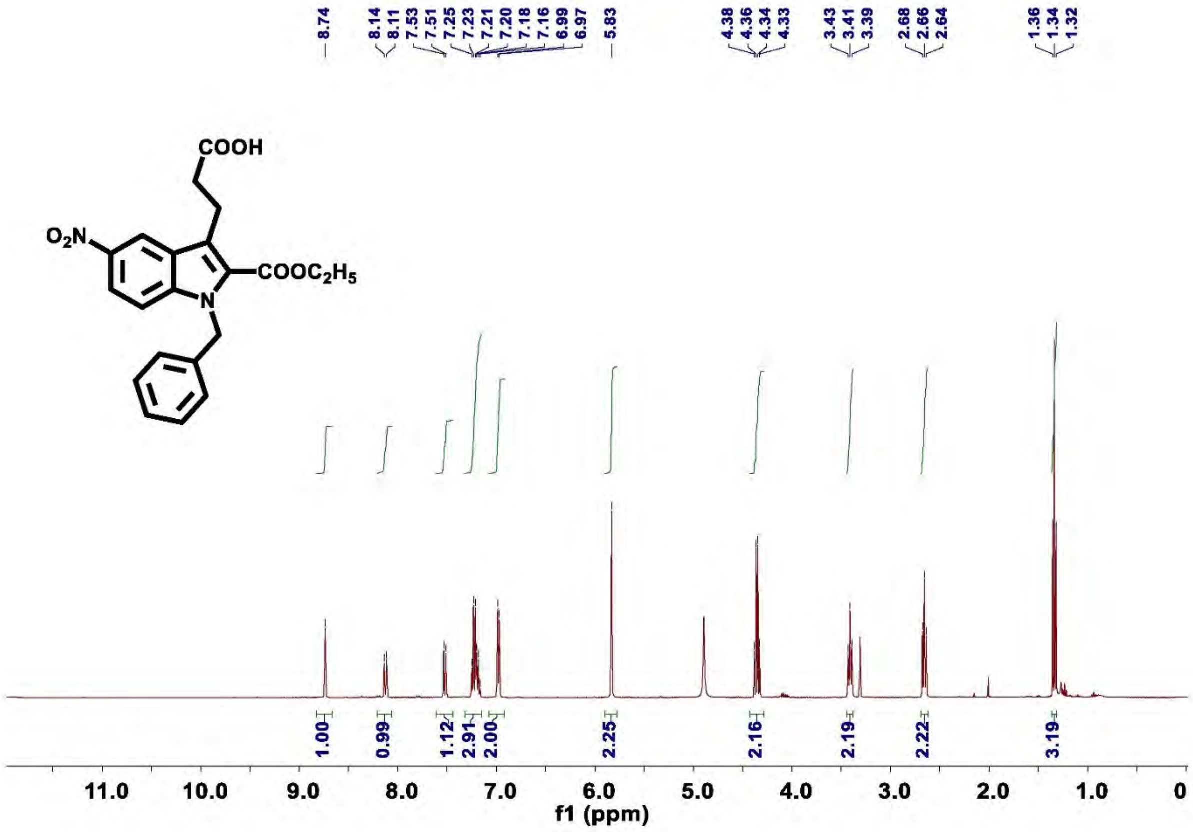

**Figure S74.** $^{1}$H NMR of compound 13b

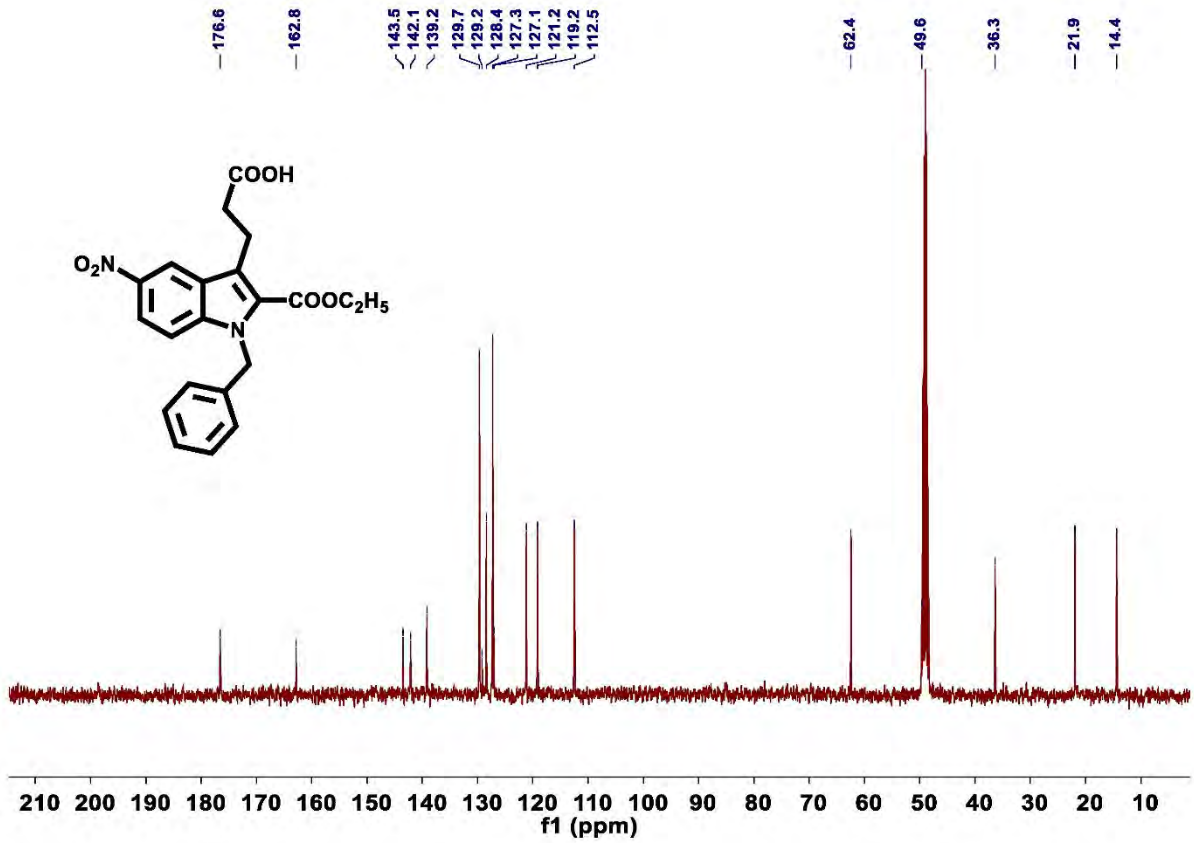

Figure S75. $^{13}C$ NMR of compound 13b

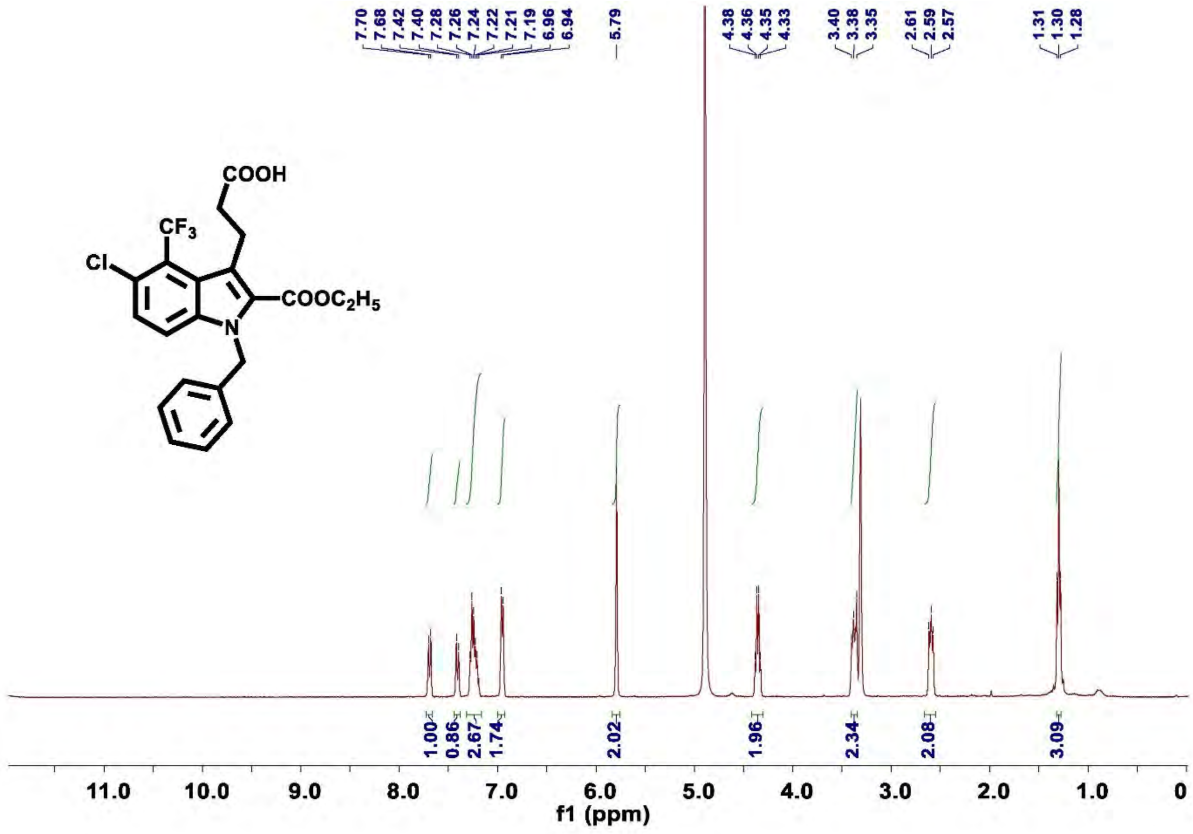

Figure S76. $^{1}H$ NMR of compound 13c

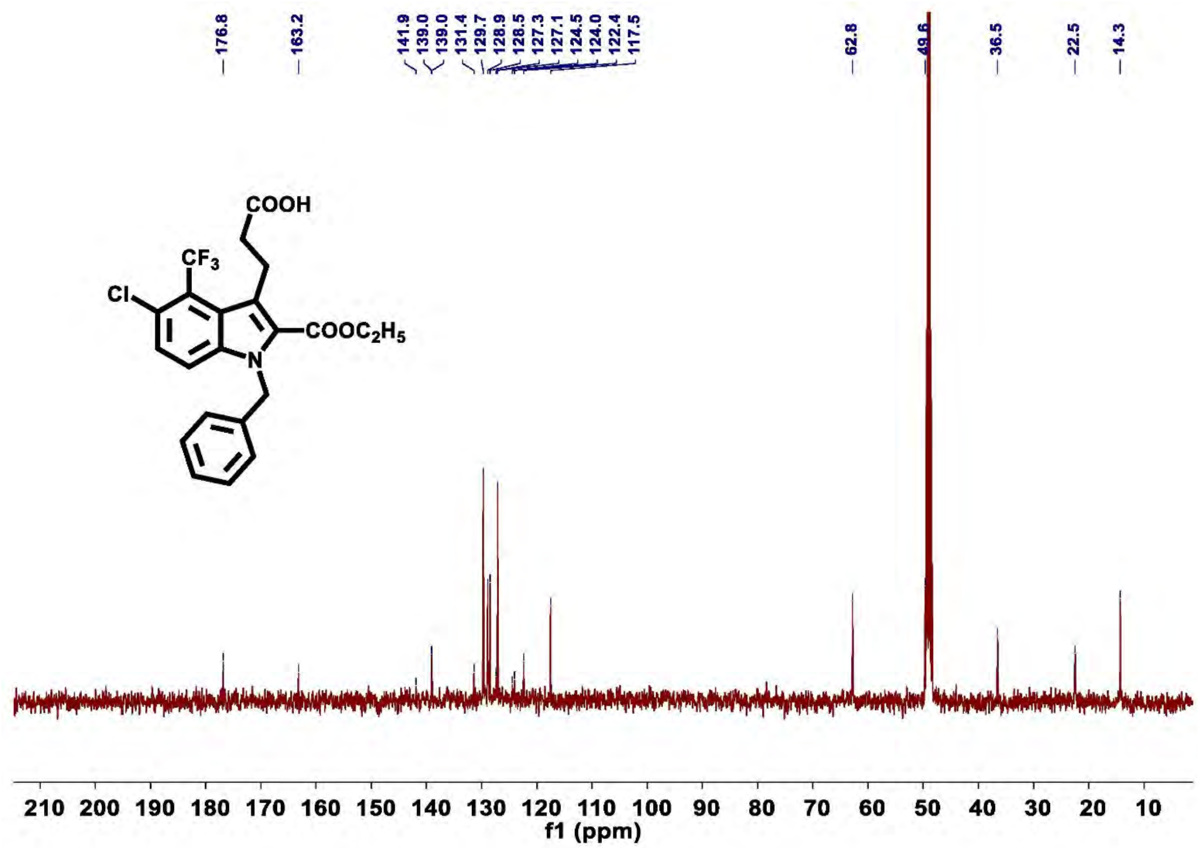

Figure S77. $^{13}$C NMR of compound 13c

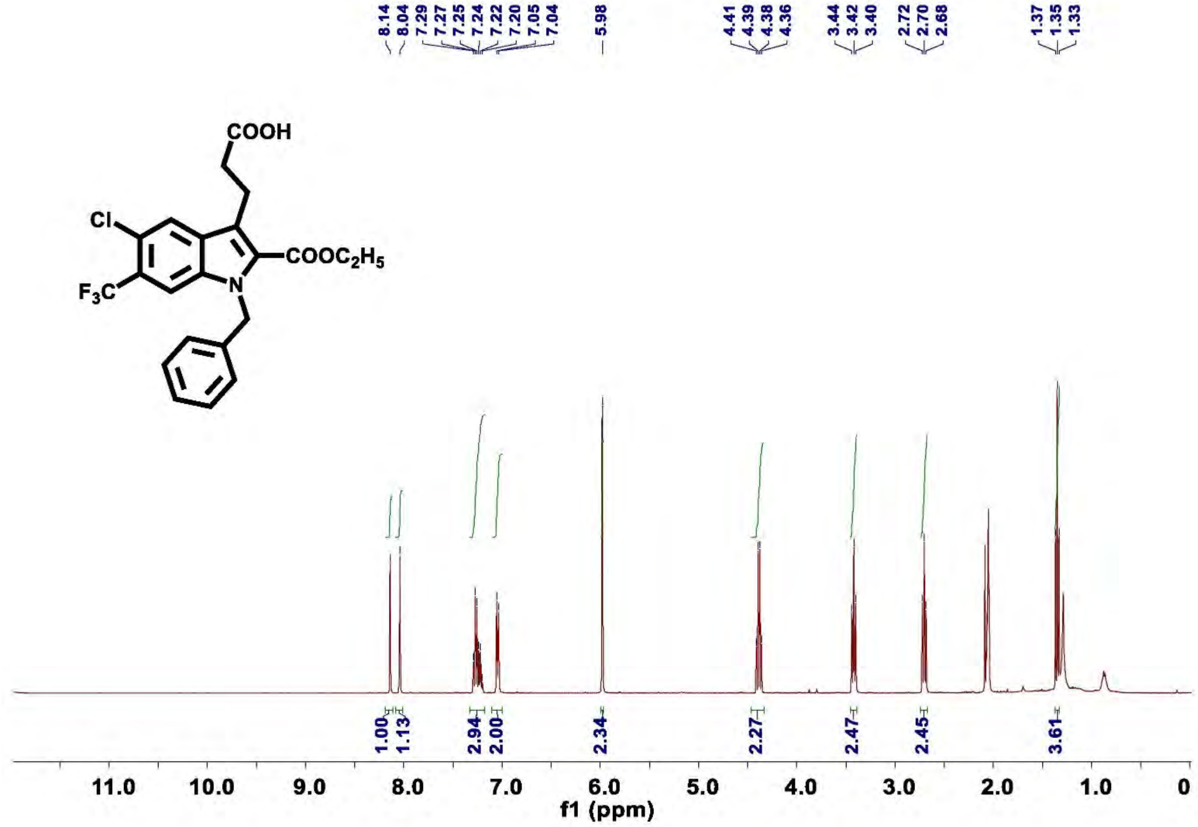

Figure S78. $^{1}$H NMR of compound 13c'

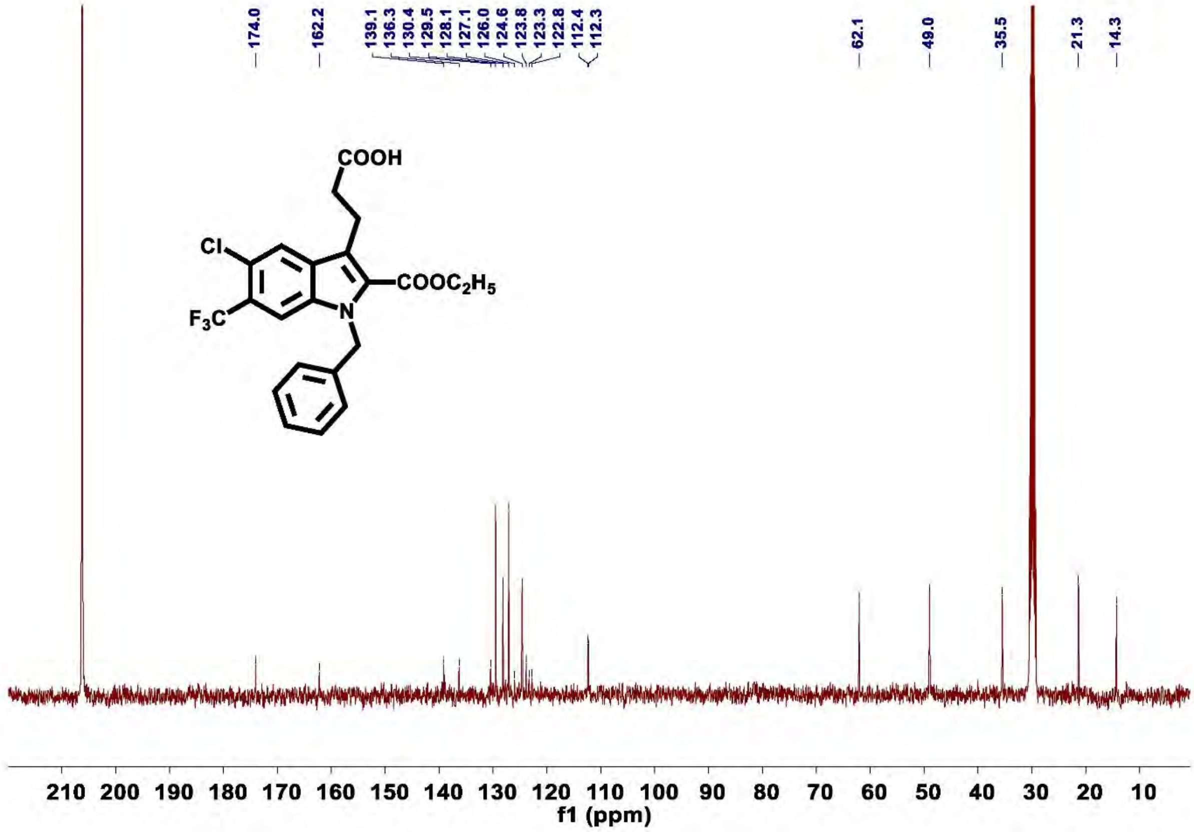

Figure S79. $^{13}$C NMR of compound 13c'

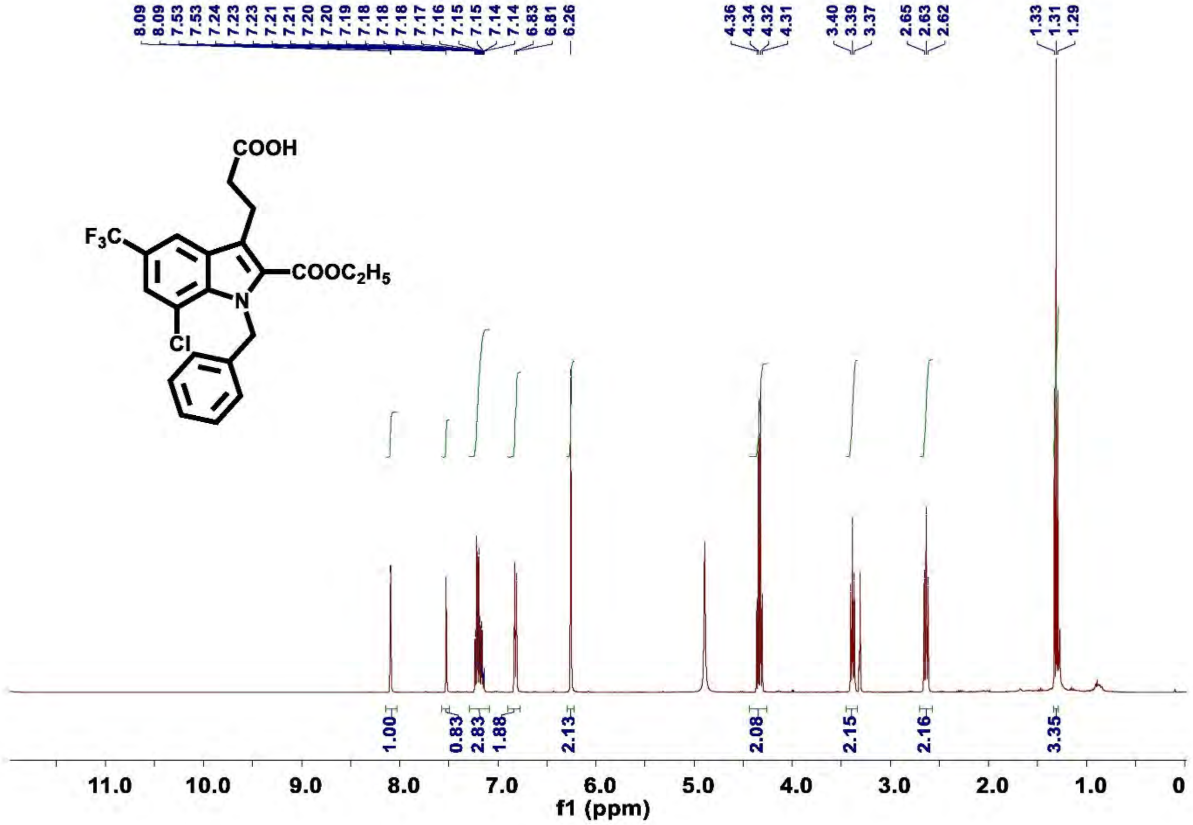

Figure S80. $^{1}$H NMR of compound 13d

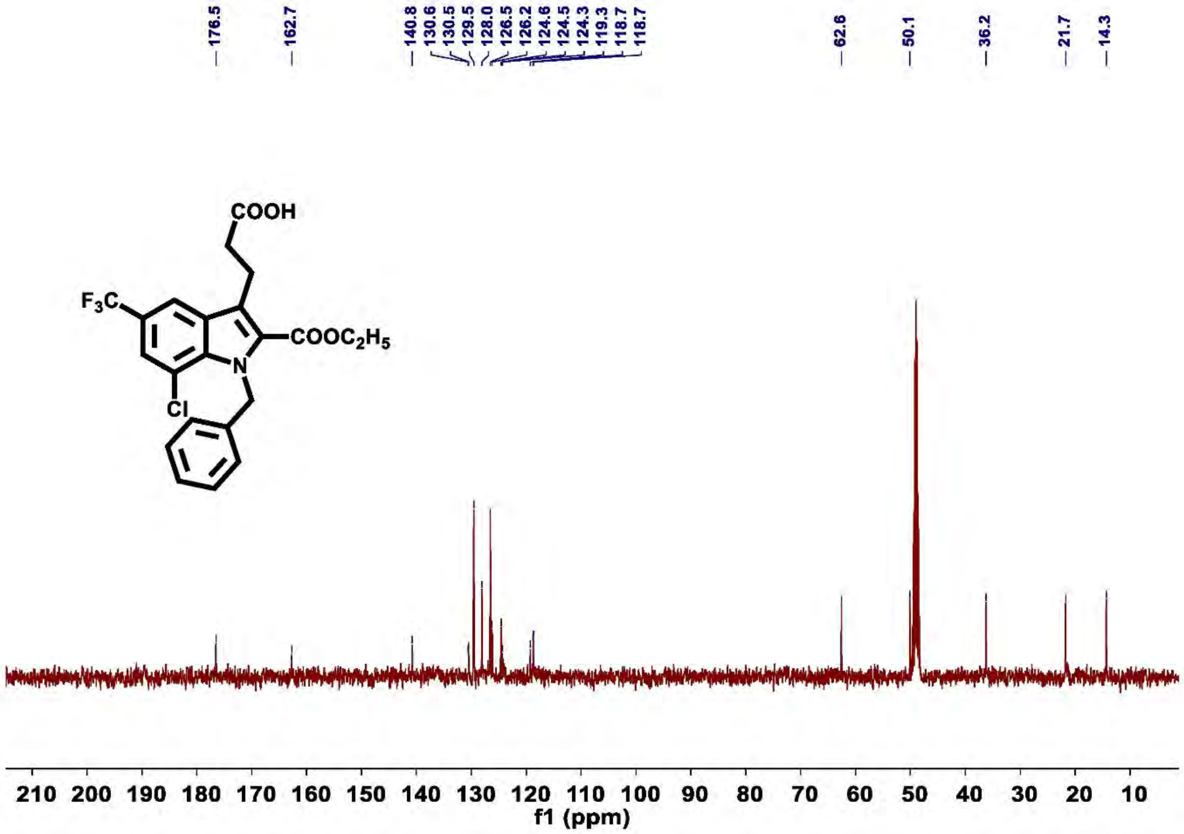

**Figure S81.** $^{13}$C NMR of compound 13d**

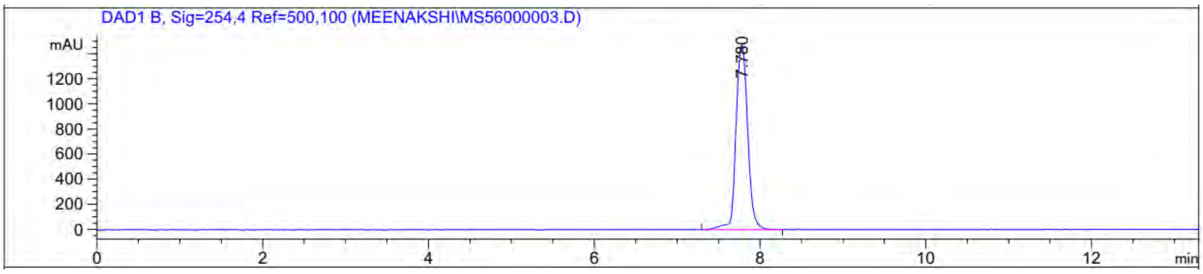

Signal 2: DAD1 B, Sig=254,4 Ref=500,100

| Peak # | RetTime [min] | Type | Width [min] | Area [mAU*s] | Height [mAU] | Area % |
|---|---|---|---|---|---|---|
| 1 | 7.780 | BV | 0.1569 | 1.45001e4 | 1472.21118 | 100.0000 |

| | | | | | |
|---|---|---|---|---|---|
| Totals : | | | | 1.45001e4 | 1472.21118 |

**Figure S82**. HPLC chromatograph of purified compound **13c**; $t_R$ ~7.78 min, Purity: 99.80%, λ = 254 nm.

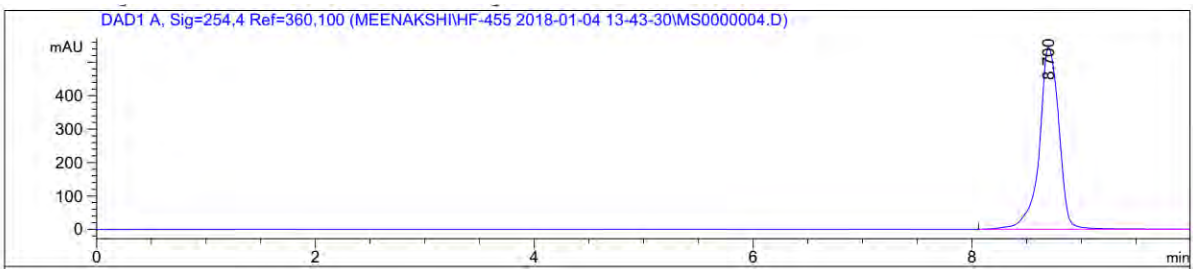

Signal 1: DAD1 A, Sig=254,4 Ref=360,100

| Peak # | RetTime [min] | Type | Width [min] | Area [mAU*s] | Height [mAU] | Area % |
|---|---|---|---|---|---|---|
| 1 | 8.700 | BBA | 0.1957 | 6805.29883 | 544.96045 | 100.0000 |
| Totals : | | | | 6805.29883 | 544.96045 | |

**Figure S83**. HPLC chromatograph of purified compound **13d**; $t_R$ ~8.7 min, Purity: 100%, λ = 254 nm.

**Table S1**

| Molecule name | Name used in: Ilic S. et al Scientific Reports 2016 | ZincID | Vendor | Molecular structure |
|---|---|---|---|---|
| A1 | 1 | ZINC4218923 | Enamine | 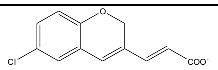 |
| A2 | 2 | ZINC4218626 | Enamine | 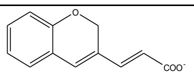 |
| A3 | 3 | ZINC00103556 | ChemBridge Economical | 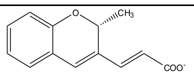 |
| A4 | | ZINC59598 | Princeton BioMolecular Research | 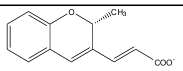 |
| A5 | 5 | ZINC64777050 | Enamine | 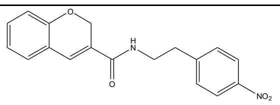 |
| A6 | 6 | ZINC68717 | Vitas-M Laboratory | 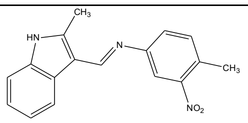 |
| A7 | 7 | ZINC59598 | Vitas-M Laboratory | 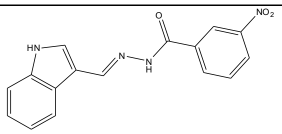 |
| A8 | 8 | ZINC94399 | Vitas-M Laboratory | 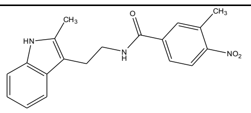 |
| A9 | 9 | ZINC77364 | Vitas-M Laboratory | 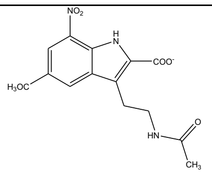 |
| A10 | 10 | ZINC152215 | eMolecules | 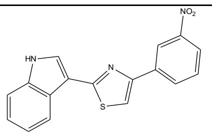 |
| A11 | | ZINC156165 | Sigma | 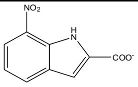 |
| A12 | 12 | ZINC23894 | Sigma | 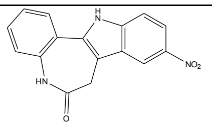 |
| A13 | 13 | ZINC113142 | Maybridge | 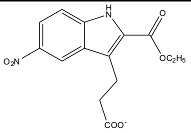 |
| A14 | 14 | ZINC4763 | Maybridge | 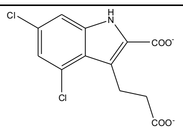 |

| | | | | |
|---|---|---|---|---|
| A15 | 15 | ZINC171585 | Maybridge | 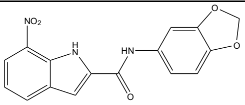 |
| A16 | 16 | ZINC171580 | Maybridge | 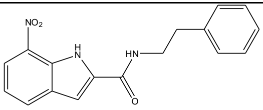 |
| A17 | 17 | ZINC156165 | Maybridge | 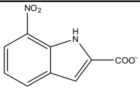 |
| A18 | 18 | ZINC158651 | Maybridge | 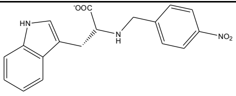 |

Fig2c_data

| Molecule symbol | Primase activity (%) | Gyrase (ATPase activity, %) | Gyrase (supercoiling activity, %) |
|---|---|---|---|
| **No molecule (C+)** | A | A | A |
| **A** | A | A | 97.6 |
| **B** | A | 92 | 80.3 |
| **C** | A | A | 71.9 |
| **D** | A | A | 95.9 |
| **E** | A | A | 98.3 |
| **F** | A | A | 76.6 |
| **G** | 98.9 | 86.7 | 92 |
| **H** | A | 97.1 | 88.4 |
| **I** | 75.3 | 99.6 | 96.8 |
| **J** | 65.9 | 64.3 | 10.9 |
| **K** | A | A | 92.2 |
| **L** | 88.7 | 48.6 | 3.2 |
| **M** | I | 44.5 | 0.9 |
| **N** | 84.8 | 77.3 | 89 |
| **O** | 64.6 | 42.2 | 1.7 |
| **P** | 93.1 | 89.2 | 85.2 |
| **Q** | I | 27.8 | 0.3 |
| **R** | A | 99.5 | 88.6 |
| **A = Full activity (no observed inhibition)** | | | |
| **I = Full inhibition (no activity)** | | | |

Fig3c_data

| Molecule symbol | Primase activity (%) | Gyrase (ATPase activity, %) | Gyrase (supercoiling activity, %) |
|---|---|---|---|
| **3a** | A | 101.0 | 69.6 |
| **3b** | A | 84.4 | 31.5 |
| **3c** | A | 79.4 | 36.8 |
| **3c'** | A | 56.8 | 0.3 |
| **3d** | A | 66.8 | 28.9 |
| **3e** | A | 82.5 | 75.7 |
| **3f** | A | 86.7 | 44.4 |
| **3g** | A | 102.1 | 84.7 |
| **4a** | 83.7 | 94.0 | 82.0 |
| **5a** | A | 95.2 | 74.1 |
| **6a** | A | 85.8 | 84.9 |
| **6b** | A | 74.4 | 79.6 |
| **6c** | A | 59.9 | 78.2 |
| **6c'** | 78.2 | 98.4 | 79.2 |
| **6d** | A | 60.2 | 88.1 |
| **6e** | A | 67.6 | 84.0 |
| **6f** | 95.1 | 71.2 | 69.5 |
| **6g** | A | 109.6 | 84.3 |
| **7a** | A | 5.2 | 0.8 |
| **8b** | 65.3 | 79.1 | 91.2 |
| **8c** | I | 57.8 | 2.8 |
| **8c'** | A | 116.7 | 93.2 |
| **8d** | 26.7 | 76.3 | 59.7 |
| **8f** | I | 29.8 | 0.2 |
| **9a** | A | 68.3 | 85.2 |
| **9b** | 91.1 | 119.2 | 81.8 |
| **9c** | 80.7 | 70.4 | 79.9 |
| **9c'** | 80.8 | 43.4 | 35.9 |
| **9d** | A | 78.6 | 96.2 |
| **9e** | 63.5 | 52.3 | 68.5 |
| **9f** | A | 84.5 | 34.4 |
| **10a** | A | 89.9 | 61.1 |
| **10b** | A | 50.2 | 70.8 |
| **10c** | A | 114.5 | 47.5 |
| **10c'** | A | 80.8 | 73.7 |
| **10d** | A | 66.5 | 30.6 |
| **11a** | A | 77.1 | 88.7 |
| **11b** | A | 99.1 | 99.8 |
| **11c'** | 3.7 | 31.5 | 39.5 |
| **11d** | A | 89.4 | 66.5 |
| **12a** | 75.2 | 67.6 | 92.4 |
| **12b** | A | 83.5 | 95.2 |
| **12c** | 35.7 | 13.5 | 0.7 |
| **12c'** | 44.5 | 51.7 | 65.8 |

| | | | |
|---|---|---|---|
| **13a** | 5.2 | 34.6 | 2.1 |
| **13b** | 18.2 | 74.1 | 97.4 |
| **13c** | I | 26.3 | 0.0 |
| **13c'** | 83.3 | 42.0 | 0.0 |
| **13d** | I | 34.8 | 0.0 |
| **A = Fully active (no observed inhibition)** | | | |
| **I = Full inhibition (No activity)** | | | |

Molecular Formula String

| Compound | SMILES |
|---|---|
| **3a** | ClC1=CC=C(NC(C(OCC)=O)=C2CCC(OCC)=O)C2=C1 |
| **3b** | O=C(C1=C(CCC(OCC)=O)C2=CC([N+]([O-])=O)=CC=C2N1)OCC |
| **3c** | ClC1=CC=C(NC(C(OCC)=O)=C2CCC(OCC)=O)C2=C1C(F)(F)F |
| **3c’** | ClC1=C(C(F)(F)F)C=C(NC(C(OCC)=O)=C2CCC(OCC)=O)C2=C1 |
| **3d** | ClC1=C(NC(C(OCC)=O)=C2CCC(OCC)=O)C2=CC(C(F)(F)F)=C1 |
| **3e** | ClC1=C2C(NC(C(OCC)=O)=C2CCC(OCC)=O)=C(Cl)C=C1[N+]([O-])=O |
| **3f** | O=C(C1=C(CCC(OCC)=O)C2=C(C(F)(F)F)C=C(C(F)(F)F)C=C2N1)OCC |
| **3g** | CC1=CC=C(NC(C(OCC)=O)=C2CCC(OCC)=O)C2=C1 |
| **4a** | ClC1=CC=C(NC(CO)=C2CCCO)C2=C1 |
| **5a** | ClC1=CC=C(NC(C=O)=C2CCC=O)C2=C1 |
| **6a** | ClC1=CC=C(NC(C(O)=O)=C2CCC(O)=O)C2=C1 |
| **6b** | O=C(C1=C(CCC(O)=O)C2=CC([N+]([O-])=O)=CC=C2N1)O |
| **6c** | ClC1=CC=C(NC(C(O)=O)=C2CCC(O)=O)C2=C1C(F)(F)F |
| **6c'** | ClC1=C(C(F)(F)F)C=C(NC(C(O)=O)=C2CCC(O)=O)C2=C1 |
| **6d** | ClC1=C(NC(C(O)=O)=C2CCC(O)=O)C2=CC(C(F)(F)F)=C1 |
| **6e** | ClC1=C2C(NC(C(O)=O)=C2CCC(O)=O)=C(Cl)C=C1[N+]([O-])=O |
| **6f** | O=C(C1=C(CCC(O)=O)C2=C(C(F)(F)F)C=C(C(F)(F)F)C=C2N1)O |
| **6g** | CC1=CC=C(NC(C(O)=O)=C2CCC(O)=O)C2=C1 |
| **7a** | ClC1=CC=C(NC(C(NC2=CC=CC=C2)=O)=C3CCC(NC4=CC=CC=C4)=O)C3=C1 |
| **8b** | O=C(C1=C(CCC(OCC)=O)C2=CC([N+]([O-])=O)=CC=C2N1)O |
| **8c** | ClC1=CC=C(NC(C(O)=O)=C2CCC(OCC)=O)C2=C1C(F)(F)F |
| **8c'** | ClC1=C(C(F)(F)F)C=C(NC(C(O)=O)=C2CCC(OCC)=O)C2=C1 |
| **8d** | ClC1=C(NC(C(O)=O)=C2CCC(OCC)=O)C2=CC(C(F)(F)F)=C1 |
| **8f** | O=C(C1=C(CCC(OCC)=O)C2=C(C(F)(F)F)C=C(C(F)(F)F)C=C2N1)O |
| **9a** | ClC1=CC=C(NC(C(OCC)=O)=C2CCC(OCC)=O)C2=C1 |
| **9b** | O=C(C1=C(CCC(O)=O)C2=CC([N+]([O-])=O)=CC=C2N1)OCC |
| **9c** | ClC1=CC=C(NC(C(OCC)=O)=C2CCC(O)=O)C2=C1C(F)(F)F |
| **9c'** | ClC1=C(C(F)(F)F)C=C(NC(C(OCC)=O)=C2CCC(OCC)=O)C2=C1 |
| **9d** | ClC1=C(NC(C(OCC)=O)=C2CCC(OCC)=O)C2=CC(C(F)(F)F)=C1 |
| **9e** | ClC1=C2C(NC(C(OCC)=O)=C2CCC(O)=O)=C(Cl)C=C1[N+]([O-])=O |
| **9f** | FC(C1=CC(C(F)(F)F)=C2C(NC(C(OCC)=O)=C2CCC(O)=O)=C1)(F)F |
| **10a** | ClC1=CC=C(N(CC2=CC=CC=C2)C(C(OCC)=O)=C3CCC(OCC)=O)C3=C1 |

| | |
|---|---|
| **10b** | O=[N+](C1=CC=C(N(CC2=CC=CC=C2)C(C(OCC)=O)=C3CCC(OCC)=O)C3=C1)[O-] |
| **10c** | ClC1=CC=C(N(CC2=CC=CC=C2)C(C(OCC)=O)=C3CCC(OCC)=O)C3=C1C(F)(F)F |
| **10c'** | ClC1=C(C(F)(F)F)C=C(N(CC2=CC=CC=C2)C(C(OCC)=O)=C3CCC(OCC)=O)C3=C1 |
| **10d** | ClC1=C(N(CC2=CC=CC=C2)C(C(OCC)=O)=C3CCC(OCC)=O)C3=CC(C(F)(F)F)=C1 |
| **11a** | ClC1=CC=C(N(CC2=CC=CC=C2)C(C(O)=O)=C3CCC(O)=O)C3=C1 |
| **11b** | O=C(C1=C(CCC(O)=O)C2=CC([N+]([O-])=O)=CC=C2N1CC3=CC=CC=C3)O |
| **11c'** | ClC1=C(C(F)(F)F)C=C(N(CC2=CC=CC=C2)C(C(O)=O)=C3CCC(O)=O)C3=C1 |
| **11d** | ClC1=C(N(CC2=CC=CC=C2)C(C(O)=O)=C3CCC(O)=O)C3=CC(C(F)(F)F)=C1 |
| **12a** | ClC1=CC=C(N(CC2=CC=CC=C2)C(C(O)=O)=C3CCC(OCC)=O)C3=C1 |
| **12b** | O=[N+](C1=CC=C(N(CC2=CC=CC=C2)C(C(O)=O)=C3CCC(OCC)=O)C3=C1)[O-] |
| **12c** | ClC1=CC=C(N(CC2=CC=CC=C2)C(C(O)=O)=C3CCC(OCC)=O)C3=C1C(F)(F)F |
| **12c'** | ClC1=C(C(F)(F)F)C=C(N(CC2=CC=CC=C2)C(C(O)=O)=C3CCC(OCC)=O)C3=C1 |
| **13a** | ClC1=CC=C(N(CC2=CC=CC=C2)C(C(OCC)=O)=C3CCC(O)=O)C3=C1 |
| **13b** | O=C(C1=C(CCC(O)=O)C2=CC([N+]([O-])=O)=CC=C2N1CC3=CC=CC=C3)OCC |
| **13c** | ClC1=CC=C(N(CC2=CC=CC=C2)C(C(OCC)=O)=C3CCC(O)=O)C3=C1C(F)(F)F |
| **13c'** | ClC1=C(C(F)(F)F)C=C(N(CC2=CC=CC=C2)C(C(OCC)=O)=C3CCC(O)=O)C3=C1 |
| **13d** | ClC1=C(N(CC2=CC=CC=C2)C(C(OCC)=O)=C3CCC(O)=O)C3=CC(C(F)(F)F)=C1 |
